# Systematic Benchmarking of AI-Based Molecular Generation Models for Structure-Based Drug Design

**DOI:** 10.64898/2026.08.14.744939

**Authors:** Himansu Kumar, Zhengxiao Yang, Yankai Yu, Jianguo Wen, Pora Kim, Xiaobo Zhou

## Abstract

Generative artificial intelligence is accelerating molecular design, yet the relative suitability of available models for different targets and stages of preclinical drug discovery remains unclear. Here we benchmarked 12 molecular generation and optimization methods across 176 curated protein–ligand systems spanning diverse therapeutic target classes, with experimentally validated ligands providing reference chemical space. The evaluated methods encompassed pocket-conditioned 3D generation, diffusion and flow-based modeling, autoregressive construction, reference-conditioned optimization and synthesis-aware design. Performance was assessed using operational robustness, chemical validity, uniqueness, molecular and scaffold diversity, quantitative estimate of drug-likeness, synthetic accessibility, docking, physicochemical and ADMET properties, and computational resource requirements. The results revealed architecture-dependent trade off such as receptor-conditioned methods exploited binding-pocket geometry, flow-based approaches enabled efficient sampling, reference-conditioned methods favored analogue generation, and synthesis-aware approaches improved chemical feasibility, but no method consistently optimized all criteria. To address the functional potential of generated molecules, we further developed a state-aware functional classifier (SAFC) that integrates molecular dynamics derived receptor ensembles, ensemble docking and protein ligand interaction graphs. SAFC provided dynamics-aware functional activity rankings for generated molecules that were partly complementary to docking, drug-likeness and synthetic accessibility scores. These findings support hybrid, stage specific deployment of generative models rather than reliance on any single architecture or evaluation metric. This study provides practical guidelines for generative AI based preclinical drug development processes.

## Introduction

AlphaFold^1^ and related advances in protein-structure prediction have transformed structure-based drug discovery by expanding access to structural information across the proteome. However, structural availability alone does not solve the central challenge of converting protein models into chemically synthesizable, drug-like and functionally active small-molecule therapeutics. Conventional virtual screening pipelines typically search pre-existing compounds from resources such as Enamine, ZINC^2, 3^, and ChEMBL^4^ and prioritize candidates using docking programs such as AutoDock^5^, Vina^6^, GLIDE^7^. Although these approaches remain valuable, they are constrained by the coverage of existing chemical libraries and often struggle to simultaneously optimize binding affinity, molecular novelty, drug-likeness, pharmacokinetic properties and synthetic feasibility.

Generative AI has emerged as a complementary strategy for exploring chemical space beyond enumerated compound libraries. Recent models span several molecular representations and sampling paradigms, including SMILES-based generators, graph-based models, pocket-conditioned three-dimensional generators, diffusion and score-based models, flow-based and flow-matching frameworks, autoregressive molecular construction methods, ligand/reference-conditioned optimization models and synthesis-aware reaction-based systems. Diffusion and score-based approaches such as DiffSBDD^8^, TargetDiff^9^, DiffBP^10^, DiffGui^11^, DiffSMol^12^, DecompDiff^13^, NucleusDiff^14^, EQGAT-diff^15^, MiDi^16^, JODO^17^, GEOLDM^18^ and SILVR^19^ learn denoising processes over molecular graphs or three-dimensional coordinates. Autoregressive and pocket-conditioned models such as Pocket2Mol^20^ and DeepICL^21^ construct molecules stepwise within binding-site environments, whereas flow-based and flow-matching approaches, including MoFlow^22^, GraphAF^23^, GraphNVP^24^, FlowMol3^25^, DrugFlow^26^, PFM^27^, FLOWR^28^ and FastFlows^29^, learn continuous transformations from simple distributions to molecular representations. In addition, ligand/reference-conditioned methods such as MolJO^30^, ShEPhERD^31^ and REINVENT4^32^ Mol2Mol support analogue generation, while synthesis-aware models such as ReaSyn^33^, SynFormer^34^, and PrexSyn^35^ incorporate chemical reaction feasibility into molecular design. This rapid methodological expansion creates new opportunities for molecular discovery but also complicates objective comparison across model families.

The translational relevance of AI-enabled drug discovery is increasingly supported by clinical-stage examples. AI-assisted pipelines have contributed to target identification, hit generation, lead optimization and candidate prioritization across several therapeutic programs involving generative chemistry, reinforcement learning, physics-informed modeling and structure-based optimization. A prominent example is rentosertib (ISM001-055/INS018_055), an AI-discovered and AI-designed inhibitor of TRAF2- and NCK-interacting kinase (TNIK) developed for idiopathic pulmonary fibrosis Rentosertib^36, 37^. Rentosertib demonstrated acceptable safety and tolerability in a randomized Phase IIa trial, together with an exploratory dose-dependent lung-function signal, and subsequently progressed toward Phase III development Rentosertib^36, 37^. Its clinical advancement provides a translationally relevant positive-control opportunity for examining whether an integrated computational workflow can recover a clinically advanced molecule using complementary chemical-quality, structural and dynamics-aware criteria. We therefore retrospectively evaluated rentosertib using our QED, synthetic-accessibility, docking and state-aware functional classifier (SAFC) framework, while recognizing that this single-compound analysis does not constitute prospective validation. Other examples, including ISM3091^38^, REC-1245^39^, SGR-1505^40^ and lirafugratinib^41^, further illustrate the growing role of AI-guided workflows in advancing candidates toward clinical evaluation. However, many clinically advanced AI-discovered molecules originate from proprietary platforms whose model architectures, training data and optimization procedures are not fully accessible. This limited transparency underscores the need for systematic benchmarking of open and reproducibly executable generative models across diverse targets, input conditions and medicinal-chemistry criteria.

Despite the growing number of generative molecular design tools, systematic comparison remains challenging. Existing methods differ substantially in their design objective, molecular representation, input requirements, conditioning strategy, training data and output format. Some models generate molecules without target-specific conditioning, some require seed ligands or textual prompts, some generate three-dimensional ligands directly in protein pockets, and others optimize chemical structures using ligand similarity or reaction feasibility. As a result, apparent performance differences may reflect differences in task formulation rather than differences in molecular design capability. Because complete input standardization would erase the intended task definition of several methods, a fair benchmark requires transparent documentation of native conditioning inputs, consistent output processing, and a common downstream evaluation framework for operational robustness, molecular properties, diversity, docking-score performance, and predicted ADMET profiles.

Here, we present a systematic benchmark of AI-based molecular generation and optimization methods for structure-guided drug discovery. We curated 176 protein– ligand systems from AlignDockBench^42^, PoseBusters^43^ and PLINDER^44^, covering diverse protein classes, including kinases, hydrolases, transferases, oxidoreductases, GPCRs, proteases, nuclear receptors and transporters. We evaluated 12 representative molecular design tools spanning pocket-conditioned 3D generation, diffusion- and flow-based modeling, autoregressive molecular construction, ligand- or reference-conditioned optimization, SMILES-based generation and synthesis-aware design (Table 1). All outputs were processed through a unified evaluation pipeline measuring operational robustness, molecular validity, uniqueness, chemical and scaffold diversity, docking performance, QED, synthetic accessibility, physicochemical properties, ADME and toxicity liabilities, and computational scalability. To extend evaluation beyond static docking and conventional molecular descriptors, we developed a state-aware functional classifier (SAFC) that integrates MD-derived receptor ensembles, ensemble docking and protein–ligand interaction representations for dynamics-aware candidate prioritization. SAFC was examined in an exploratory BRAF-specific application and further applied retrospectively to rentosertib as a clinically grounded positive-control case, these analyses were intended to demonstrate downstream prioritization rather than establish calibrated prediction of biological activity. Collectively, this benchmark provides practical guidance for selecting models according to available inputs and design objectives and proposes a hybrid preclinical workflow integrating molecular generation, chemical-quality assessment, structure-based evaluation, ADME and toxicity filtering, dynamics-aware functional prioritization and synthesis-aware candidate selection.

**Table 1.**
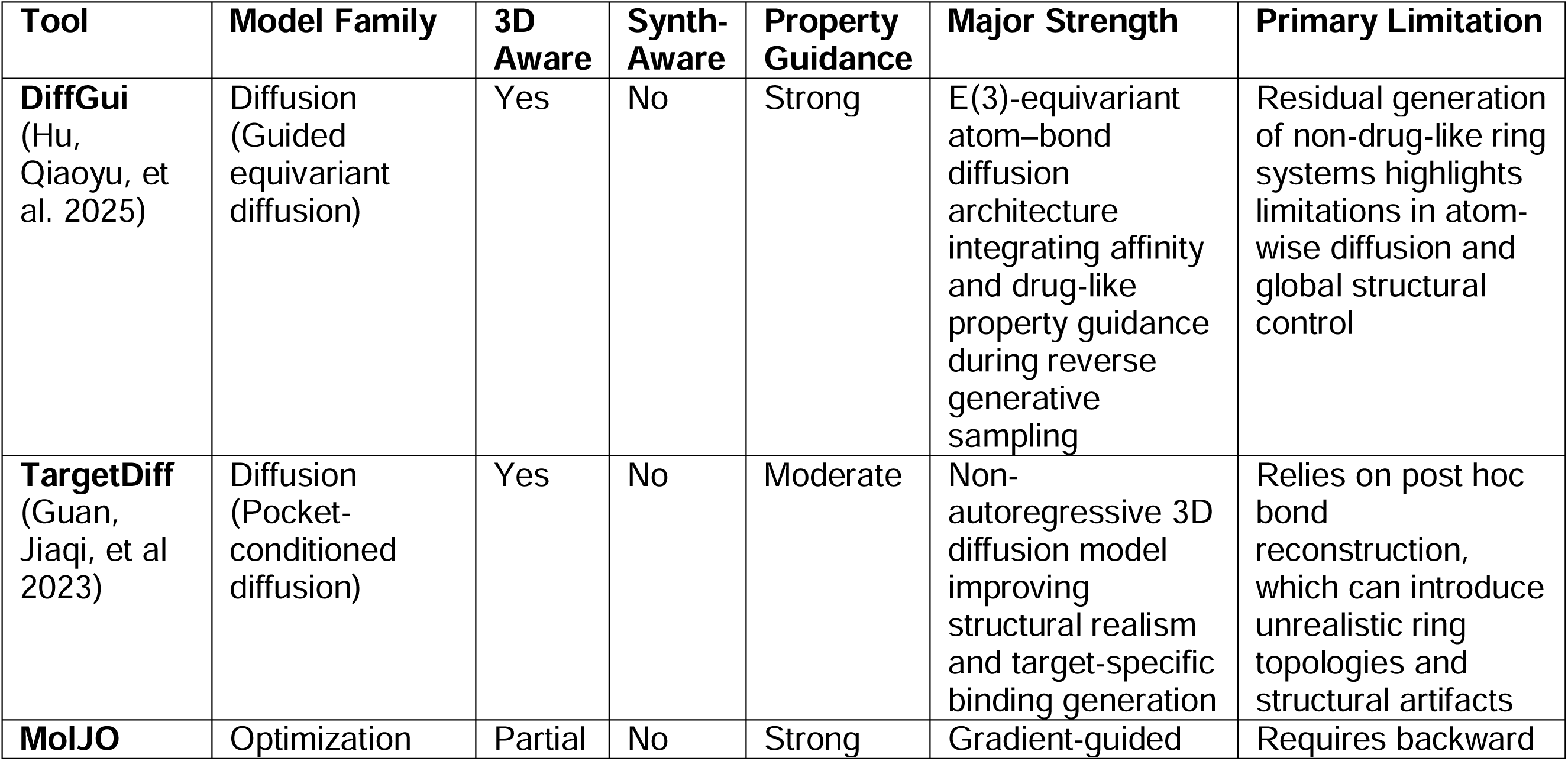
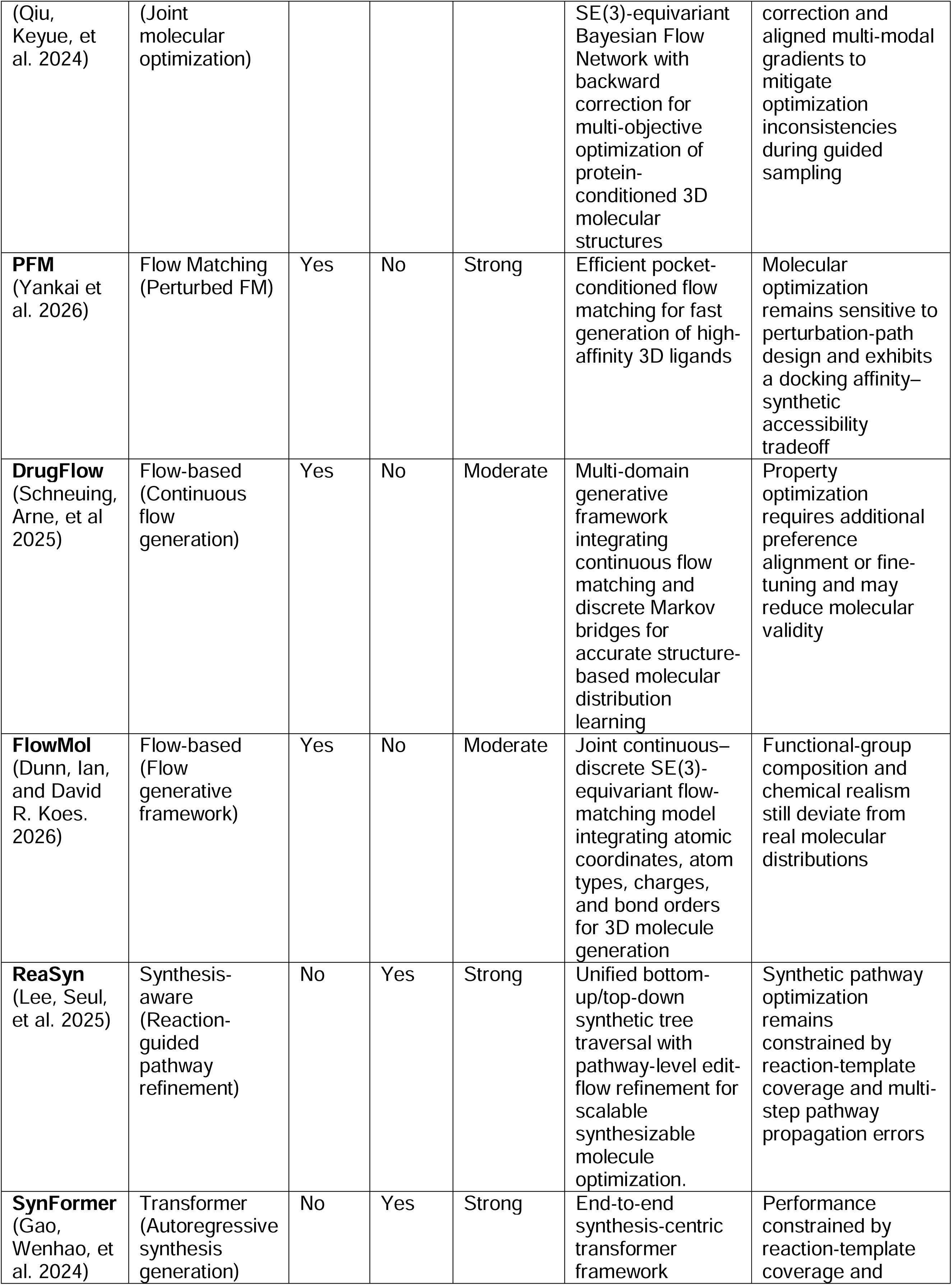
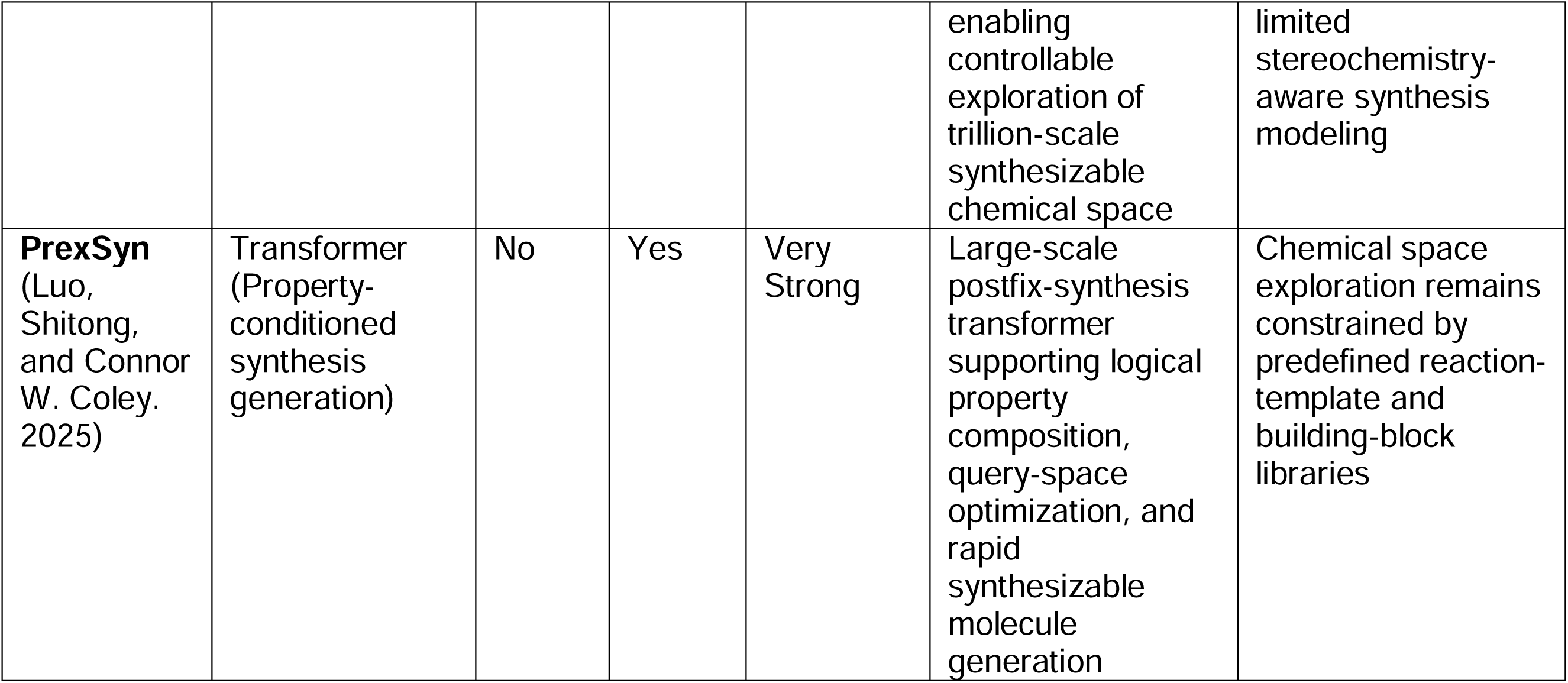
Comparative Benchmarking of AI-Based Generative Models for Structure-Based Drug Discovery. Comprehensive comparison of diffusion-based, flow-based, optimization-based, and synthesis-aware generative frameworks evaluated in this benchmarking study.

| **Tool** | **Model Family** | **3D Aware** | **Synth-Aware** | **Property Guidance** | **Major Strength** | **Primary Limitation** |
| --- | --- | --- | --- | --- | --- | --- |
| **DiffGui**<br>(Hu, Qiaoyu, et al. 2025) | Diffusion<br>(Guided equivariant diffusion) | Yes | No | Strong | E(3)-equivariant atom–bond diffusion architecture integrating affinity and drug-like property guidance during reverse generative sampling | Residual generation of non-drug-like ring systems highlights limitations in atom-wise diffusion and global structural control |
| **TargetDiff**<br>(Guan, Jiaqi, et al 2023) | Diffusion<br>(Pocket-conditioned diffusion) | Yes | No | Moderate | Non-autoregressive 3D diffusion model improving structural realism and target-specific binding generation | Relies on post hoc bond reconstruction, which can introduce unrealistic ring topologies and structural artifacts |
| **MolJO** | Optimization | Partial | No | Strong | Gradient-guided | Requires backward |
| (Qiu, Keyue, et al. 2024) | (Joint molecular optimization) |  |  |  | SE(3)-equivariant Bayesian Flow Network with backward correction for multi-objective optimization of protein-conditioned 3D molecular structures | correction and aligned multi-modal gradients to mitigate optimization inconsistencies during guided sampling |
| **PFM**<br>(Yankai et al. 2026) | Flow Matching (Perturbed FM) | Yes | No | Strong | Efficient pocket-conditioned flow matching for fast generation of high-affinity 3D ligands | Molecular optimization remains sensitive to perturbation-path design and exhibits a docking affinity–synthetic accessibility tradeoff |
| **DrugFlow**<br>(Schneuing, Arne, et al 2025) | Flow-based (Continuous flow generation) | Yes | No | Moderate | Multi-domain generative framework integrating continuous flow matching and discrete Markov bridges for accurate structure-based molecular distribution learning | Property optimization requires additional preference alignment or fine-tuning and may reduce molecular validity |
| **FlowMol**<br>(Dunn, Ian, and David R. Koes. 2026) | Flow-based (Flow generative framework) | Yes | No | Moderate | Joint continuous–discrete SE(3)-equivariant flow-matching model integrating atomic coordinates, atom types, charges, and bond orders for 3D molecule generation | Functional-group composition and chemical realism still deviate from real molecular distributions |
| **ReaSyn**<br>(Lee, Seul, et al. 2025) | Synthesis-aware (Reaction-guided pathway refinement) | No | Yes | Strong | Unified bottom-up/top-down synthetic tree traversal with pathway-level edit-flow refinement for scalable synthesizable molecule optimization. | Synthetic pathway optimization remains constrained by reaction-template coverage and multi-step pathway propagation errors |
| **SynFormer**<br>(Gao, Wenhao, et al. 2024) | Transformer (Autoregressive synthesis generation) | No | Yes | Strong | End-to-end synthesis-centric transformer framework | Performance constrained by reaction-template coverage and |
|  |  |  |  |  | enabling controllable exploration of trillion-scale synthesizable chemical space | limited stereochemistry-aware synthesis modeling |
| **PrexSyn**<br>(Luo, Shitong, and Connor W. Coley. 2025) | Transformer (Property-conditioned synthesis generation) | No | Yes | Very Strong | Large-scale postfix-synthesis transformer supporting logical property composition, query-space optimization, and rapid synthesizable molecule generation | Chemical space exploration remains constrained by predefined reaction-template and building-block libraries |

## Results

### A task-aware framework for benchmarking AI-based molecular design models

We first established a benchmarking framework to compare AI-based molecular generation and optimization models across consistent target sets, input conditions and downstream evaluation criteria. Because de novo molecular generation does not have a single experimental “accuracy” label, we evaluated each model across three complementary dimensions: molecular quality, generation robustness and practical scalability. Molecular quality was assessed using chemical validity, scaffold diversity, QED, normalized synthetic-accessibility desirability, docking-score distributions and ADME/toxicity profiles. Robustness was evaluated by measuring whether each method could reproducibly generate processable molecules across diverse protein targets and binding-site environments. Scalability was assessed using runtime, throughput, GPU-memory usage, job-completion behavior and output size. Molecular generation was standardized across 176 protein–ligand systems, with up to 100 molecules requested per target under controlled sampling and computational settings. Detailed configurations, benchmark composition, resource allocation and retained outputs are provided in Supplementary Table 1. This framework allowed heterogeneous model families, including pocket-conditioned 3D generators, diffusion- and flow-based models, ligand/reference-conditioned methods, SMILES-based optimization tools and synthesis-aware generators, to be compared within a unified structure-guided drug discovery pipeline as shown in Fig. 1.

**Fig. 1.**
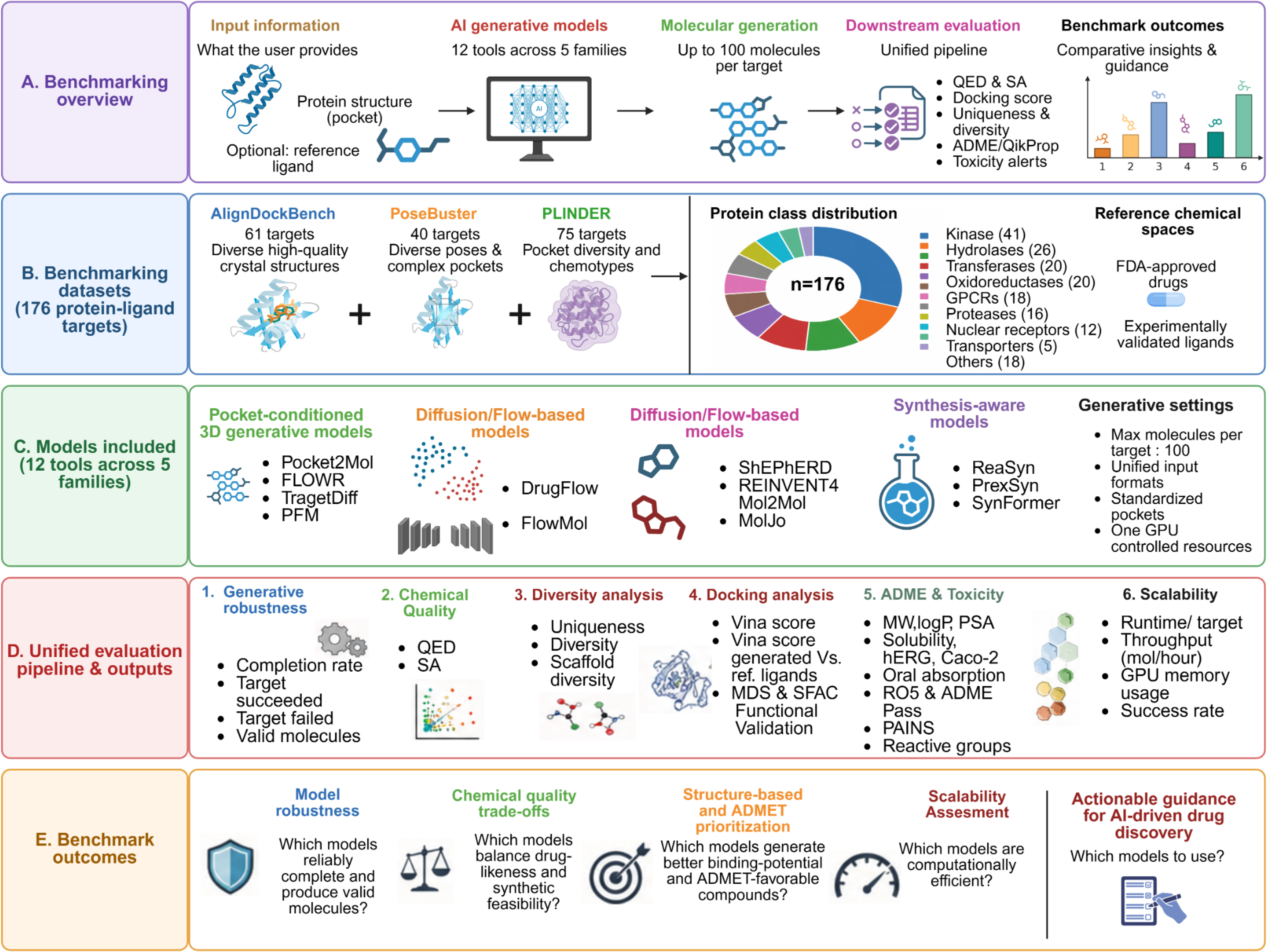
Benchmarking framework and study overview. **a,** Overview of the standardized workflow, from protein-pocket and optional reference-ligand inputs through AI-based molecular generation and downstream assessment of chemical quality, diversity, docking, ADME and toxicity. Each model was requested to generate up to 100 molecules per target. **b,** Benchmark composition, integrating 61 AlignDockBench, 40 PoseBusters and 75 PLINDER protein–ligand systems spanning diverse protein classes, binding pockets and reference chemical spaces. **c,** Molecular generative models included in the study, grouped by architectural or task-design family, together with the standardized generation settings used for comparison. **d,** Unified evaluation framework comprising generative robustness, chemical quality, molecular diversity, docking performance, ADME and toxicity profiling, and computational scalability. **e,** Principal benchmarking outcomes used to characterize model robustness, chemical-quality trade-offs, structure-based prioritization, computational efficiency and practical model-selection guidance for AI-enabled drug discovery.

### Benchmark dataset construction

The benchmarking dataset comprised 176 structurally diverse protein–ligand systems from AlignDockBench^42^ (ADB; n = 61), PoseBusters^43^ (PB; n = 40) and PLINDER^44^ (PL; n = 75), spanning kinases, hydrolases, transferases, oxidoreductases, GPCRs, proteases, nuclear receptors and additional protein classes (Fig. 2a,b, & Supplementary Notes S1 & S2). The three cohorts sampled complementary chemical spaces, with PL exhibiting the broadest molecular-weight and physicochemical-property distributions, whereas ADB contained a more compact drug-like chemical space (Fig. 2c–f). High Murcko scaffold-to-ligand ratios in ADB (58/60) and PB (37/40), together with a lower but still substantial diversity in PL (47/62), indicated limited scaffold redundancy overall, with comparatively greater scaffold reuse in PL (Fig. 2g). Medicinal-chemistry filters were most frequently satisfied by ADB ligands, while the broader PL chemical space produced lower Lipinski, Veber, molecular-weight, cLogP and TPSA pass rates, establishing a heterogeneous benchmark with distinct structural and chemical challenges for generative-model evaluation (Fig. 2h).

**Fig. 2.**
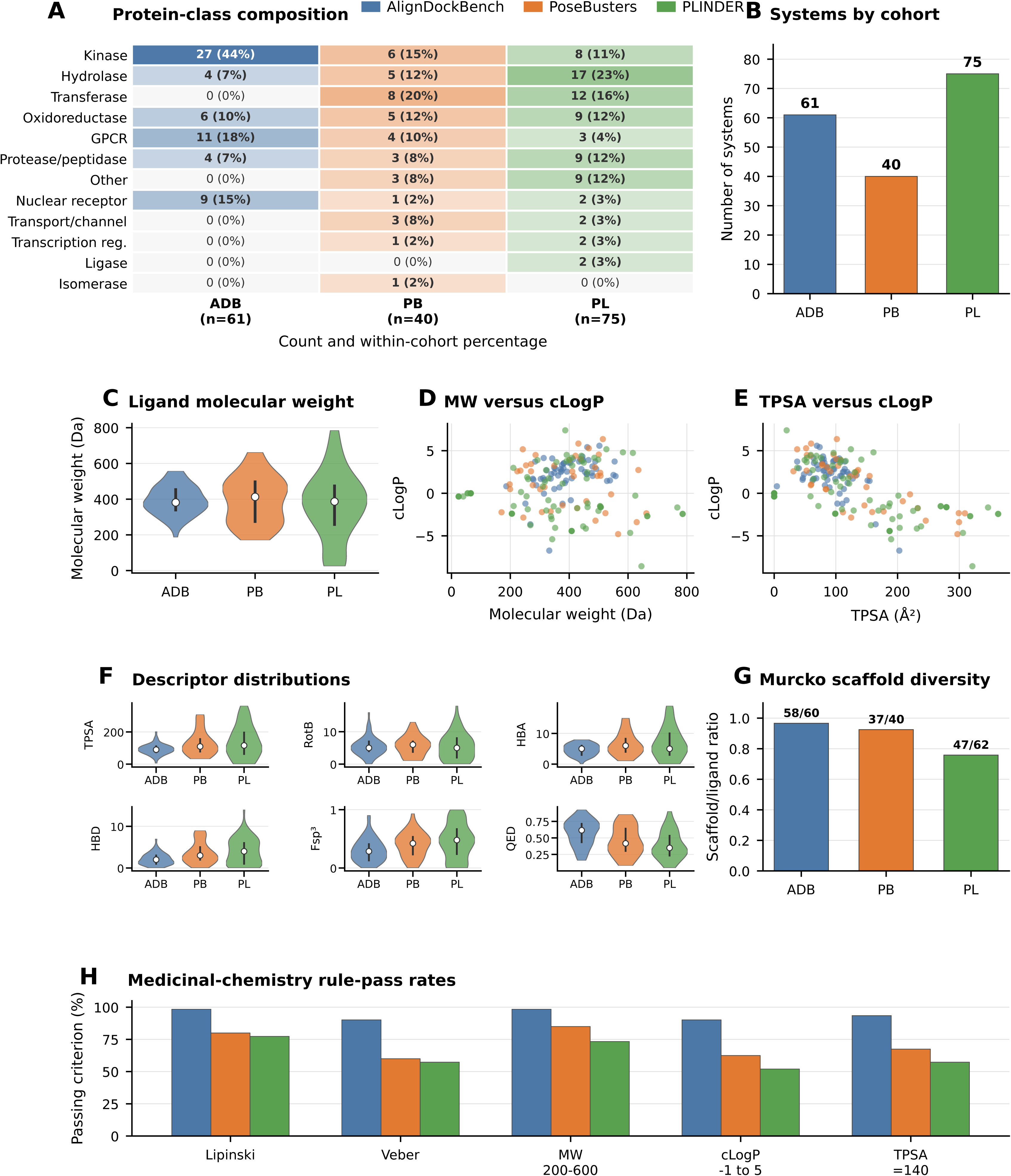
Composition and chemical diversity of the benchmarking dataset. **a,** Protein-class composition of the AlignDockBench (ADB), PoseBusters (PB) and PLINDER (PL) cohorts. Cells report the number and within-cohort percentage of systems assigned to each protein class. **b,** Number of protein–ligand systems in ADB (n = 61), PB (n = 40) and PL (n = 75), comprising 176 benchmark targets in total. **c,** Distribution of ligand molecular weights across the three cohorts. **d, e,** Relationships between molecular weight and calculated logP (cLogP) (d) and between topological polar surface area (TPSA) and cLogP (e). **f,** Cohort-level distributions of TPSA, rotatable-bond count (RotB), hydrogen-bond acceptors (HBA), hydrogen-bond donors (HBD), fraction of sp³-hybridized carbon atoms (Fsp³) and quantitative estimate of drug-likeness (QED). **g,** Murcko scaffold diversity, expressed as the ratio of unique scaffolds to ligands; labels indicate the corresponding scaffold and ligand counts. **h,** Percentage of ligands satisfying the Lipinski and Veber criteria and the indicated molecular weight, cLogP and TPSA thresholds.

### Generative models differ in target and valid output robustness

We evaluated the operational robustness of 12 molecular generation models across 176 protein targets by measuring target completion and the delivery of RDKit-valid molecules (Supplementary Table2). Pocket2Mol, FlowMol and MolJO achieved the highest completion rates, each successfully processing 175 targets (99.4%), followed by TargetDiff and PFM at 98.9% and 98.3%, respectively (Fig. 3a). By contrast, ReaSyn and REINVENT4 completed 86.9% and 64.8% of targets. Completion, however, did not consistently translate into full output delivery. Pocket2Mol and FlowMol supplied at least 100 valid molecules for 99% and 98% of targets, whereas TargetDiff reached this quota for only 1% despite its near-complete target coverage (Fig. 3b). MolJO, ShEPhERD and FLOWR also showed strong high-quota performance, while ReaSyn and REINVENT4 exhibited lower attainment across progressively stringent output thresholds. Cohort-stratified analysis showed that Pocket2Mol and FlowMol maintained robust performance across ADB, PB and PL, whereas several models—including TargetDiff, PFM, SynFormer, PrexSyn and DrugFlow—showed reduced high-quota attainment in the PL cohort (Fig. 3c). Target-level outcome composition further distinguished models that consistently delivered near-complete molecular sets from those producing partial or absent outputs (Fig. 3d). Pocket2Mol, FlowMol and MolJO generated near-complete valid sets for almost all targets, whereas TargetDiff frequently returned usable but sub-quota outputs, and ReaSyn and REINVENT4 showed the largest fractions of partial or missing results. Chemical validity among emitted molecules was nearly saturated across models (99.8–100%). Together, these findings show that execution success alone is insufficient to characterize generative-model robustness and should be evaluated alongside the quantity of valid molecules delivered for downstream prioritization. Some of the models are excluded from benchmarking and their details are provided in Supplementary Note S3.

**Fig. 3.**
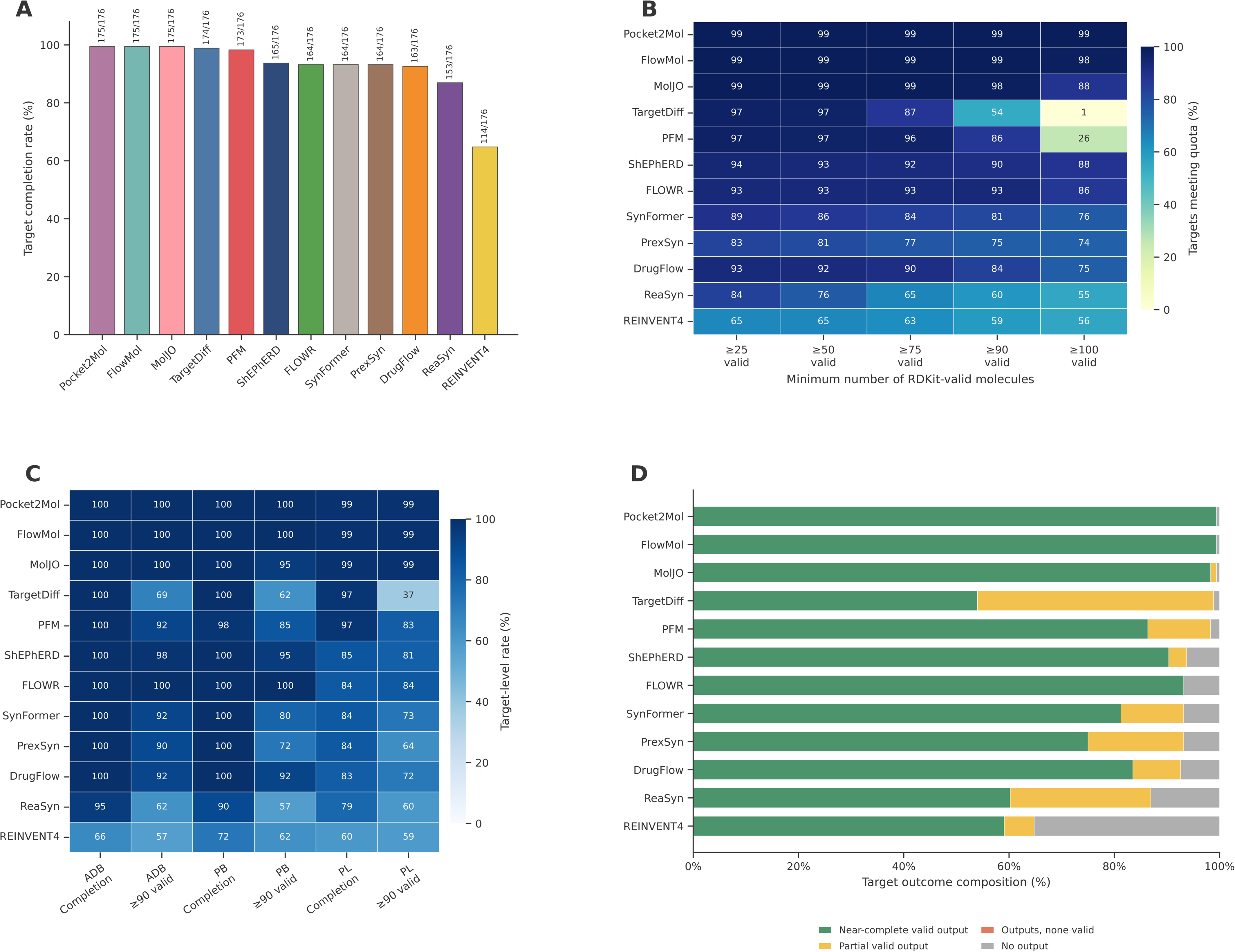
Comparative assessment of molecular generation robustness and output validity. **a,** Target completion rates across 176 benchmark targets. Labels above the bars indicate the number of completed targets relative to the full target set. **b,** Percentage of all benchmark targets for which each model generated at least 25, 50, 75, 90 or 100 RDKit-valid molecules. **c,** Target completion and ≥90-valid-molecule attainment rates stratified by the AlignDockBench (ADB; n = 61), PoseBusters (PB; n = 40) and PLINDER (PL; n = 75) cohorts. **d,** Target-level outcome composition classified as near-complete valid output (≥90 valid molecules), partial valid output (1–89 valid molecules), output containing no valid molecules, or no output. Chemical validity among emitted molecules ranged from 99.8% to 100% across models.

### Generative models occupy distinct chemical quality–diversity regimes

We assessed descriptor-based molecular quality and chemical-space exploration in a complete-case cohort of 54 targets for which all 12 models produced sufficient valid outputs for matched analysis. QED, normalized synthetic accessibility (SA), Morgan-fingerprint diversity and normalized Bemis–Murcko scaffold entropy were summarized at the target level; diversity estimates were calculated using repeated equal-size subsampling to control for differences in output size. The models showed marked differences in both QED and SA (Fig. 4a,b). FlowMol achieved the highest median QED and normalized SA, followed by Pocket2Mol, whereas DrugFlow, FLOWR and ShEPhERD displayed intermediate profiles. PrexSyn, SynFormer and ReaSyn produced lower and more target-variable QED distributions, while TargetDiff and PFM showed comparatively low normalized SA. Joint analysis confirmed that drug-likeness and synthetic accessibility were only partially coupled: FlowMol was the only model whose median profile clearly exceeded both the QED (0.70) and normalized SA (0.667) thresholds, with Pocket2Mol positioned close to this high-quality region (Fig. 4c). Chemical-space exploration separated the models into distinct diversity regimes. TargetDiff, PFM and ShEPhERD generated the most fingerprint-diverse molecular populations, with Pocket2Mol, MolJO, DrugFlow, FLOWR and FlowMol also maintaining relatively high diversity (Fig. 4d). ReaSyn, REINVENT4, SynFormer and PrexSyn produced more concentrated chemical distributions. Scaffold entropy showed a related but non-identical pattern: ShEPhERD and PFM retained the broadest scaffold repertoires, followed by TargetDiff, MolJO and FlowMol, whereas the synthesis-aware models and REINVENT4 exhibited lower and more variable scaffold diversity across targets (Fig. 4e). These differences indicate that high molecular diversity does not necessarily imply equivalent exploration of core chemical frameworks.

**Fig. 4.**
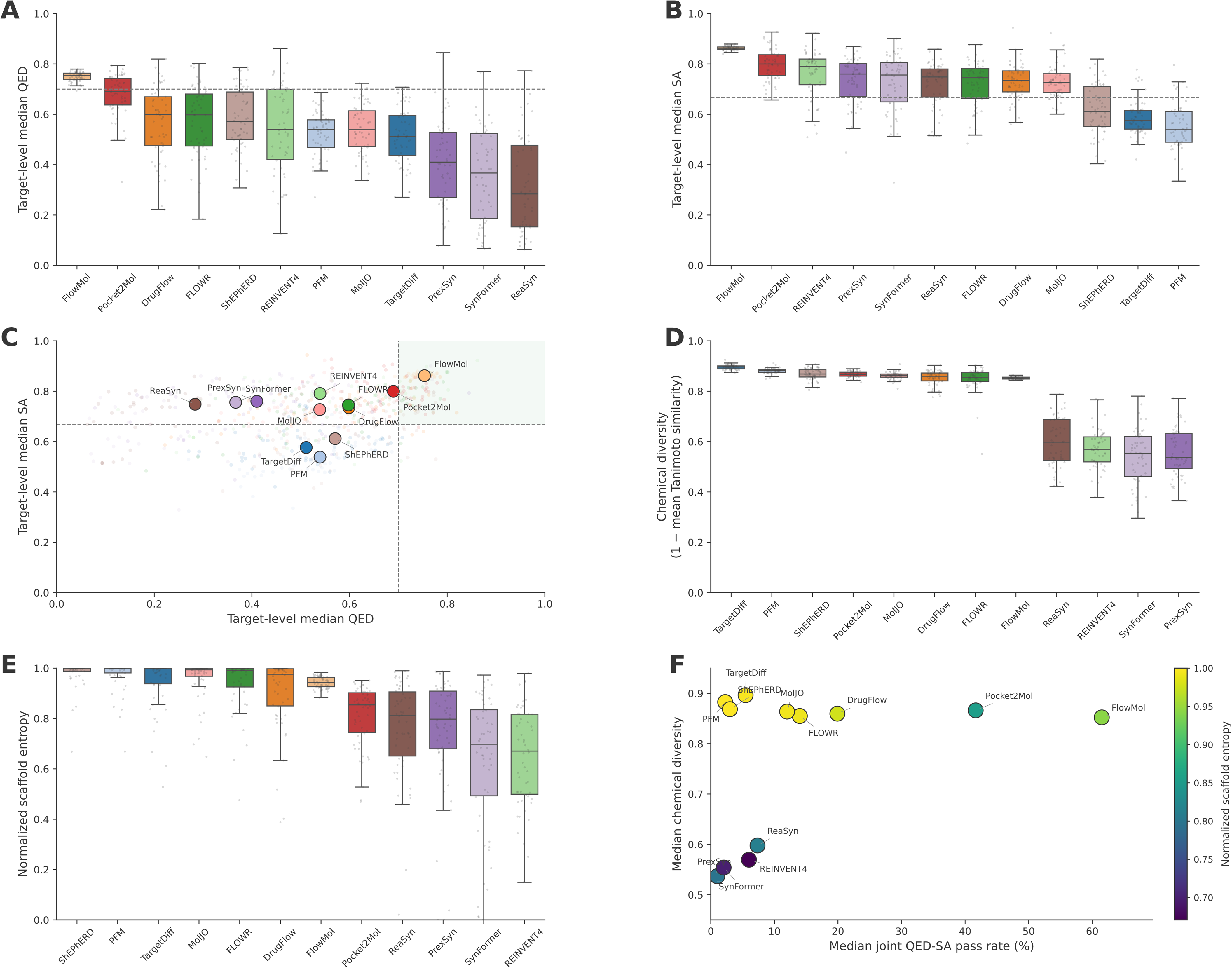
Drug-likeness, synthetic accessibility and chemical scaffold diversity across generative models. Analyses were restricted to 54 targets represented by all 12 models. **a,b,** Target-level distributions of median quantitative estimate of drug-likeness (QED; a) and normalized synthetic accessibility (SA; b), where higher values indicate more favorable properties. Dashed lines denote the predefined QED (0.70) and normalized SA (0.667) thresholds. **c,** Joint distribution of target-level median QED and normalized SA. Small points represent individual model–target combinations, and labelled circles indicate model-level medians; the shaded upper-right region denotes simultaneous passage of both thresholds. **d,** Target-level chemical diversity, calculated as one minus the mean pairwise Morgan-fingerprint Tanimoto similarity. **e,** Target-level normalized Bemis–Murcko scaffold entropy. Chemical and scaffold diversity metrics were estimated using repeated equal-size subsampling within each model–target combination to control for unequal output sizes. **f,** Integrated quality–diversity landscape showing the median joint QED–SA pass rate, median chemical diversity and normalized scaffold entropy, represented by the x axis, y axis and point colour, respectively. In a, b, d and e, centre lines indicate medians, boxes represent the interquartile range, whiskers extend to 1.5 times the interquartile range and points denote individual targets.

Integration of quality and diversity metrics revealed model-specific operating regimes rather than a single dominant method (Fig. 4f). FlowMol provided the strongest overall balance, achieving the highest median joint QED–SA pass rate while retaining high chemical and scaffold diversity. Pocket2Mol showed the next most favorable compromise, whereas DrugFlow occupied an intermediate quality–diversity region. TargetDiff, PFM and ShEPhERD prioritized broad molecular and scaffold exploration but generated smaller proportions of molecules satisfying both quality thresholds. Conversely, ReaSyn, REINVENT4, SynFormer and PrexSyn combined low joint pass rates with comparatively restricted chemical-space exploration. Thus, model selection should reflect the intended discovery objective: FlowMol and Pocket2Mol are better suited to quality-focused generation, whereas TargetDiff, PFM and ShEPhERD may be valuable when structural exploration is prioritized. These findings support multi-objective evaluation rather than ranking generative models using drug-likeness, synthesizability or diversity in isolation.

### Integrated evaluation uncovers trade-offs between docking performance and molecular quality

We integrated docking performance, drug-likeness and synthetic accessibility across 65 targets represented by all 12 models (Supplementary Notes S4 to S15). Target-normalized AutoDock Vina docking scores revealed substantial model-dependent differences (Fig. 5a). PFM achieved the strongest median docking performance, followed by MolJO, whereas TargetDiff, DrugFlow and ShEPhERD showed more modest positive values. FlowMol, Pocket2Mol, ReaSyn and SynFormer had lower median normalized performance, although their broad distributions indicated target-dependent variation. Within-target ranking showed that PFM and MolJO most frequently occupied the highest docking-score ranks, while the relative ordering of the remaining models varied considerably across targets (Fig. 5b). Integration with QED and normalized SA exposed a marked trade-off between docking and molecular quality (Fig. 5c). FlowMol and Pocket2Mol occupied the most favorable QED–SA region but showed comparatively weak target-normalized docking performance. Conversely, PFM combined the strongest docking performance with low median QED and SA, while MolJO combined favorable docking scores with intermediate descriptor-based molecular quality. These differences were reflected in the target-level selection rates (Fig. 5d). FlowMol achieved the highest median joint QED–SA–Vina pass rate (44.4%), followed by Pocket2Mol (39.0%). MolJO and DrugFlow showed lower triple-pass rates of 13.0% and 10.0%, respectively, despite satisfying the Vina threshold for approximately 99% of target-level molecule sets. PFM similarly passed the docking criterion for 99.0% of targets but achieved only a 2.0% median triple-pass rate because comparatively few molecules met the QED and SA thresholds. Thus, strong docking scores alone did not predict multi-parameter prioritization success. Model specific QED, SA, and Vina docking scores have been shown in Supplementary Fig. S1 to S12.

**Fig. 5.**
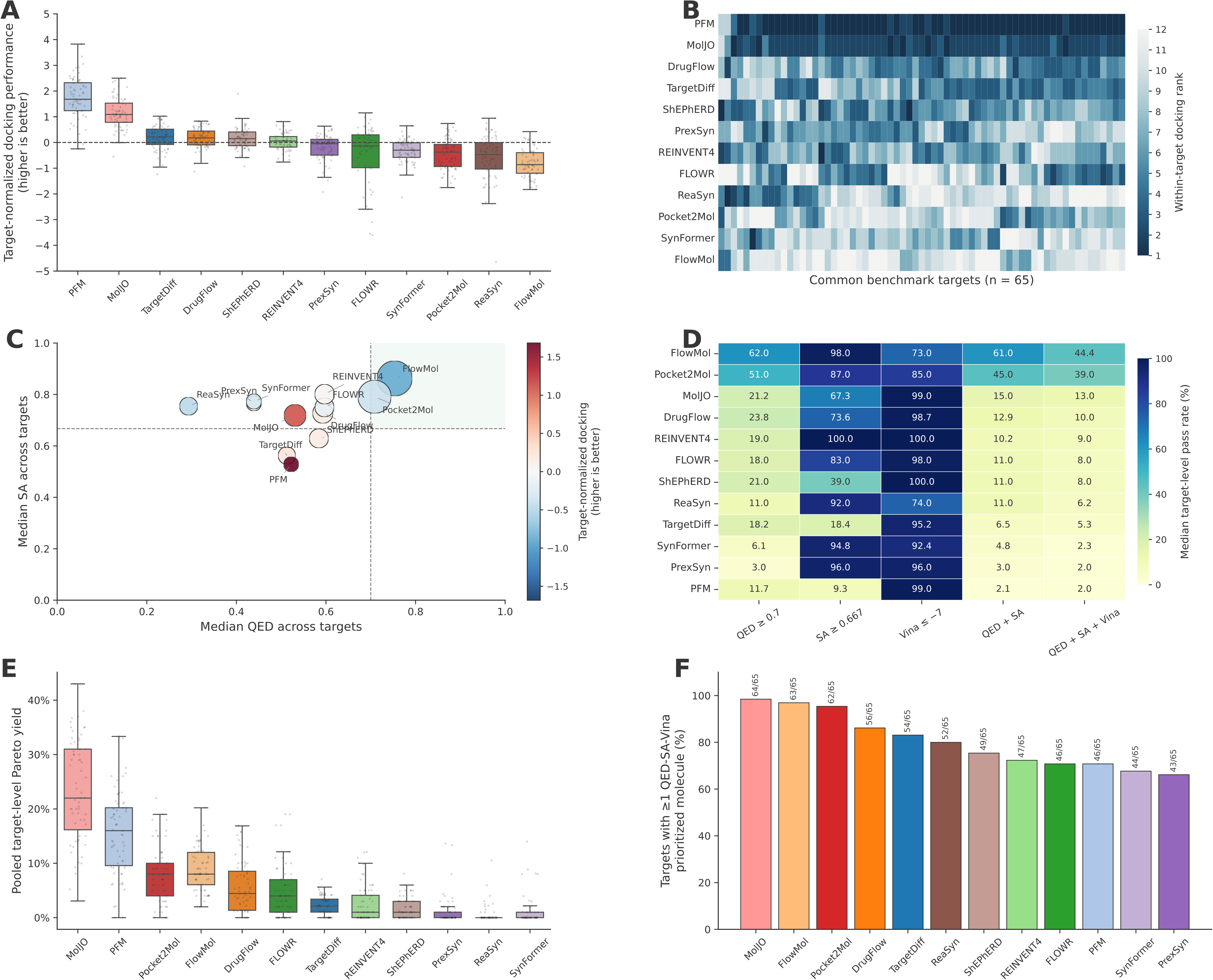
Integrated evaluation of drug-likeness, synthetic accessibility and docking performance. Analyses were restricted to 65 benchmark targets represented by all 12 models. **a,** Distribution of target-normalized docking performance derived from target-level median AutoDock Vina scores, with higher values indicating better relative docking performance. **b,** Within-target ranking of models by median docking performance across the common targets; rank 1 indicates the best-performing model. **c,** Integrated comparison of model-level median QED and normalized synthetic accessibility (SA). Point colour represents target-normalized docking performance, and point size indicates the median joint QED–SA–Vina pass rate. Dashed lines and the shaded region denote the QED ≥ 0.70 and normalized SA ≥ 0.667 selection thresholds. **d,** Median target-level percentage of molecules satisfying the individual QED, normalized SA and Vina criteria and their indicated combinations. The docking criterion was defined as a Vina score ≤−7 kcal mol⁻¹. **e,** Target-standardized Pareto-optimal yield considering QED, normalized SA and docking performance simultaneously. **f,** Percentage of targets for which each model generated at least one molecule satisfying all three criteria; labels indicate the corresponding number of successful targets among the 65 common targets. In a and e, centre lines indicate medians, boxes represent the interquartile range, whiskers extend to 1.5 times the interquartile range and points denote individual targets.

Pareto analysis provided a complementary measure that did not require simultaneous passage of fixed thresholds (Fig. 5e). MolJO produced the largest target-standardized Pareto-optimal yield, followed by PFM, indicating that both models frequently generated molecules representing favorable compromises among docking, QED and SA. By contrast, FlowMol and Pocket2Mol achieved lower Pareto yields but substantially broader target coverage under the strict joint criteria. MolJO generated at least one QED–SA–Vina-prioritized molecule for 64 of 65 targets, followed by FlowMol (63), Pocket2Mol (62) and DrugFlow (56) (Fig. 5f). Collectively, these results identify distinct model operating regimes: PFM and MolJO prioritize docking and Pareto efficiency, whereas FlowMol and Pocket2Mol provide the strongest balance of drug-likeness, synthetic accessibility and docking-threshold attainment. Multi-objective model selection is therefore more informative than ranking generators by docking score or molecular quality alone.

### Predicted ADME profiling reveals model-specific trade-offs in drug-likeness, absorption and metabolic liability

We compared the predicted ADME properties of molecules generated by all 12 models across a common cohort of 97 targets (Supplementary Note S16). Pocket2Mol, FlowMol and ShEPhERD produced the largest proportions of molecules with no more than one Lipinski Rule-of-Five violation, whereas ReaSyn and SynFormer generated substantially larger fractions with two or more violations (Fig. 6a). Continuous-property distributions further revealed systematic model-dependent shifts in predicted aqueous solubility, Caco-2 permeability, plasma protein binding, volume of distribution, hepatocyte clearance and half-life (Fig. 6b). Although the distributions overlapped, their differences indicate that the models sampled distinct physicochemical and pharmacokinetic property regimes. Predicted human intestinal absorption and bioavailability were consistently high across models, ranging from 88.9–99.9% and 87.7–97.8%, respectively (Fig. 6c). FlowMol achieved 99.0% predicted intestinal absorption and 97.8% bioavailability, while Pocket2Mol combined similarly high values (97.6% and 97.1%) with the lowest predicted P-glycoprotein inhibition prevalence (9.6%). MolJO showed the highest predicted intestinal absorption (99.9%) but also the greatest P-glycoprotein inhibition liability (57.4%), illustrating that favorable absorption predictions did not necessarily coincide with reduced transporter-mediated risk. Predicted blood– brain barrier penetration varied from 49.1% for ReaSyn to 95.9% for FlowMol; this endpoint represents a distribution characteristic rather than a universally favorable property and should be interpreted in the context of the intended therapeutic indication. Major CYP inhibition profiles provided an additional source of model separation (Fig. 6d). TargetDiff, PFM, Pocket2Mol and ShEPhERD generated comparatively larger fractions of molecules predicted to inhibit zero or one major CYP isoform, whereas MolJO, FlowMol and SynFormer showed greater multi-CYP inhibition burdens. Target-balanced integration of absorption, distribution, metabolism and excretion endpoints identified PFM, REINVENT4, TargetDiff, ShEPhERD and Pocket2Mol among the models with the highest median ADME soft-pass rates (Fig. 6e). Across-target distributions nevertheless revealed substantial variability for several models, including SynFormer, ReaSyn, REINVENT4 and PrexSyn, whose lower tails indicated target-specific deterioration in ADME readiness (Fig. 6f). Together, these results show that no model was uniformly optimal across all pharmacokinetic dimensions: some prioritized conventional drug-likeness and reduced CYP liability, whereas others favored absorption or tissue-penetration characteristics. Multi-endpoint, target-balanced profiling is therefore necessary to identify model-specific liabilities that would be obscured by any single ADME measure. Model specific ADME profiling has been shown in Supplementary Fig. S13 to S24.

**Fig. 6.**
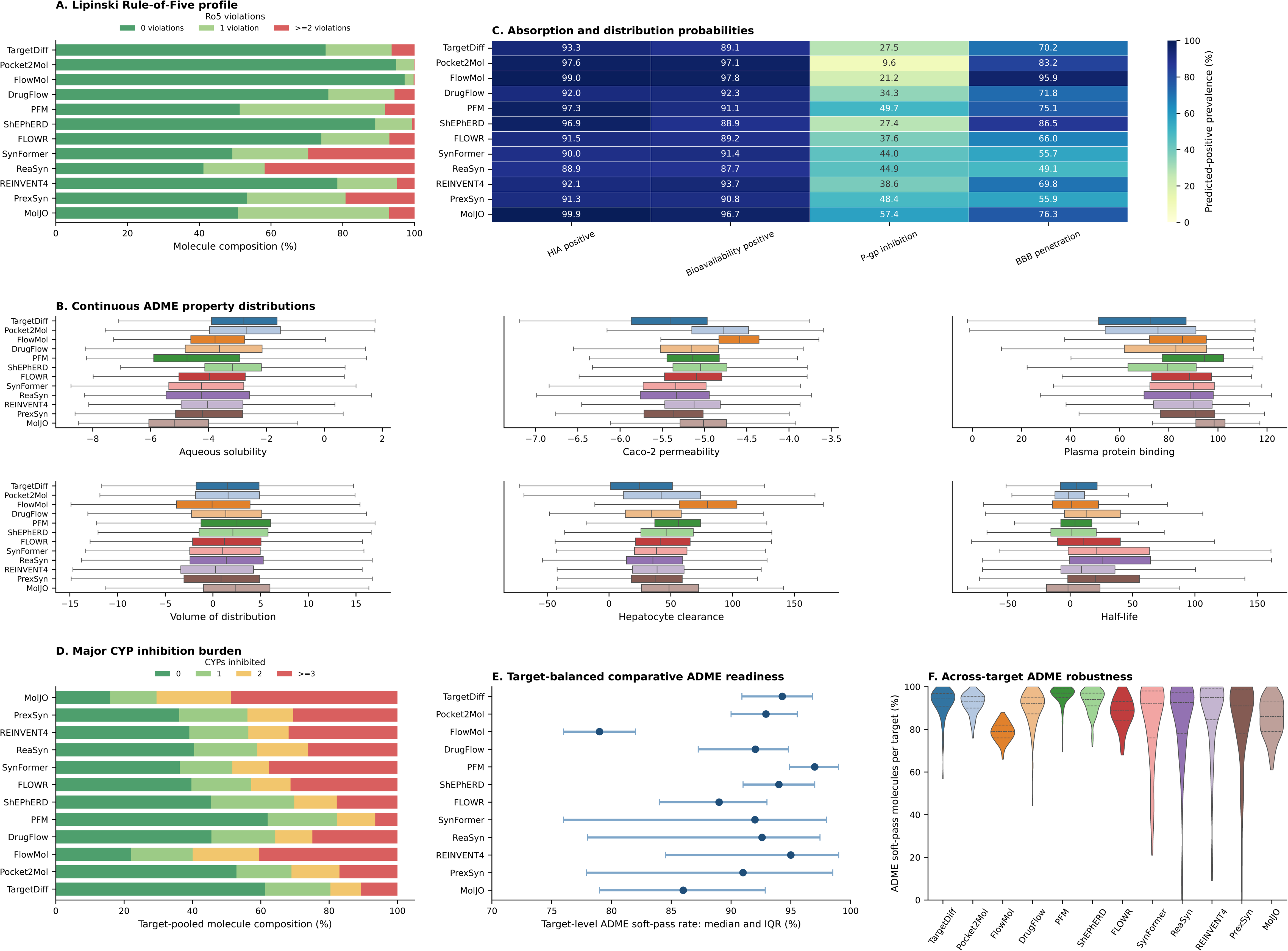
Comparative ADME profiling of generated molecules. Analyses were performed across a common cohort of 97 targets. **a,** Model-level composition of molecules containing zero, one or at least two Lipinski Rule-of-Five violations. Violations were defined as molecular weight >500 Da, calculated logP >5, more than five hydrogen-bond donors or more than ten hydrogen-bond acceptors; molecules with no more than one violation were considered Rule-of-Five compliant. **b,** Model-specific distributions of ADMET-AI-predicted aqueous solubility, Caco-2 permeability, plasma protein binding, volume of distribution, hepatocyte clearance and half-life. Centre lines indicate medians, boxes represent the interquartile range and whiskers extend to 1.5 times the interquartile range. **c,** Predicted-positive prevalence of human intestinal absorption (HIA), oral bioavailability, P-glycoprotein inhibition and blood–brain barrier penetration. Binary endpoints were classified using a probability threshold of 0.5. **d,** Model-level composition of molecules predicted to inhibit zero, one, two or at least three of the five major CYP isoforms CYP1A2, CYP2C9, CYP2C19, CYP2D6 and CYP3A4. **e,** Target-balanced ADME readiness, summarized as the median and interquartile range of target-level soft-pass rates. A molecule was considered an ADME soft pass when it satisfied at least three of the four evaluated absorption, distribution, metabolism and excretion domains, as defined in the Methods. **f,** Across-target distributions of the percentage of molecules satisfying the ADME soft-pass criterion for each model. Violin widths indicate density, and internal lines denote the quartiles.

### Toxicity profiling identifies distinct and endpoint-specific liability burdens

We profiled ADMET-AI-predicted toxicity liabilities for molecules generated by all 12 models across the common 97-target cohort (Supplementary Note S17). The endpoint-level analysis revealed pronounced differences among both toxicity categories and models (Fig. 7a). General toxicity signals were dominated by predicted drug-induced liver injury, hERG liability, skin reactions and Ames mutagenicity, whereas most nuclear-receptor endpoints showed lower prevalence. Antioxidant-response and mitochondrial-membrane-potential liabilities were the most frequent stress-response signals. MolJO displayed particularly high prevalence across multiple general-toxicity and stress-response endpoints, while TargetDiff and Pocket2Mol generally produced lower endpoint-positive rates. These predictions represent computational liability alerts rather than evidence of experimentally confirmed toxicity. Because 18 endpoints were evaluated simultaneously, most models generated a high proportion of molecules with at least one predicted liability (Fig. 7b). The median prevalence was lowest for TargetDiff and PFM but approached saturation for FlowMol, SynFormer, ReaSyn, REINVENT4, PrexSyn and MolJO. The number of positive endpoints per molecule provided greater discrimination than the binary presence of any alert (Fig. 7c). TargetDiff and Pocket2Mol generated the largest fractions of molecules with zero or one predicted liability, whereas MolJO produced the highest multi-endpoint burden, with most molecules positive for at least four endpoints and approximately half positive for six or more. The remaining models occupied intermediate regimes, indicating that a single any-liability measure can obscure substantial differences in cumulative toxicity burden.

**Fig. 7.**
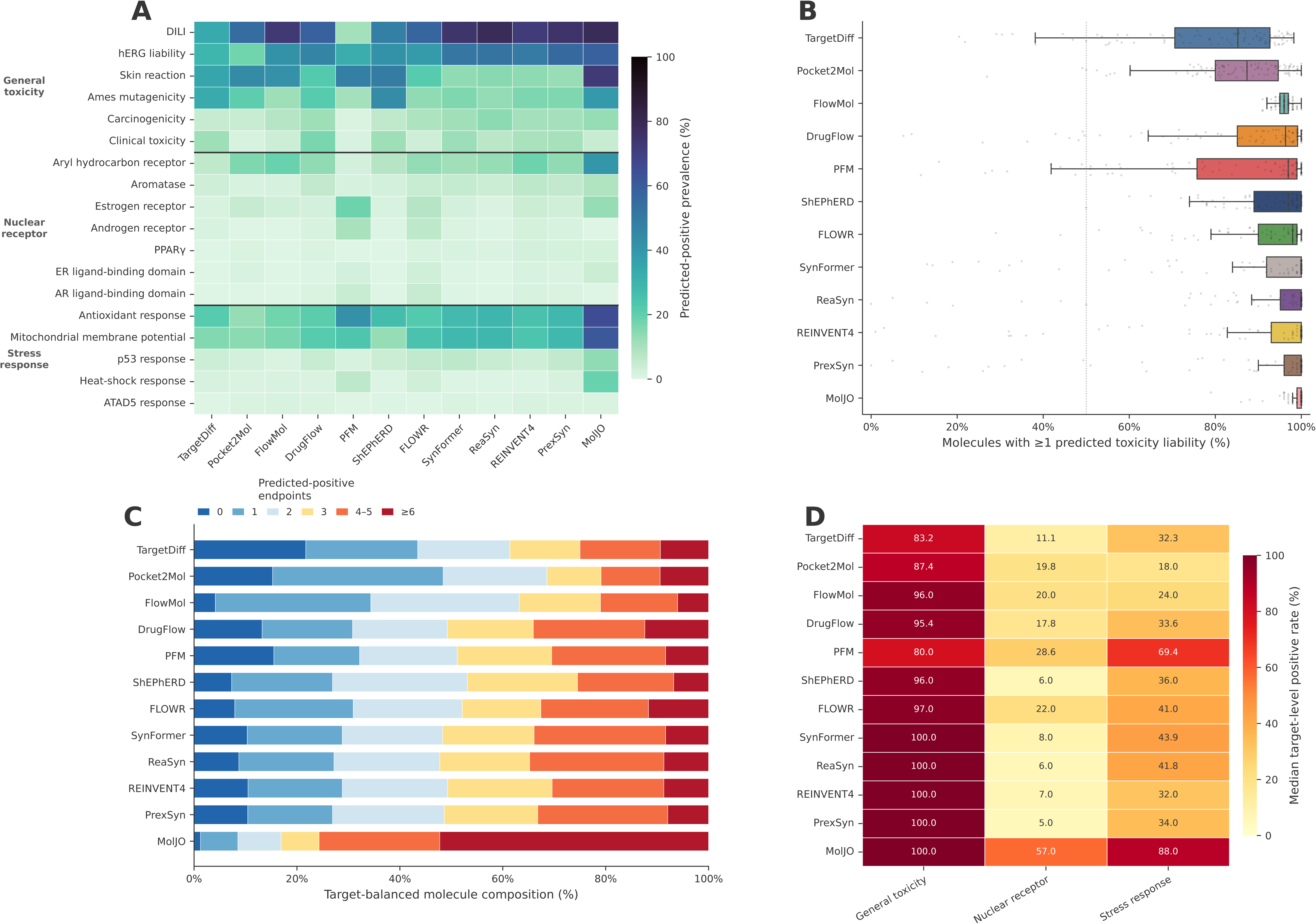
Comparative toxicity profiling identifies model- and endpoint-specific liability burdens. Analyses were performed across the common 97-target cohort using 18 ADMET-AI-predicted toxicity endpoints and a positive-classification threshold of 0.5. **a,** Predicted-positive prevalence of individual endpoints across the 12 models. Endpoints were grouped into general toxicity (drug-induced liver injury (DILI), hERG liability, skin reaction, Ames mutagenicity, carcinogenicity, clinical toxicity and aryl hydrocarbon receptor), nuclear-receptor activity (aromatase, estrogen receptor, androgen receptor, PPARγ and estrogen- and androgen-receptor ligand-binding domains) and stress response (antioxidant response, mitochondrial membrane potential, p53 response, heat-shock response and ATAD5 response). **b,** Target-level distributions of the percentage of molecules with at least one predicted toxicity liability. Centre lines denote medians, boxes represent the interquartile range, whiskers extend to 1.5 times the interquartile range and points indicate individual targets. **c,** Target-balanced molecule composition according to the cumulative number of positive toxicity endpoints per molecule, categorized as 0, 1, 2, 3, 4–5 or ≥6 liabilities. **d,** Median target-level positive rates for general-toxicity, nuclear-receptor and stress-response endpoint groups. Values represent computational liability predictions rather than experimentally confirmed toxicity.

Target-balanced category summaries further revealed that toxicity profiles were not uniformly aligned across liability classes (Fig. 7d). General-toxicity positive rates were high for all models, ranging from 80.0% for PFM to 100% for SynFormer, ReaSyn, REINVENT4, PrexSyn and MolJO. Nuclear-receptor liabilities were comparatively limited for most models, with median target-level rates of 5–29%, except for MolJO, which reached 57%. Stress-response liability varied more strongly: Pocket2Mol showed the lowest median rate (18%), whereas PFM and MolJO reached 69.4% and 88%, respectively. Thus, PFM combined relatively low general-toxicity prevalence with elevated stress-response liability, while ShEPhERD showed low nuclear-receptor activity but high general-toxicity prevalence. Collectively, these results demonstrate that toxicity cannot be represented adequately by a single aggregate score. Endpoint-resolved and burden-based profiling is required to distinguish models that generate fewer liabilities overall from those whose risks are concentrated in specific biological pathways. Model specific toxicity evaluation has been shown in Supplementary Fig. S25 to S36.

### State-aware functional prioritization distinguishes candidates beyond conventional molecular scores

To assess whether generated molecules retained potentially relevant interaction patterns across dynamic receptor conformations, we developed a state-aware functional classifier (SAFC) as a downstream prioritization module (Supplementary note S18 & Supplementary Fig.S37). SAFC integrates MD-derived receptor ensembles, representative-state selection, ensemble docking and protein–ligand interaction graphs with an EquiScore-inspired transfer-learning model, as described in the Methods. Scores range from 0 to 1, with higher values indicating stronger computational support for state-aware functional relevance. Among the 176 target–structure systems included in the benchmark, the 3SKC/BRAF system was selected as a case study for downstream SAFC-based prioritization. The selection was motivated by the availability of experimentally resolved BRAF structures representing three complementary inhibitor-bound kinase conformations: the Type I DFG-in/αC-in state represented by 2FB8, the Type II DFG-out/αC-in state represented by 1UWH, and the Type I.5 DFG-in/αC-out state represented by 3SKC^45–47^. These structurally distinct states made the 3SKC/BRAF benchmark system particularly suitable for examining whether a molecular library generated for a single benchmark structure could be further prioritized using receptor-state information beyond that structure alone. In the downstream analysis, 3SKC therefore served both as the benchmark structure from which the candidate library was generated and as one member of the three-state BRAF receptor ensemble used by SAFC, together with 2FB8 and 1UWH. Collectively, the three structures span both DFG-in and DFG-out states and both αC-in and αC-out arrangements, representing the principal inhibitor-bound BRAF conformational classes considered in this analysis. The BRAF/3SKC analysis was used as an illustrative within-target application of SAFC rather than as a cross-target validation of SAFC scores or a general ranking of the generative workflows. SAFC score distributions differed substantially across the 12 generative models (Fig. 8a). Within this BRAF/3SKC case study, SynFormer produced the highest overall score distribution, followed by PrexSyn, whereas FlowMol and Pocket2Mol generated predominantly lower-scoring candidates. REINVENT4 occupied an intermediate regime, while the remaining models displayed broad, overlapping distributions that crossed the predefined prioritization threshold of 0.5. The highest-ranked candidate from each model further demonstrated that SAFC captured information complementary to QED, synthetic accessibility and docking score. SynFormer and PrexSyn yielded the highest individual SAFC scores of 0.817 and 0.765, respectively, whereas the top PFM and MolJO candidates combined strong docking scores (−17.912 and −16.244 kcal mol⁻¹) with SAFC scores of 0.694 and 0.626. For this BRAF case study, favorable Vina scores and conventional drug-likeness metrics did not consistently correspond to high SAFC ranking scores, indicating that the computational criteria were not fully redundant. For exploratory MD follow-up, candidates were prioritized using a Pareto-style multi-criterion rule considering SAFC, docking score, QED, and SA. Candidates were favored when improvement in docking or SAFC did not require a marked deterioration in others. PFM and MolJO were selected because they yielded representative candidates in this balanced region, rather than because they achieved the highest SAFC scores overall. Representative high-priority compounds from PFM and MolJO were subsequently examined using 100-ns protein–ligand MD simulations (Fig. 8b,c). In both exploratory trajectories, the receptor backbone reached a comparatively stable global RMSD regime after initial relaxation; however, the MolJO complex exhibited greater backbone drift and more pronounced ligand rearrangement than the PFM complex. Cα RMSF profiles indicated that most receptor regions remained relatively constrained, with elevated fluctuations localized to discrete flexible segments. The ligand RMSD values nevertheless indicated substantial deviation from the initial docked poses, emphasizing that docking geometries should not be interpreted as rigid binding configurations. End-state MM/GBSA analysis over the sampled trajectory windows yielded mean interaction-energy estimates of −42.38 ± 4.12 kcal mol⁻¹ for the PFM complex and −38.94 ± 4.96 kcal mol⁻¹ for the MolJO complex, providing greater computational support for the PFM candidate under the evaluated conditions. These values are approximate comparative estimates rather than experimentally measured binding free energies. Collectively, the results show that SAFC adds a dynamics-aware prioritization layer that is partly orthogonal to QED, synthetic accessibility and docking score. Its principal value is therefore in narrowing generated libraries to candidates warranting more extensive free-energy calculations and experimental functional validation.

**Fig. 8.**
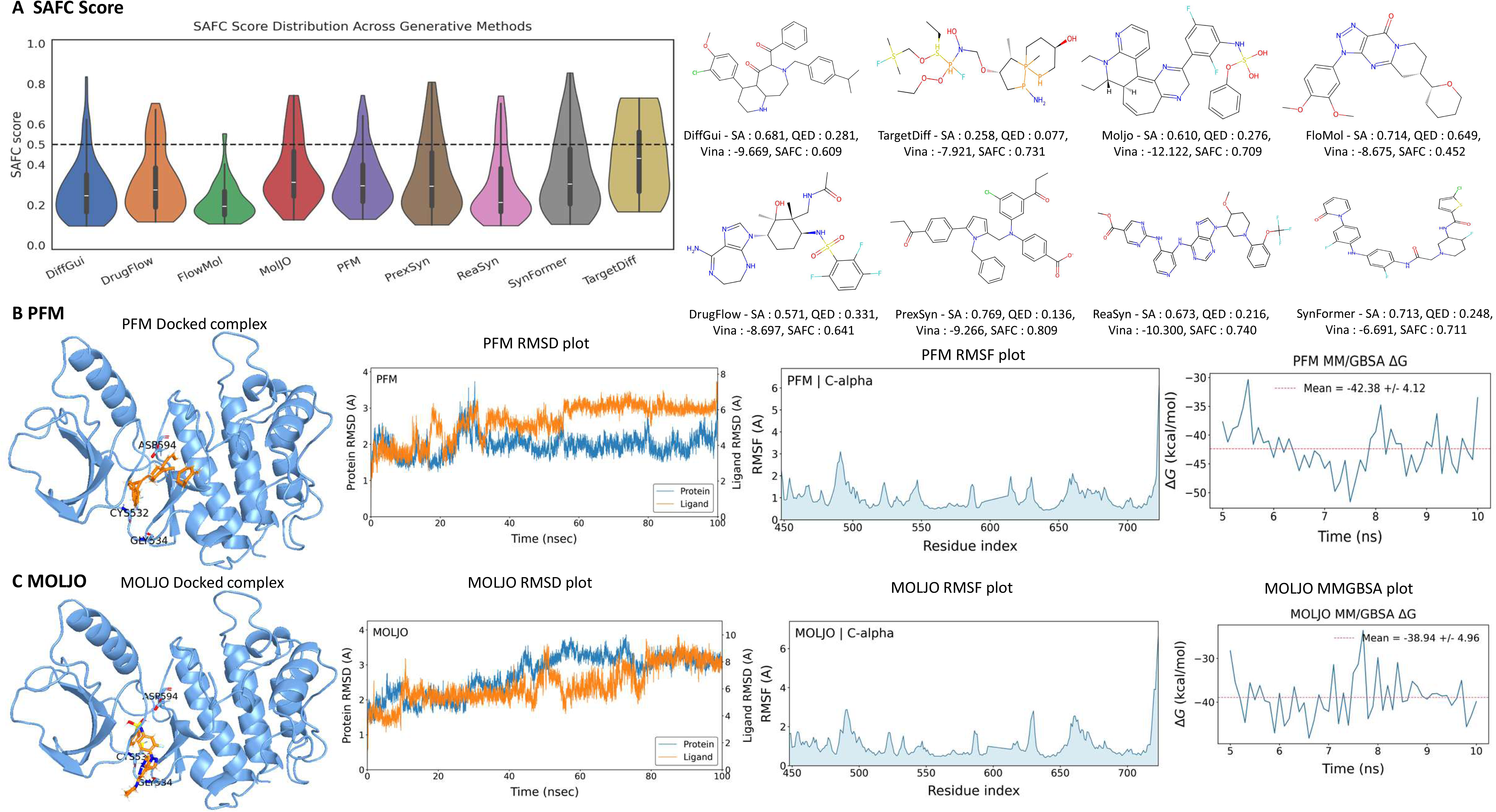
State-aware functional prioritization and dynamic assessment of selected molecules. **a,** Distribution of state-aware functional classifier (SAFC) scores across molecules generated by the 12 models. SAFC integrates receptor-state ensembles, docking-derived interaction features and protein–ligand graph representations to prioritize complexes displaying potentially relevant state-dependent interaction patterns. Violin widths indicate density, boxes represent the interquartile range and centre lines denote medians. The horizontal dashed line indicates the SAFC prioritization threshold of 0.5. Chemical structures show the highest-ranked candidate from each model, with normalized synthetic accessibility (SA), quantitative estimate of drug-likeness (QED), AutoDock Vina score and SAFC score indicated. **b,c,** Dynamic evaluation of representative high-priority candidates generated by PFM (b) and MolJO (c). From left to right, panels show the initial docked complex and selected interacting residues, protein-backbone and ligand root-mean-square deviation (RMSD) during 100-ns molecular dynamics simulations, Cα root-mean-square fluctuation (RMSF) by residue and MM/GBSA interaction-energy estimates over the displayed trajectory window. Mean MM/GBSA estimates were −42.38 ± 4.12 kcal mol⁻¹ for PFM and −38.94 ± 4.96 kcal mol⁻¹ for MolJO. SAFC and MM/GBSA values represent computational prioritization metrics rather than experimental evidence of functional activity or binding affinity.

### Evaluation of rentosertib supports clinically relevant candidate prioritization

As an external translational case study, we applied the complete prioritization workflow to rentosertib (ISM001-055/INS018_055), an AI-discovered and AI-designed inhibitor of TRAF2- and NCK-interacting kinase (TNIK) developed for idiopathic pulmonary fibrosis^36^. Rentosertib has demonstrated acceptable safety and tolerability in a randomized Phase IIa trial, together with an exploratory dose-dependent lung-function signal, and entered Phase III development in 2026^37^. Using the TNIK structure (PDB 8ZML), the pipeline assigned rentosertib favorable QED (0.76), normalized synthetic-accessibility (0.68) and Vina score of −9.91 kcal mol⁻¹ values, together with an SAFC score of 0.6949. Thus, the clinically advanced compound satisfied all predefined prioritization criteria and was recovered as a high-priority candidate by the integrated chemical-quality, binding and state-aware assessment. Although this retrospective single-compound analysis does not establish prospective predictive accuracy or independently validate clinical efficacy, it provides a clinically grounded positive-control example demonstrating that the framework can recognize a translationally advanced molecule without relying on any single molecular metric.

## Discussion

This study establishes a task-aware framework for comparing AI-based molecular generators across 176 protein–ligand systems, multiple conditioning paradigms and successive stages of computational candidate prioritization. Its principal advantage is that models were evaluated beyond chemical validity or docking performance alone. The benchmark jointly considered operational robustness, drug-likeness, synthetic accessibility, molecular and scaffold diversity, docking, ADME and toxicity liabilities, receptor-ensemble-aware prioritization was illustrated separately in the BRAF/3SKC case study, and computational efficiency was assessed from recorded runtime and hardware-use statistics. This design revealed that model performance is intrinsically multi-dimensional, methods that reliably generated large candidate sets were not necessarily those producing the most drug-like, diverse, synthetically accessible or binding-compatible molecules. Accordingly, the benchmark is intended not to identify a universally superior model, but to define where different model classes are most useful and how their outputs should be integrated into a practical discovery workflow. The first implication is that chemical validity is an insufficient measure of generative performance. Validity was nearly saturated among emitted molecules, whereas target completion and valid-output yield differed substantially across methods (Fig. 3). Some models completed nearly all targets and consistently returned close to the requested 100 valid molecules, whereas others frequently produced partial candidate sets despite high nominal completion. Operational robustness should therefore be reported using both target-level execution and usable molecular yield. The property analyses further identified a trade-off between chemical optimization and exploration (Fig. 4). Within the complete-case analysis FlowMol and Pocket2Mol provided comparatively favorable balances of drug-likeness and synthetic accessibility, whereas TargetDiff, PFM and ShEPhERD explored broader molecular or scaffold space but yielded fewer molecules satisfying the joint property criteria. These regimes may serve different purposes, broad exploration can support hit discovery and scaffold hopping, whereas higher property-pass rates may be preferable when synthesis and experimental testing capacity are limited.

Integration with docking, ADME and toxicity predictions demonstrated that advantages at one stage rarely persisted across the complete workflow. Favorable docking scores were not consistently accompanied by high QED, synthetic accessibility or joint pass rates, supporting the use of target-standardized Pareto analysis rather than ranking models by raw docking energy alone (Fig. 5). Similarly, molecules with favorable absorption-related predictions could retain liabilities in solubility, plasma-protein binding, cytochrome P450 inhibition or clearance (Fig. 6). Toxicity burdens were also model and endpoint dependent, and high drug-likeness or docking performance did not imply a low predicted liability burden (Fig. 7). Together, these findings argue against collapsing molecular quality into a single composite score without retaining endpoint-level information. A more defensible strategy is sequential prioritization, in which independent filters identify different failure modes while uncertainty and target-level variation remain visible. The exploratory BRAF/3SKC case study extends this staged prioritization framework by introducing receptor-ensemble-aware SAFC ranking after conventional molecular-property and docking analyses (Fig. 8). Static docking evaluates a molecule against a limited receptor configuration and may overlook changes in pocket geometry or interaction networks across conformational states. SAFC combines molecular-dynamics-derived receptor ensembles, ensemble docking and protein–ligand interaction representations to prioritize candidates displaying interaction patterns that persist across sampled states. The selected PFM and MolJO candidates retained plausible protein–ligand configurations during 100-ns simulations and showed favorable MM/GBSA estimates, providing dynamics-based consistency checks for the prioritization procedure. Rather than establishing binding, these analyses illustrate the value of extending molecular prioritization beyond static structures and isolated physicochemical metrics. The combined results demonstrate that chemical quality, docking performance, ADME behavior, toxicity liabilities and receptor-state-dependent interactions capture distinct and only partially overlapping dimensions of candidate suitability. Accordingly, generated molecules should progress through a staged, multi-objective workflow in which conventional molecular filters remove chemically unfavorable candidates, docking evaluates target compatibility, ADME and toxicity models identify developability liabilities, and dynamics-aware methods assess the persistence of protein–ligand interactions across receptor conformations. Within this framework, SAFC serves as classification layer for selecting functional candidates for more rigorous free-energy calculations and experimental testing.

The benchmark supports a practical model-selection framework based on the information available at the start of a project and the intended design objective (Fig. 9). When reliable binding-pocket coordinates are available, pocket-conditioned generators are appropriate for exploring receptor-complementary chemical space. When a known ligand provides three-dimensional shape, electrostatic or pharmacophore information, ligand-guided methods can support analogue generation and scaffold replacement. Reference-SMILES models are suited to chemical-series optimization, whereas reaction-aware generators are preferable when synthetic feasibility must be enforced during generation. Methods operating without target-specific structural information can provide broad chemical hypotheses but require subsequent target-specific docking and pose assessment. These model classes should be regarded as complementary components rather than interchangeable competitors. We therefore propose a hybrid workflow in which generation is followed by chemical-validity assessment, QED and synthetic-accessibility filtering, docking and pose inspection, ADME and toxicity profiling, receptor-state evaluation and synthesis-aware prioritization. Model choice should also reflect the discovery stage. Early hit discovery may tolerate lower predicted synthesizability in exchange for scaffold diversity, whereas lead optimization requires tighter control of potency, selectivity, metabolism, toxicity and route feasibility. The framework thus converts benchmark results into actionable guidance: select the generator according to input availability, retain sufficient chemical diversity during initial exploration and apply progressively more stringent, target-aware filters before experimental nomination.

**Fig. 9.**
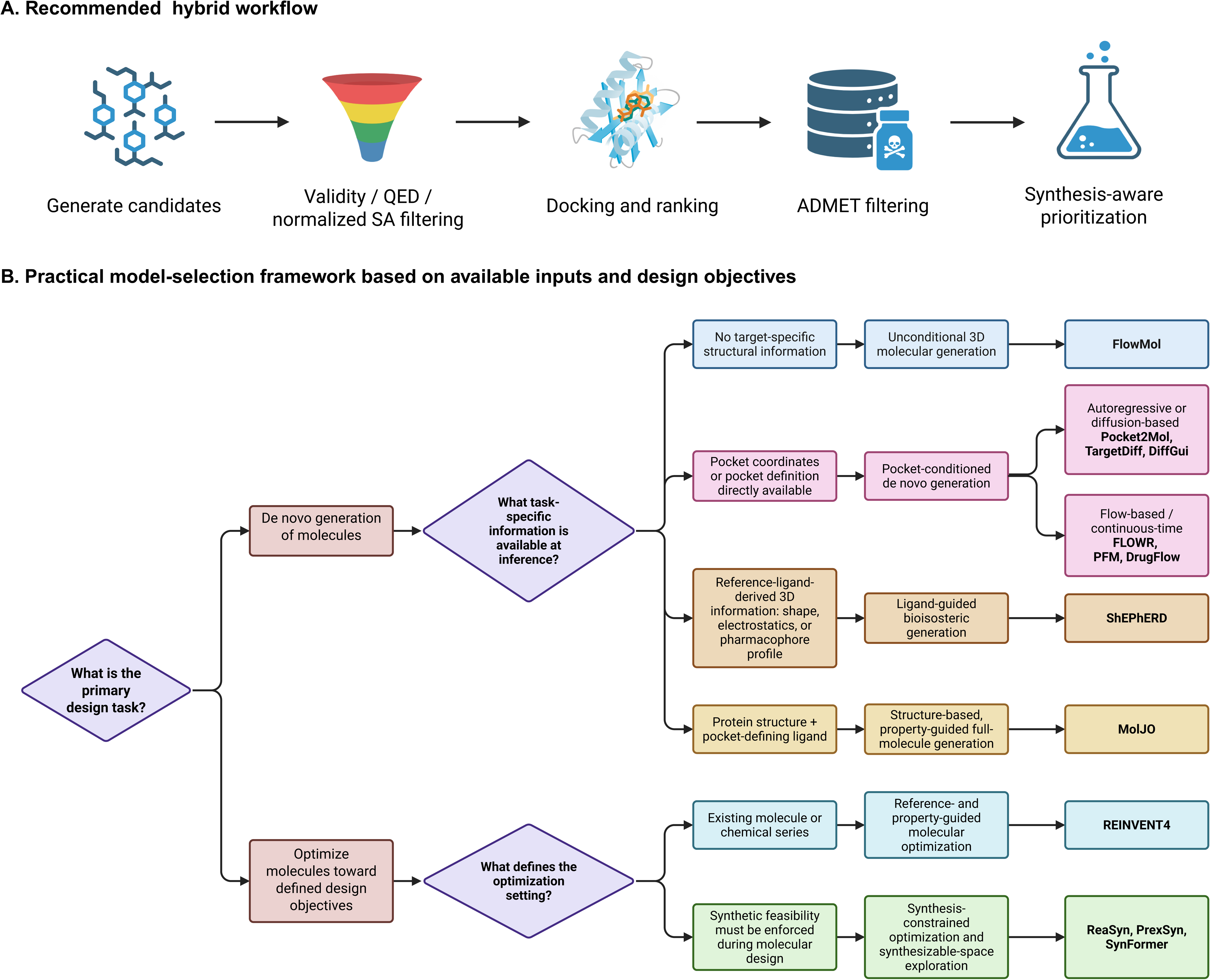
Recommended framework for selecting molecular generative models. **a,** Recommended hybrid workflow for prioritizing AI-generated molecules. Candidate generation is followed by chemical-validity assessment, QED and normalized synthetic-accessibility filtering, structure-based docking and ranking, ADMET evaluation and synthesis-aware prioritization. **b,** Practical model-selection decision tree based on the available input information and intended design objective. For de novo generation, unconditional molecular generation can be performed with FlowMol when target-specific structural information is unavailable. Pocket coordinates support pocket-conditioned methods, including Pocket2Mol, TargetDiff, DiffGui, FLOWR, PFM and DrugFlow, whereas reference-ligand shape, electrostatic or pharmacophore information supports ShEPhERD. Availability of both a protein structure and pocket-defining ligand supports structure- and property-guided generation with MolJO. For optimization tasks, REINVENT4 enables reference- or property-guided molecular refinement, whereas ReaSyn, PrexSyn and SynFormer prioritize synthetic feasibility and reaction-aware chemical-space exploration. The framework emphasizes selecting models according to input modality and discovery objective rather than relying on a single overall performance ranking.

Several challenges remain before this workflow can produce experimentally actionable candidates routinely (Fig. 10). First, chemically valid outputs may still contain ligand strain, unfavorable interactions or implausible pocket geometries. Generators will need more explicit stereochemical, conformational-energy, steric and pharmacophore constraints. Second, descriptor-based synthetic-accessibility scores do not capture whether a compound can be produced through a reliable and economical route. Integration with reaction templates, reagent availability, retrosynthetic confidence and route complexity is therefore needed. Third, most models treat receptors as static structures and incompletely represent induced fit, alternative protonation states, water-mediated interactions and allosteric conformations. Ensemble generation, enhanced sampling and state-aware scoring may improve performance for flexible targets, although they will increase computational cost.

**Fig. 10.**
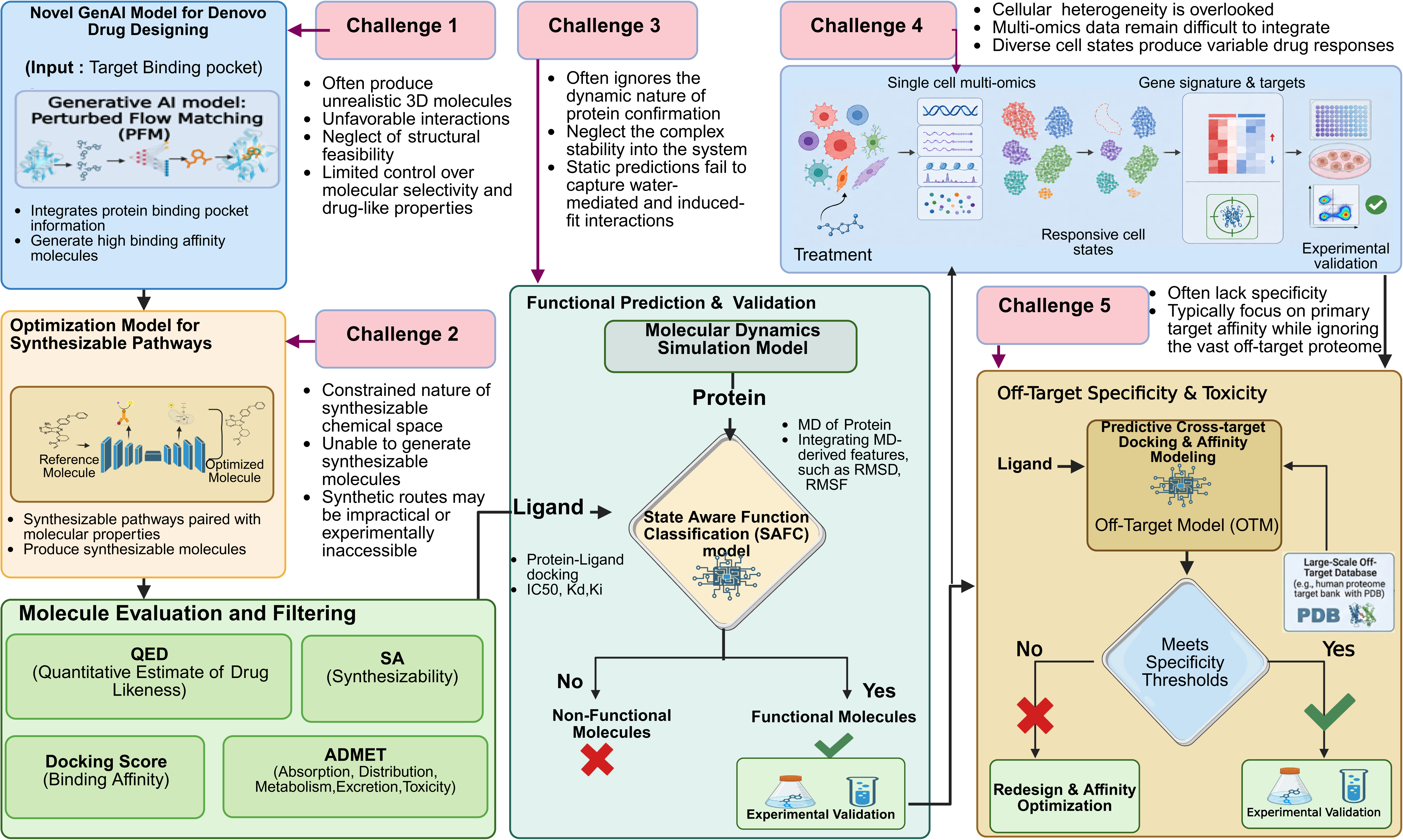
Key challenges in AI-enabled de novo drug design. Schematic representation of five major challenges limiting the translation of AI-generated molecules into experimentally validated drug candidates. Challenge 1, generative models can produce chemically or structurally unrealistic molecules, unfavorable protein–ligand interactions and insufficient control over molecular selectivity and drug-like properties. Challenge 2, synthesis-aware optimization remains restricted by limited accessible chemical space, uncertain route feasibility and experimentally inaccessible reaction proposals. Challenge 3, static structural evaluation may neglect receptor dynamics, complex stability and water-mediated or induced-fit interactions; molecular-dynamics-derived receptor ensembles and state-aware functional classification provide a route toward dynamics-informed prioritization. Challenge 4, cellular heterogeneity and incomplete integration of single-cell multi-omics can obscure cell-type- and cell-state-specific responses, gene signatures and drug targets, requiring perturbational profiling and experimental validation. Challenge 5, optimization against a primary target does not ensure proteome-wide selectivity or acceptable toxicity; cross-target docking, off-target modeling and large-scale target databases can support specificity filtering before experimental testing. The proposed workflow integrates chemical-property assessment, docking, ADMET prediction, dynamics-aware functional prioritization, single-cell response analysis and off-target evaluation, with iterative redesign and experimental validation at critical decision points.

A further challenge is connecting molecular design to biological context. Protein– ligand complementarity alone cannot determine whether target engagement will produce the intended response in disease-relevant cell types or states, particularly when target expression, pathway activity, chromatin accessibility, cellular composition and microenvironmental signaling vary across patients. Integrating generated-molecule representations with perturbational single-cell and spatial multi-omics could provide a functional layer for predicting which cell populations respond to treatment, the magnitude and direction of response, and whether exposure induces therapeutic, resistant or off-target cell-state transitions. In such a framework, chemical embeddings could be combined with baseline transcriptomic, epigenomic and spatial profiles from patient-derived organoids, tissue models or biopsies to predict post-treatment molecular states and gene-expression changes. Perturbation-prediction approaches—including conditional autoencoder, transformer, optimal-transport and graph-based models— could then infer drug-response signatures, context-specific sensitivity and resistance, and candidate mechanisms of action. Mechanistic inference would require linking predicted target engagement to downstream pathway modulation, transcription-factor activity, gene-regulatory-network rewiring and phenotypic changes across responsive and non-responsive cell populations. Comparison with genetic perturbations of the intended target, together with dose- and time-resolved molecular profiles, could help distinguish target-dependent mechanisms from compensatory responses and nonspecific cellular stress. Analysis of differentially expressed genes, pathway activity and regulatory programs could consequently identify which molecular processes are activated or suppressed by each candidate and generate experimentally testable mechanism-of-action hypotheses. Patient-derived organoids and spatially resolved tissue models would additionally preserve clonal heterogeneity, cell–cell communication and microenvironmental organization that are absent from conventional monocultures, enabling computational predictions to be evaluated through target-engagement assays, single-cell transcriptomics, viability measurements, imaging and functional phenotyping. Ultimately, linking chemical design to cell-type-specific response and mechanism-of-action signatures could support precision-medicine strategies in which candidates are matched to patients whose tumors contain susceptible molecular and cellular states.

However, predicting a desirable response in a specific cellular context does not ensure molecular target selectivity. A complementary challenge is therefore to determine whether generated compounds preferentially engage the intended target rather than homologous proteins or unrelated proteome targets. Optimization against a single binding pocket can overlook such cross-target interactions and produce candidates with unintended pharmacology or toxicity. Proteome-scale structural screening, cross-target docking, chemoproteomic profiling and experimentally designed counter-screens should therefore be incorporated earlier in lead optimization. Integrating these molecular-specificity assessments with single-cell perturbation profiles could distinguish on-target therapeutic responses from off-target or cell-state-specific adverse effects, providing a more complete basis for functional and translational prioritization.

This study has several limitations. The target set was diverse but unevenly distributed across protein classes, with fewer transporters, transcription regulators and other structurally challenging targets. Cross-workflow summaries may consequently be influenced disproportionately by the better-represented protein families. The evaluated models also differed in their conditioning information and intended application; pocket-conditioned generators, reference-ligand methods, SMILES optimizers and synthesis-aware models cannot be considered equivalent even when their outputs undergo the same downstream evaluation. Complete-case analyses were restricted to 54 targets for molecular quality and diversity and 65 targets for integrated docking, improving matched comparability but excluding targets for which one or more workflows failed to provide sufficient outputs and thereby introducing potential selection bias toward easier systems. All prioritization endpoints were computational. AuroDock Vina scores, MM/GBSA estimates, QED, synthetic-accessibility scores and predicted ADME or toxicity endpoints are imperfect surrogates for binding affinity, efficacy, pharmacokinetics, safety and experimental synthesizability. Receptor flexibility, water networks, protonation and tautomeric uncertainty, induced fit and allosteric behavior were not modeled systematically for every molecule. Computational scalability measurements are also dependent on hardware, software versions, batching and implementation efficiency and should therefore be interpreted under the tested conditions. Finally, the study did not include prospective compound synthesis or experimental testing, preventing direct assessment of whether differences in benchmark performance translate into improved discovery success. Future benchmarks should expand target coverage, incorporate prospective and blind evaluation, and connect computational prioritization to compound synthesis, biophysical binding measurements, cellular target engagement and functional phenotyping. Standardized reporting should document model-conditioning inputs, attempted and completed targets, requested and obtained molecule counts, failed-output handling, validity and diversity metrics, property thresholds, docking protocols, ADME and toxicity applicability domains, and computational resource use.

The principal contribution of this study is therefore not a single model ranking, but an end-to-end benchmark and practical decision framework that treats AI-enabled molecular design as a multi-stage, multi-objective preclinical process. Unlike benchmarks focused predominantly on validity, docking or descriptor-based quality, our framework jointly evaluates operational robustness, chemical and scaffold diversity, drug-likeness, synthesizability, structural compatibility, predicted ADME and toxicity liabilities, and computational efficiency. It further introduces SAFC as an exploratory dynamics-aware layer for prioritizing candidates whose receptor-state-dependent interaction patterns may not be captured by static docking or conventional molecular descriptors alone. The BRAF case study and retrospective evaluation of rentosertib illustrate how chemical quality, docking, receptor dynamics and state-aware functional evidence can be integrated without interpreting any single computational score as proof of biological activity. No model was uniformly favorable across all evaluation dimensions; effective deployment therefore requires selecting models according to available inputs and project objectives and combining complementary generators with transparent downstream prioritization and experimental validation. By linking model selection to available inputs, design objectives and downstream validation requirements, the proposed framework offers actionable guidance while highlighting the chemical, structural, functional, cellular and translational barriers that must be addressed before AI-generated molecules can advance reliably through preclinical development. Together, this framework provides a reproducible foundation for moving beyond generator-centric comparison toward prospective assessment of whether AI-designed molecules are synthesizable, target-engaging, functionally active and therapeutically relevant.

## Methodology

### Benchmarking Dataset Construction

The benchmarking dataset comprised 176 experimentally resolved protein–ligand complexes assembled from three complementary structural resources: AlignDockBench (61 systems), PoseBusters (40 systems) and PLINDER (75 systems). AlignDockBench served as the primary cohort. Its targets were identified using a curated annotation table containing PDB identifiers, structural classifications, experimental resolutions and source organisms. When an AlignDockBench target contained multiple candidate complexes, one complex was selected using a deterministic procedure, all selections were recorded in a target-level manifest. The PoseBusters cohort was derived from the 308-system journal subset. Structures already represented in AlignDockBench were excluded, and ligand, binding-pocket and receptor descriptors were calculated for the remaining systems. Receptor sequences were clustered using MMseqs2, and 40 systems were selected to maximize structural and chemical diversity. The PLINDER cohort was sampled from the official test split after retaining quality-controlled, single-ligand systems and applying ligand-molecular-weight and pocket-size filters. PDB entries present in either AlignDockBench or the selected PoseBusters cohort were removed. Seventy-five PLINDER systems were then sampled across sequence and structural clusters, ligand-uniqueness categories, size strata and the availability of linked apo or predicted structures. External-cohort sampling used a fixed random seed of 2026. Cohort and target identifiers were retained throughout the analysis to prevent molecule-level observations from different systems being treated as independent replicates.

### Dataset integration and pairing strategy

Protein–ligand inputs were organized using native and cross-target pairing schemes. In the native scheme, each crystallographic ligand was retained with its experimentally resolved receptor complex, providing the reference pocket and ligand pose for target-specific generation and evaluation. In the cross-target scheme, selected ligands were evaluated against additional receptors using a standardized docking workflow to examine transferability and potential target specificity. Model inputs were prepared according to their intended conditioning modality: receptor structures or extracted binding pockets were supplied to pocket-conditioned generators; three-dimensional reference ligands were used for shape-, electrostatic- or pharmacophore-conditioned methods; and reference SMILES or reaction information was supplied to ligand-optimization and synthesis-aware models. Regardless of input modality, generated molecules were processed through the same downstream validity, property, docking and ADMET evaluation pipeline. Native target–ligand associations and model-specific input provenance were retained throughout the analysis.

### Structural and chemical quality control

Protein–ligand systems were inspected before model preparation to confirm the availability of a resolved receptor structure, an identifiable bound ligand and sufficient binding-site coordinates for pocket definition. Systems with missing or unreadable structural files, absent ligands, or binding sites that could not be consistently extracted were excluded. Protein identifiers, chain assignments and ligand records were harmonized across cohorts, and duplicate PDB entries were removed during dataset assembly. Ligand structures were parsed with RDKit and screened for chemical validity, including sanitization and valence consistency. Molecules that could not be parsed or sanitized were excluded, and valid structures were represented using canonical SMILES for molecule-level tracking. Standardized protein and ligand records were subsequently converted into the model-specific formats required by each method, including PDB or pocket-coordinate files for receptors and SMILES, SDF or three-dimensional reference-ligand files for molecular inputs. Unique target and molecule identifiers were preserved throughout the workflow to link generated outputs to their source systems and downstream evaluations. An overview of dataset preparation and benchmarking is provided in Fig. 1.

### Target completion and output robustness

Target completion was defined as the generation of at least one molecular output for a benchmark target, irrespective of its subsequent chemical validity. Output robustness was assessed using the number of RDKit-valid molecules returned relative to the requested maximum of 100 molecules per target. For each model, quota-attainment rates were calculated as the proportion of all 176 targets yielding at least 25, 50, 75, 90 or 100 valid molecules. Target-level outcomes were further classified as near-complete output (≥90 valid molecules), partial output (1–89 valid molecules), output with no valid molecules, or no output. Targets that failed during generation, postprocessing or validity assessment were retained in the denominator, such that the reported rates measured end-to-end operational performance rather than performance conditional on successful execution.

### Drug-likeness, synthetic accessibility and chemical diversity

Quantitative estimate of drug-likeness (QED) and synthetic-accessibility (SA) scores were calculated for every RDKit-valid molecule before docking-based selection. Because lower raw SA scores indicate greater predicted synthetic accessibility, raw SA scores were transformed using SA_norm = (10 − SA_raw)/9, where values approaching 1 indicate easier predicted synthesis. Molecules satisfying QED ≥ 0.70 and normalized SA ≥ 0.667—equivalent to a raw SA score ≤ 4.0—were classified as passing the joint drug-likeness and synthetic-accessibility criterion. Before diversity analysis, molecules were standardized by retaining the largest covalent fragment and generating canonical isomeric SMILES with RDKit. Chemical diversity was defined as one minus the mean pairwise Tanimoto similarity calculated from chirality-aware Morgan fingerprints with a radius of 2 and 2,048 bits. Bemis–Murcko scaffolds were extracted from the standardized molecules, and scaffold diversity was quantified using normalized Shannon entropy. To limit bias arising from unequal output sizes, molecular and scaffold diversity were estimated using repeated equal-size subsampling within each model–target combination. The primary complete-case comparison included 54 targets for which all 12 models generated enough valid molecules for matched analysis.

### Integrated docking and multi-objective prioritization

Docking performance was evaluated on the 65 protein targets for which all 12 models produced comparable valid molecular outputs. Generated molecules were docked against their corresponding target structures using a common AutoDock Vina protocol, and the best-scoring pose for each molecule was retained. Because raw Vina scores are strongly influenced by pocket size, composition and ligand properties, scores were normalized within each target before comparisons among models. For each molecule, target-normalized docking performance was calculated as:

Normalized docking performance = target median Vina score − molecule Vina score

Thus, a value of zero represents the median docking performance for that target, positive values indicate more favorable docking than the target median, and negative values indicate less favorable docking. This transformation reverses the direction of the original Vina scale—on which more negative values indicate stronger predicted binding—while preserving the score difference in kcal mol⁻¹. Model-level distributions were generated from the median normalized performance obtained for each model– target combination. Within each target, models were also ranked according to their median raw Vina score, with rank 1 assigned to the most favorable median score. Docking performance was integrated with QED and normalized SA using predefined thresholds of Vina score ≤ −7.0 kcal mol⁻¹, QED ≥ 0.70 and normalized SA ≥ 0.667. For each model and target, the proportions of molecules passing the individual criteria and the combined QED–SA and QED–SA–Vina criteria were calculated. Target-level percentages were subsequently summarized using the median across the 65 common targets so that each target contributed equally, irrespective of the number of molecules generated. Multi-objective performance was additionally assessed using Pareto analysis. QED and normalized SA were maximized, whereas the raw Vina score was minimized. A molecule was considered Pareto optimal when no other molecule for the same target performed at least as well for all three objectives and strictly better for at least one objective. Pareto yield was calculated as the percentage of valid molecules belonging to the target-specific Pareto set and was summarized across targets for each model. Target coverage was defined as the proportion of the 65 targets for which a model generated at least one molecule simultaneously satisfying QED ≥ 0.70, normalized SA ≥ 0.667 and Vina ≤ −7.0 kcal mol⁻¹.

### State-aware functional prioritization using SAFC

To further evaluate whether AI-generated molecules were likely to retain functional relevance under dynamic receptor conditions, we developed an in-house state-aware functional classifier (SAFC) as a downstream prioritization module (Supplementary Note S18). The design of SAFC was conceptually inspired by dynamics-informed ligand efficacy modeling, such as Dynamic-GLEP, but was independently implemented as a task-specific workflow for functional prioritization of AI-generated molecules. SAFC integrates state-defined receptor structures, molecular dynamics (MD)-derived receptor ensembles, representative-state selection, ensemble docking, complex-level graph construction, and EquiScore-inspired transfer learning. In this framework, receptor conformations are sampled from MD trajectories and filtered to preserve local pocket diversity and state-sensitive structural descriptors. Each generated ligand is docked against representative receptor conformations, and the resulting protein–ligand complexes are converted into heterogeneous interaction graphs containing ligand chemistry, pocket–ligand contacts, geometric descriptors, and docking-derived features. The graph-based model maps each protein–ligand complex into a learned embedding, which is aggregated across receptor states to produce a ligand-level prediction score. The resulting SAFC score ranges from 0 to 1 and is interpreted as a predicted probability that a generated molecule exhibits functionally relevant interaction patterns under dynamic receptor-state conditions. In this benchmarking study, SAFC was used as a computational downstream prioritization layer rather than as direct experimental evidence of biological activity. As illustrated in Supplementary Fig.S37, the SAFC workflow combines MD-derived receptor ensembles, representative-state selection, ensemble docking, and protein–ligand interaction-graph construction before graph-based functional scoring. The figure also highlights the use of EquiScore-inspired transfer learning as an interim classification module for separating complexes with predicted functional relevance from non-functional or uncertain interaction states.

The BRAF-specific receptor ensemble used for SAFC was constructed from three experimentally determined inhibitor-bound crystal structures selected to represent complementary kinase conformations 2FB8, representing a Type I DFG-in/αC-in state 1UWH, representing a Type II DFG-out/αC-in state, and 3SKC, representing a Type I.5 DFG-in/αC-out state^45–47^. Each structure was used as the starting receptor for the corresponding state-specific molecular-dynamics and representative-conformation selection procedure. Molecules generated for the 3SKC benchmark system were subsequently evaluated against representative receptor conformations derived from all three state-specific ensembles rather than against 3SKC alone. The resulting receptor-conditioned protein–ligand representations were integrated by the independently developed BRAF-specific SAFC model to generate ligand-level within-target ranking scores. This three-state design incorporated complementary DFG and αC-helix arrangements relevant to BRAF inhibitor recognition without if the three structures exhaust all theoretically possible kinase conformations.

## Data and code availability

The datasets and codes are available at GitHub (https://github.com/HimansuKumarIIITA/DrugDesignAI-Benchmark).

## Acknowledgements

This work was partially supported by the National Institutes of Health [R01LM014156, R01GM153822, R01CA241930, R01AA032723-01 to X.Z], the National Science Foundation [2217515, 2326879 to X.Z], the Cancer Prevention and Research Institute of Texas [RP250043 to X.Z], The funders had no role in study design, data collection and analysis, decision to publish or preparation of the manuscript.

## Supplementary figure legend

**Supplementary Fig. 1. QED, synthetic accessibility and AutoDock Vina scores of PFM-generated molecules. a,** Distribution of quantitative estimate of drug-likeness (QED) values; the dashed line denotes the QED threshold of 0.70. Among 16,235 molecules, 1,926 met this criterion (median QED, 0.49). **b,** Distribution of normalized synthetic-accessibility (SA) desirability scores, for which higher values indicate greater predicted accessibility; 2,836 molecules met the threshold of 0.667 (median, 0.53). **c,** Distribution of AutoDock Vina scores; 13,109 molecules achieved scores ≤−7 kcal mol⁻¹ (median, −10.17 kcal mol⁻¹). **d,** Joint QED–SA distribution colored by Vina score. In total, 393 molecules simultaneously satisfied the QED, SA and docking thresholds.

**Supplementary Fig. 2. QED, synthetic accessibility and AutoDock Vina scores of MolJO-generated molecules. a,** QED distribution, with the threshold of 0.70 indicated by the dashed line; 3,381 of 16,706 molecules passed this criterion (median, 0.54). **b,** Distribution of normalized SA desirability scores; 10,721 molecules met the threshold of 0.667 (median, 0.71). **c,** AutoDock Vina score distribution; 13,181 molecules achieved scores ≤−7 kcal mol⁻¹ (median, −9.60 kcal mol⁻¹). **d,** Joint QED–SA distribution colored by Vina score, identifying 1,883 molecules that satisfied all three thresholds.

**Supplementary Fig. 3. QED, synthetic accessibility and AutoDock Vina scores of DrugFlow-generated molecules. a,** QED distribution, with the threshold of 0.70 indicated by the dashed line; 2,897 of 12,018 molecules passed this criterion (median, 0.53). **b,** Distribution of normalized SA desirability scores; 7,226 molecules met the threshold of 0.667 (median, 0.70). **c,** AutoDock Vina score distribution; 9,058 molecules achieved scores ≤−7 kcal mol⁻¹ (median, −8.67 kcal mol⁻¹). **d,** Joint QED–SA distribution colored by Vina score. A total of 1,614 molecules simultaneously met the QED, SA and docking criteria.

**Supplementary Fig. 4. QED, synthetic accessibility and AutoDock Vina scores of FlowMol-generated molecules. a,** QED distribution; 10,808 of 17,183 molecules met the threshold of 0.70 (median, 0.75). **b,** Distribution of normalized SA desirability scores; 16,809 molecules met the threshold of 0.667 (median, 0.86). **c,** AutoDock Vina score distribution; 9,805 molecules achieved scores ≤−7 kcal mol⁻¹ (median, −7.25 kcal mol⁻¹). **d,** Joint QED–SA distribution colored by Vina score, identifying 5,968 molecules satisfying all three criteria.

**Supplementary Fig. 5. QED, synthetic accessibility and AutoDock Vina scores of Pocket2Mol-generated molecules. a,** QED distribution; 6,696 of 17,296 molecules met the threshold of 0.70 (median, 0.65). **b,** Distribution of normalized SA desirability scores; 13,703 molecules met the threshold of 0.667 (median, 0.78). **c,** AutoDock Vina score distribution; 10,441 molecules achieved scores ≤−7 kcal mol⁻¹ (median, −7.50 kcal mol⁻¹). **d,** Joint QED–SA distribution colored by Vina score. In total, 4,458 molecules simultaneously satisfied the QED, SA and docking thresholds.

**Supplementary Fig. 6. QED, synthetic accessibility and AutoDock Vina scores of ShEPhERD-generated molecules. a,** QED distribution; 3,984 of 16,107 molecules met the threshold of 0.70 (median, 0.57). **b,** Distribution of normalized SA desirability scores; 6,077 molecules met the threshold of 0.667 (median, 0.62). **c,** AutoDock Vina score distribution; 11,861 molecules achieved scores ≤−7 kcal mol⁻¹ (median, −8.47 kcal mol⁻¹). **d,** Joint QED–SA distribution colored by Vina score, identifying 1,637 molecules that passed all three thresholds.

**Supplementary Fig. 7. QED, synthetic accessibility and AutoDock Vina scores of REINVENT4-generated molecules. a,** QED distribution; 2,692 of 10,866 molecules met the threshold of 0.70 (median, 0.55). **b,** Distribution of normalized SA desirability scores; 9,014 molecules met the threshold of 0.667 (median, 0.77). **c,** AutoDock Vina score distribution; 7,886 molecules achieved scores ≤−7 kcal mol⁻¹ (median, −8.22 kcal mol⁻¹). **d,** Joint QED–SA distribution colored by Vina score. A total of 1,900 molecules simultaneously satisfied all three criteria.

**Supplementary Fig. 8. QED, synthetic accessibility and AutoDock Vina scores of ReaSyn-generated molecules. a,** QED distribution; 1,201 of 8,048 molecules met the threshold of 0.70 (median, 0.31). **b,** Distribution of normalized SA desirability scores; 5,650 molecules met the threshold of 0.667 (median, 0.73). **c,** AutoDock Vina score distribution; 5,268 molecules achieved scores ≤−7 kcal mol⁻¹ (median, −7.79 kcal mol⁻¹). **d,** Joint QED–SA distribution colored by Vina score, identifying 697 molecules satisfying all three thresholds.

**Supplementary Fig. 9. QED, synthetic accessibility and AutoDock Vina scores of SynFormer-generated molecules. a,** QED distribution; 1,909 of 14,661 molecules met the threshold of 0.70 (median, 0.33). **b,** Distribution of normalized SA desirability scores; 8,667 molecules met the threshold of 0.667 (median, 0.70). **c,** AutoDock Vina score distribution; 9,489 molecules achieved scores ≤−7 kcal mol⁻¹ (median, −7.76 kcal mol⁻¹). **d,** Joint QED–SA distribution colored by Vina score. In total, 1,297 molecules passed the QED, SA and docking criteria.

**Supplementary Fig. 10. QED, synthetic accessibility and AutoDock Vina scores of PrexSyn-generated molecules. a,** QED distribution; 1,632 of 13,910 molecules met the threshold of 0.70 (median, 0.36). **b,** Distribution of normalized SA desirability scores; 8,838 molecules met the threshold of 0.667 (median, 0.72). **c,** AutoDock Vina score distribution; 9,739 molecules achieved scores ≤−7 kcal mol⁻¹ (median, −8.09 kcal mol⁻¹). **d,** Joint QED–SA distribution colored by Vina score, identifying 1,005 molecules that simultaneously satisfied all three thresholds.

**Supplementary Fig. 11. QED, synthetic accessibility and AutoDock Vina scores of TargetDiff-generated molecules. a,** QED distribution; 2,344 of 14,872 molecules met the threshold of 0.70 (median, 0.46). **b,** Distribution of normalized SA desirability scores; 3,454 molecules met the threshold of 0.667 (median, 0.57). **c,** AutoDock Vina score distribution; 10,804 molecules achieved scores ≤−7 kcal mol⁻¹ (median, −8.49 kcal mol⁻¹). **d,** Joint QED–SA distribution colored by Vina score. A total of 605 molecules simultaneously met the QED, SA and docking criteria.

**Supplementary Fig. 12. QED, synthetic accessibility and AutoDock Vina scores of FLOWR-generated molecules. a,** QED distribution; 3,563 of 16,047 molecules met the threshold of 0.70 (median, 0.51). **b,** Distribution of normalized SA desirability scores; 9,450 molecules met the threshold of 0.667 (median, 0.70). **c,** AutoDock Vina score distribution; 8,995 molecules achieved scores ≤−7 kcal mol⁻¹ (median, −7.57 kcal mol⁻¹). **d,** Joint QED–SA distribution colored by Vina score, identifying 1,792 molecules satisfying all three criteria.

**Supplementary Fig. 13. ADME profiles of PFM-generated molecules. a,** Proportions of molecules predicted positive for selected absorption, distribution and metabolic endpoints, including human intestinal absorption (HIA), bioavailability, P-glycoprotein inhibition, blood–brain barrier penetration and inhibition of major cytochrome P450 (CYP) enzymes. **b,** Distributions of predicted aqueous solubility, Caco-2 permeability, plasma protein binding, volume of distribution, hepatocyte clearance and half-life; dashed lines indicate medians. **c,** Proportions satisfying the Lipinski rule-of-five and the predefined absorption, distribution, metabolism, excretion and overall ADME criteria. **d,** Per-molecule metabolic and physicochemical burden, represented by the numbers of major CYP enzymes predicted to be inhibited and Lipinski violations. The analysis included 9,449 molecular records representing 9,377 unique SMILES across 97 common benchmark targets. Classification endpoints used a probability threshold of 0.5; distribution and excretion pass ranges were defined using pooled 5th–95th percentile intervals.

**Supplementary Fig. 14. ADME profiles of MolJO-generated molecules. a,** Proportions of molecules predicted positive for selected absorption, distribution and metabolic endpoints, including HIA, bioavailability, P-glycoprotein inhibition, blood–brain barrier penetration and inhibition of major CYP enzymes. **b,** Distributions of predicted aqueous solubility, Caco-2 permeability, plasma protein binding, volume of distribution, hepatocyte clearance and half-life; dashed lines indicate medians. **c,** Proportions satisfying the Lipinski rule-of-five and the predefined absorption, distribution, metabolism, excretion and overall ADME criteria. **d,** Per-molecule metabolic and physicochemical burden, represented by the numbers of major CYP enzymes predicted to be inhibited and Lipinski violations. The analysis included 9,689 molecular records representing 9,256 unique SMILES across 97 common benchmark targets. Classification endpoints used a probability threshold of 0.5; distribution and excretion pass ranges were defined using pooled 5th–95th percentile intervals.

**Supplementary Fig. 15. ADME profiles of DrugFlow-generated molecules. a,** Proportions of molecules predicted to satisfy selected absorption, distribution and metabolic endpoints, including human intestinal absorption (HIA), bioavailability, P-glycoprotein (P-gp) inhibition, blood–brain barrier (BBB) penetration and inhibition of major cytochrome P450 (CYP) enzymes. **b,** Distributions of predicted aqueous solubility, Caco-2 permeability, plasma protein binding, volume of distribution, hepatocyte clearance and half-life. Dashed lines indicate distribution medians. **c,** Proportions satisfying the Lipinski rule-of-five and the predefined absorption, distribution, metabolism, excretion and overall ADME criteria. **d,** Per-molecule burden represented by the number of major CYP enzymes predicted to be inhibited and the number of Lipinski violations. The analysis included 10,641 molecular records representing 10,488 unique SMILES across 97 common benchmark targets. Classification endpoints used a probability threshold of 0.5; distribution and excretion pass ranges were defined using pooled 5th–95th percentile intervals.

**Supplementary Fig. 16. ADME profiles of FlowMol-generated molecules. a,** Predicted prevalence of favorable absorption and distribution properties and inhibition of major CYP enzymes. **b,** Distributions of aqueous solubility, Caco-2 permeability, plasma protein binding, volume of distribution, hepatocyte clearance and half-life; dashed lines denote medians. **c,** Percentages of molecules satisfying the Lipinski rule-of-five and the absorption, distribution, metabolism, excretion and overall ADME criteria. **d,** Numbers of major CYP enzymes predicted to be inhibited and Lipinski violations per molecule. The analysis included 9,699 molecular records representing 9,653 unique SMILES across 97 common benchmark targets. Classification endpoints used a probability threshold of 0.5; distribution and excretion pass ranges were defined using pooled 5th–95th percentile intervals.

**Supplementary Fig. 17. ADME profiles of Pocket2Mol-generated molecules. a,** Predicted prevalence of favorable absorption and distribution properties and inhibition of major CYP enzymes. **b,** Distributions of six continuous ADME properties, with dashed lines indicating medians. **c,** Percentages satisfying the Lipinski rule-of-five and the absorption, distribution, metabolism, excretion and overall ADME criteria. **d,** Per-molecule distributions of the number of inhibited major CYP enzymes and Lipinski violations. The analysis included 10,749 molecular records representing 10,385 unique SMILES across 97 common benchmark targets. Classification endpoints used a probability threshold of 0.5; distribution and excretion pass ranges were defined using pooled 5th–95th percentile intervals.

**Supplementary Fig. 18. ADME profiles of ShEPhERD-generated molecules. a,** Predicted prevalence of categorical absorption, distribution and CYP-inhibition endpoints. **b,** Distributions of aqueous solubility, Caco-2 permeability, plasma protein binding, volume of distribution, hepatocyte clearance and half-life; dashed lines indicate medians. **c,** Proportions satisfying the Lipinski rule-of-five and domain-specific and overall ADME criteria. **d,** Per-molecule metabolic and physicochemical burden, quantified by the numbers of inhibited major CYP enzymes and Lipinski violations. The analysis included 9,609 molecular records representing 9,608 unique SMILES across 97 common benchmark targets. Classification endpoints used a probability threshold of 0.5; distribution and excretion pass ranges were defined using pooled 5th–95th percentile intervals.

**Supplementary Fig. 19. ADME profiles of REINVENT4-generated molecules. a,** Predicted prevalence of categorical absorption, distribution and CYP-inhibition endpoints. **b,** Distributions of six continuous ADME properties, with dashed lines indicating medians. **c,** Proportions satisfying the Lipinski rule-of-five and the absorption, distribution, metabolism, excretion and overall ADME criteria. **d,** Numbers of major CYP enzymes predicted to be inhibited and Lipinski violations per molecule. The analysis included 9,499 molecular records representing 9,185 unique SMILES across 97 common benchmark targets. Classification endpoints used a probability threshold of 0.5; distribution and excretion pass ranges were defined using pooled 5th–95th percentile intervals.

**Supplementary Fig. 20. ADME profiles of ReaSyn-generated molecules. a,** Predicted prevalence of categorical absorption, distribution and CYP-inhibition endpoints. **b,** Distributions of aqueous solubility, Caco-2 permeability, plasma protein binding, volume of distribution, hepatocyte clearance and half-life; dashed lines indicate medians. **c,** Percentages satisfying the Lipinski rule-of-five and domain-specific and overall ADME criteria. **d,** Per-molecule distributions of predicted major-CYP inhibition and Lipinski violations. The analysis included 8,526 molecular records representing 8,453 unique SMILES across 97 common benchmark targets. Classification endpoints used a probability threshold of 0.5; distribution and excretion pass ranges were defined using pooled 5th–95th percentile intervals.

**Supplementary Fig. 21. ADME profiles of SynFormer-generated molecules. a,** Predicted prevalence of categorical absorption, distribution and CYP-inhibition endpoints. **b,** Distributions of six continuous ADME properties, with dashed lines denoting medians. **c,** Percentages satisfying the Lipinski rule-of-five and the absorption, distribution, metabolism, excretion and overall ADME criteria. **d,** Numbers of major CYP enzymes predicted to be inhibited and Lipinski violations per molecule. The analysis included 9,105 molecular records representing 9,028 unique SMILES across 97 common benchmark targets. Classification endpoints used a probability threshold of 0.5; distribution and excretion pass ranges were defined using pooled 5th–95th percentile intervals.

**Supplementary Fig. 22. ADME profiles of PrexSyn-generated molecules. a,** Predicted prevalence of categorical absorption, distribution and CYP-inhibition endpoints. **b,** Distributions of aqueous solubility, Caco-2 permeability, plasma protein binding, volume of distribution, hepatocyte clearance and half-life; dashed lines indicate medians. **c,** Proportions satisfying the Lipinski rule-of-five and domain-specific and overall ADME criteria. **d,** Per-molecule metabolic and physicochemical burden quantified by the numbers of inhibited major CYP enzymes and Lipinski violations. The analysis included 9,065 molecular records representing 9,006 unique SMILES across 97 common benchmark targets. Classification endpoints used a probability threshold of 0.5; distribution and excretion pass ranges were defined using pooled 5th–95th percentile intervals.

**Supplementary Fig. 23. ADME profiles of TargetDiff-generated molecules. a,** Predicted prevalence of categorical absorption, distribution and CYP-inhibition endpoints. **b,** Distributions of six continuous ADME properties, with dashed lines indicating medians. **c,** Percentages satisfying the Lipinski rule-of-five and the absorption, distribution, metabolism, excretion and overall ADME criteria. **d,** Numbers of major CYP enzymes predicted to be inhibited and Lipinski violations per molecule. The analysis included 8,700 molecular records representing 8,653 unique SMILES across 97 common benchmark targets. Classification endpoints used a probability threshold of 0.5; distribution and excretion pass ranges were defined using pooled 5th–95th percentile intervals.

**Supplementary Fig. 24. ADME profiles of FLOWR-generated molecules. a,** Predicted prevalence of categorical absorption, distribution and CYP-inhibition endpoints. **b,** Distributions of aqueous solubility, Caco-2 permeability, plasma protein binding, volume of distribution, hepatocyte clearance and half-life; dashed lines indicate medians. **c,** Proportions satisfying the Lipinski rule-of-five and domain-specific and overall ADME criteria. **d,** Per-molecule distributions of predicted major-CYP inhibition and Lipinski violations. The analysis included 9,695 molecular records representing 9,674 unique SMILES across 97 common benchmark targets. Classification endpoints used a probability threshold of 0.5; distribution and excretion pass ranges were defined using pooled 5th–95th percentile intervals.

**Supplementary Fig. 25. Predicted toxicity profiles of PFM-generated molecules. a,** Prevalence of predicted liabilities across individual toxicity endpoints. **b,** Probability distributions for selected clinically relevant endpoints, including drug-induced liver injury (DILI), hERG liability, skin reaction, Ames mutagenicity, carcinogenicity and clinical toxicity. The dashed line marks the classification threshold of 0.5. **c,** Distribution of molecules according to the number of toxicity endpoints predicted positive. **d,** Toxicity burden within the general-toxicity, nuclear-receptor and stress-response classes, shown as the mean endpoint-positive rate and the proportion positive for at least one endpoint. The analysis included 16,430 molecules across 173 targets.

**Supplementary Fig. 26. Predicted toxicity profiles of MolJO-generated molecules. a,** Prevalence of predicted liabilities across individual toxicity endpoints. **b,** Probability distributions for selected clinically relevant toxicity endpoints; the dashed line indicates the 0.5 classification threshold. **c,** Distribution of the number of positive toxicity endpoints per molecule. **d,** Mean endpoint-positive rate and prevalence of at least one positive endpoint within the general-toxicity, nuclear-receptor and stress-response classes. The analysis included 17,383 molecules across 175 targets.

**Supplementary Fig. 27. Predicted toxicity profiles of DrugFlow-generated molecules. a,** Prevalence of predicted liabilities across individual toxicity endpoints. **b,** Probability distributions for selected clinically relevant endpoints, with the dashed line marking the 0.5 classification threshold. **c,** Per-molecule multi-endpoint toxicity burden. **d,** Mean endpoint-positive rate and prevalence of at least one positive prediction within three mechanistic toxicity classes. The analysis included 17,601 molecules across 163 targets.

**Supplementary Fig. 28. Predicted toxicity profiles of FlowMol-generated molecules. a,** Prevalence of predicted liabilities across individual toxicity endpoints. **b,** Probability distributions for selected clinically relevant endpoints; the dashed line denotes the 0.5 classification threshold. **c,** Distribution of molecules by the number of positive toxicity endpoints. **d,** Toxicity burden within the general-toxicity, nuclear-receptor and stress-response classes. The analysis included 17,498 molecules across 175 targets.

**Supplementary Fig. 29. Predicted toxicity profiles of Pocket2Mol-generated molecules. a,** Prevalence of predicted liabilities across individual toxicity endpoints. **b,** Probability distributions for selected clinically relevant endpoints, with the dashed line indicating the 0.5 classification threshold. **c,** Distribution of the number of positive toxicity endpoints per molecule. **d,** Mean endpoint-positive rate and prevalence of at least one positive endpoint within the general-toxicity, nuclear-receptor and stress-response classes. The analysis included 18,682 unique molecules.

**Supplementary Fig. 30. Predicted toxicity profiles of ShEPhERD-generated molecules. a,** Prevalence of predicted liabilities across individual toxicity endpoints. **b,** Probability distributions for selected clinically relevant endpoints; the dashed line marks the 0.5 classification threshold. **c,** Per-molecule multi-endpoint toxicity burden. **d,** Toxicity burden within the general-toxicity, nuclear-receptor and stress-response classes. The analysis included 16,307 molecules across 165 targets.

**Supplementary Fig. 31. Predicted toxicity profiles of REINVENT4-generated molecules. a,** Prevalence of predicted liabilities across individual toxicity endpoints. **b,** Probability distributions for selected clinically relevant endpoints, with the dashed line marking the 0.5 classification threshold. **c,** Distribution of molecules according to the number of positive toxicity endpoints. **d,** Mean endpoint-positive rate and prevalence of at least one positive endpoint within three mechanistic toxicity classes. The analysis included 11,137 molecules across 114 targets.

**Supplementary Fig. 32. Predicted toxicity profiles of ReaSyn-generated molecules. a,** Prevalence of predicted liabilities across individual toxicity endpoints. **b,** Probability distributions for selected clinically relevant endpoints; the dashed line denotes the 0.5 classification threshold. **c,** Per-molecule multi-endpoint toxicity burden. **d,** Toxicity burden within the general-toxicity, nuclear-receptor and stress-response classes. The analysis included 12,094 molecules across 151 targets.

**Supplementary Fig. 33. Predicted toxicity profiles of SynFormer-generated molecules. a,** Prevalence of predicted liabilities across individual toxicity endpoints. **b,** Probability distributions for selected clinically relevant endpoints, with the dashed line indicating the 0.5 classification threshold. **c,** Distribution of the number of positive toxicity endpoints per molecule. **d,** Mean endpoint-positive rate and prevalence of at least one positive endpoint within three mechanistic toxicity classes. The analysis included 14,603 molecules across 164 targets.

**Supplementary Fig. 34. Predicted toxicity profiles of PrexSyn-generated molecules. a,** Prevalence of predicted liabilities across individual toxicity endpoints. **b,** Probability distributions for selected clinically relevant endpoints; the dashed line marks the 0.5 classification threshold. **c,** Per-molecule multi-endpoint toxicity burden. **d,** Toxicity burden within the general-toxicity, nuclear-receptor and stress-response classes. The analysis included 13,804 molecules across 163 targets.

**Supplementary Fig. 35. Predicted toxicity profiles of TargetDiff-generated molecules. a,** Prevalence of predicted liabilities across individual toxicity endpoints. **b,** Probability distributions for selected clinically relevant endpoints, including skin reaction, Ames mutagenicity, drug-induced liver injury (DILI), hERG liability, clinical toxicity and carcinogenicity. The dashed line indicates the classification threshold of 0.5. **c,** Distribution of molecules according to the number of toxicity endpoints predicted positive. **d,** Toxicity burden within the general-toxicity, nuclear-receptor and stress-response classes, shown as the mean endpoint-positive rate and the proportion of molecules positive for at least one endpoint. The analysis included 14,779 molecules across 174 targets.

**Supplementary Fig. 36. Predicted toxicity profiles of FLOWR-generated molecules. a,** Prevalence of predicted liabilities across individual toxicity endpoints. **b,** Probability distributions for selected clinically relevant endpoints, including DILI, hERG liability, skin reaction, Ames mutagenicity, carcinogenicity and clinical toxicity. The dashed line indicates the classification threshold of 0.5. **c,** Distribution of molecules according to the number of toxicity endpoints predicted positive. **d,** Mean endpoint-positive rate and prevalence of at least one positive endpoint within the general-toxicity, nuclear-receptor and stress-response classes. The analysis included 16,386 molecules across 164 targets.

**Supplementary Fig. 37. State-aware functional classification pipeline.** Overview of the state-aware functional classification (SAFC) framework used to prioritize generated molecules. Representative active and inactive ZAK conformations, exemplified by PDB structures 6JUT and 5HES, were subjected to explicit-solvent molecular dynamics, and representative receptor snapshots were sampled from the resulting trajectories. Experimentally characterized ligands from ChEMBL were assigned functional or non-functional labels using available IC₅₀, Kd and Ki measurements, whereas compounds with uncertain annotations were excluded. Ligands were docked against representative receptor states, and the resulting complexes were encoded as heterogeneous graphs comprising protein, ligand and virtual aromatic nodes connected by interaction-fingerprint, covalent and distance edges. Node and edge embeddings were processed through an EquiScore-based transfer-learning architecture with iterative edge-aware attention, followed by a deep neural network that classified compounds as functional or non-functional.

**Supplementary Table 1.** Sampling configuration, benchmark composition, and computational resources used for molecular generation across 176 protein-ligand systems.

**Supplementary Table 2.** Target completion, runtime, and observed peak GPU memory across molecular generation workflows. All workflows were configured to generate 100 molecules per target. Target completion was determined from the corresponding workflow execution records and is reported over the full 176-target benchmark; non-completed targets remained in the denominator. PDB 5OHE was retained in the benchmark; for affected workflows, its Fe–N dative coordination bond caused an RDKit ligand-preprocessing failure. Runtime summarizes the available per-target timing records and was analyzed independently of target completion. Observed peak VRAM denotes the maximum recorded device-level GPU-memory use under the evaluated execution setting. †FlowMol is an unconditional, target-agnostic baseline. ‡PrexSyn and SynFormer VRAM values were obtained from run-wide rather than target-specific monitoring. NR, not recorded.

