## Supplementary Files for "Systematic Benchmarking of AI-Based Molecular Generation Models for Structure-Based Drug Design"

#### Supplementary Note S1. Dataset curation for benchmarking

The benchmarking dataset was organized into complementary structural-generation and reference-guided optimization cohorts to enable both standardized model comparison and external generalization assessment. The primary structure-based benchmark comprised 61 protein–ligand systems adopted from the previously published AlignDockBench[1] resource, spanning diverse therapeutic target classes, including kinases, proteases, nuclear receptors and G-protein–coupled receptors; the associated crystallographic ligands and binding-site definitions were retained as target-specific references, and all source PDB identifiers and original dataset provenance were recorded. To reduce dependence on a single previously established benchmark, we additionally assembled two non-overlapping external evaluation cohorts. First, 40 systems were selected from the official 308-complex PoseBusters[2] subset after excluding PDB entries present in AlignDockBench; diversity was enforced using receptor-sequence clustering with MMseqs2, Bemis–Murcko scaffold uniqueness, ligand molecular-weight strata and binding-pocket size, thereby enriching for recently deposited, chemically diverse protein–ligand complexes suitable for evaluating geometric plausibility and pose quality. Second, 75 systems were selected from the official PLINDER[3] test split using predefined quality criteria, including successful structural validation, a single proper ligand chain, ligand molecular weight of 200–800 Da, pocket size of 5–100 residues and crystallographic resolution ≤3.0 Å when available; no more than one system was retained per PLINDER interaction cluster, and PDB entries already represented in AlignDockBench or PoseBusters were excluded. The PLINDER subset was further stratified by ligand size, pocket size, uniqueness category and availability of linked apo or predicted structures to support leakage-aware assessment across heterogeneous receptor states. Pocket-conditioned de novo generation models were evaluated using the same receptor structures and pocket definitions within each cohort, whereas synthesis- and ligand-guided optimization models, including ReaSyn, PrexSyn and SynFormer, were evaluated using target-matched, co-crystallized ligands and experimentally validated compounds as starting molecules. Canonical SMILES, InChIKeys, target assignments and bioactivity annotations were cross-referenced against regulatory and curated chemical databases, and compounds without a documented target relationship were excluded from target-specific analyses. Across all cohorts, receptor and ligand inputs were processed using a consistent preparation workflow, and complete provenance was retained, including source database, dataset release, retrieval date, PDB and ligand identifiers, model-specific input files, random seeds and exclusion criteria. The three structural cohorts were analyzed separately rather than pooled, allowing AlignDockBench to serve as the primary comparison set, PoseBusters as a recent structural-generalization set and PLINDER as a similarity-aware external test set.

**Supplementary Note S2. Sampling configuration, benchmark composition and computational resources.** Molecular generation was evaluated across 176 protein–ligand systems comprising 61 AlignDockBench, 40 PoseBusters and 75 PLINDER targets. Each model was requested to generate up to 100 molecules per target using a fixed random seed of 2026. Generation was performed within a 23-Å sampling box using a beam size of 300 and a maximum of 50 generation steps. The focal-site, position, element, atom-existence and bond thresholds were set to 0.50, 0.25, 0.30, 0.60 and 0.40, respectively. Each generation process was allocated one NVIDIA A100 GPU and one CPU core, and targets were processed sequentially to maintain consistent computational conditions. Molecular structures in SDF and SMILES formats, execution logs and configuration files were retained for downstream analysis, whereas intermediate .pt files were disabled or removed to reduce storage requirements. All 12 workflows were evaluated across the same 176-target benchmark. Pocket2Mol, FlowMol, and MolJO showed the highest target completion, each completing 175 targets (99.4%). TargetDiff completed 174 targets (98.9%), followed by PFM with 173 (98.3%). ShEPhERD completed 165 targets (93.8%), while FLOWR, PrexSyn, and SynFormer each completed 164 (93.2%) and DrugFlow completed 163 (92.6%). ReaSyn completed 153 targets (86.9%), whereas REINVENT4 Mol2Mol showed the lowest target completion at 114 of 176 targets (64.8%). A notable exception not attributable to the generative methods themselves was target PDB 5OHE, where an Fe–N dative coordination bond triggered an RDKit ligand-preprocessing failure that prevented eight workflows from being run. Pocket2Mol, ShEPhERD, FLOWR, and REINVENT4 Mol2Mol were unaffected by this preprocessing issue and were evaluated on this target. Among the available timing records, the flow-based workflows were consistently faster than the evaluated diffusion-based workflows. FLOWR had a median runtime of 1.0 min per target, followed by FlowMol at 1.3 min, DrugFlow at 5.2 min, and PFM at 7.1 min. In comparison, TargetDiff and ShEPhERD required median runtimes of 27.2 and 36.8 min, respectively. Pocket2Mol occupied an intermediate position at 12.5 min, whereas MolJO had the longest median runtime at 43.4 min. The synthesis-aware workflows were also computationally rapid. SynFormer, PrexSyn, and ReaSyn had median runtimes of 0.7, 0.9, and 1.9 min per target, respectively. REINVENT4 Mol2Mol required a median of 2.4 min among its recorded successful runs, although its substantially lower target completion distinguished it from the other fast workflows. Observed peak GPU-memory use varied markedly both between and within architecture families. Among the flow-based workflows, FlowMol, DrugFlow, and PFM reached observed peaks of 7.5, 12.4, and 17.9 GiB, respectively, whereas FLOWR reached 78.6 GiB. TargetDiff reached 21.7 GiB, while ShEPhERD reached 65.4 GiB. Pocket2Mol required 18.4 GiB, and MolJO reached 39.1 GiB. Run-wide monitoring recorded 7.8 GiB for both PrexSyn and SynFormer, while GPU-memory measurements were unavailable for ReaSyn and REINVENT4 Mol2Mol. Taken together, PFM and Pocket2Mol showed favorable balances of completion, runtime, and observed memory use among the target-conditioned 3D workflows. TargetDiff and MolJO maintained similarly high target completion but required substantially longer runtimes. FLOWR provided the shortest runtime among the target-conditioned flow methods but also showed the highest observed peak VRAM. PrexSyn and SynFormer supported rapid execution with completion rates above 93%, whereas REINVENT4 Mol2Mol combined short recorded runtimes with substantially lower benchmark-wide completion.

**Supplementary Note S3. Explanation about excluding AI models in this study**

Although multiple generative AI-based small molecule generation tools have been proposed in recent years, not all methods are directly comparable due to substantial differences in their design objectives, input requirement, training datasets, and evaluation metrices protocols. Models such as EDM[4], GeoLDM[5], MolDiff[6], MiDiTo[7] are primarily designed to learn general molecular distributions and generate chemically valid molecules without explicit target-specific conditioning; although their outputs can be screened post hoc, their native objective is not target-guided molecular design. Text-to-molecule models, such as MolT5[8], BioT5[9], and MolReGPT[10], generate molecules from natural-language descriptions, making performance highly dependent on prompt formulation and textual conditioning rather than standardized molecular, structural, or target-specific inputs. Similarly, general property-optimization or classical lead-optimization models, including JT-VAE[11], MIMOSA[12], MARS[13], and MolGen[14] are typically evaluated through model-specific seed molecules, benchmark objectives, which makes their performance strongly dependent on input-seed selection and optimization settings. Therefore, these methods were excluded because their native task settings and input protocols are not sufficiently aligned with the target-specific comparison framework used in this study.

**Supplementary Note S4. Pocket2Mol implementation and benchmarking protocol**

Pocket2Mol was included as an autoregressive, pocket-conditioned three-dimensional molecular generation model. The complete benchmark comprised 176 protein–ligand systems from AlignDockBench, PoseBusters and PLINDER. For each system, the receptor structure was supplied to the model, whereas the crystallographic ligand was used only to locate the pocket. The pocket center was defined from the ligand heavy-atom bounding box, and a fixed 23 Å cubic sampling region was applied. Generation used the official pretrained checkpoint with a random seed of 2026, beam size of 300, maximum of 50 autoregressive steps and the default sampling thresholds. One hundred molecules were requested per target using one NVIDIA A100 GPU. A low-storage copy of the sampling script was used in which only intermediate PyTorch checkpoint writing was disabled; model weights and sampling decisions were unchanged. Final SDF structures, canonical SMILES, configurations and logs were retained. A target was classified as complete when at least 100 structures were produced. Molecules were standardized using RDKit, invalid structures and duplicate canonical SMILES were removed, and up to 100 valid unique molecules per target were retained. Runtime, memory, GPU utilization, power consumption, throughput and output-storage requirements were recorded using the common scalability workflow.

Pocket2Mol produced a favorable balance of chemical properties and docking performance across 17,296 molecules. Median QED, normalized SA and Vina score were 0.65, 0.78 and −7.50 kcal $mol^{-1}$, respectively; 25.8% of molecules passed all three criteria, the second-highest joint success rate among the models. Pocket2Mol ADME analysis achieved complete median target-level Lipinski and absorption pass rates and an overall ADME soft-pass rate of 92.9%. Predicted intestinal absorption and bioavailability were high, whereas P-gp inhibition was uncommon (3.9%). Its comparatively lower metabolism pass rate of 70.9% was primarily associated with predicted CYP1A2 and CYP2C19 inhibition, indicating that metabolic interaction liabilities remain relevant despite otherwise favorable absorption and physicochemical profiles. Pocket2Mol toxicity profiling included 18,682 molecules; Approximately 20.9% of molecules had no positive endpoint, whereas 7.8% were positive for at least six endpoints. General toxicity represented the principal liability class, with a mean endpoint-positive rate of 23.8% and 78.5% of molecules positive for at least one endpoint. Nuclear-receptor and stress-response liabilities were less prevalent, affecting at least one endpoint in 18.6% and 17.9% of molecules, respectively. Pocket2Mol therefore showed a predominantly general-toxicity burden with relatively limited mechanistic stress-response activity.

**Supplementary Note S5. ShEPhERD implementation and benchmarking protocol**

ShEPhERD was included as a reference-ligand-conditioned three-dimensional generative model. It does not directly use receptor coordinates during generation but conditions sampling on the shape, electrostatic surface and pharmacophore features of a reference ligand. For each benchmark system, the crystallographic ligand was sanitized, protonated and retained in its bound conformation. xTB single-point calculations in implicit water were used to assign partial charges without geometry optimization. The centered ligand coordinates were then used to extract surface, electrostatic and pharmacophore conditioning features. Generation employed the pretrained mosesaq checkpoint, which jointly models these three representations. One hundred valid unique molecules were requested for each of the 176 systems using 400 diffusion steps, a random seed of 2026, a batch size of 5 and one NVIDIA A100 GPU. Invalid structures and duplicate canonical SMILES were discarded during sampling.

Generated molecules were retained as a multi-record SDF file together with canonical SMILES, the prepared reference ligand, extracted conditioning profile, metadata and logs. Targets were classified as complete, partial or failed according to the number of valid unique molecules recovered. Because ShEPhERD is ligand-conditioned, its outputs were treated as ligand-guided bioisosteric proposals and evaluated separately through the common docking, pose, molecular-property, ADME and scalability workflows.

ShEPhERD generated 16,107 molecules with median QED, normalized SA and Vina values of 0.57, 0.62 and −8.47 kcal $mol^{-1}$. While 73.6% met the docking criterion, only 24.7% and 37.7% passed the QED and SA thresholds, respectively, producing a joint-pass rate of 10.2%. ShEPhERD produced a balanced predicted ADME profile, with complete Lipinski compliance and an overall soft-pass rate of 94.0%. Absorption, distribution and excretion pass rates were 98.0%, 88.0% and 98.0%, respectively. The metabolism pass rate was lower at 71.0%, driven mainly by predicted CYP3A4 inhibition, whereas predicted P-gp inhibition remained moderate at 25.0%. ShEPhERD toxicity profiling included 16,307 molecules across 165 targets. Only 6.8% of molecules were negative across all toxicity endpoints, whereas 6.2% were positive for at least six endpoints. The majority carried between one and three positive liabilities, with the two-endpoint category being the largest at 27.4%. General-toxicity predictions were widespread: the mean endpoint-positive rate was 33.5%, and 92.2% of molecules were positive for at least one general-toxicity endpoint. Nuclear-receptor liabilities were uncommon, affecting at least one endpoint in 10.7% of molecules, whereas stress-response liabilities affected 30.3%. ShEPhERD therefore produced widespread general-toxicity predictions but relatively limited nuclear-receptor activity.

**Supplementary Note S6. FLOWR implementation and benchmarking protocol**

FLOWR was included as a pocket-conditioned three-dimensional ligand generation model based on continuous and categorical flow matching. For each of the 176 benchmark systems, the receptor structure was supplied as the protein input, and the crystallographic ligand was used to define the local binding pocket. Generation used the official pretrained flowr_noHs.ckpt checkpoint in rigid pocket mode with arch=pocket, pocket-type=holo, a 6 Å pocket cutoff, sampled molecular sizes, categorical uniform sampling, 100 integration steps and a random seed of 2026. One hundred molecules were requested per target using one NVIDIA A100 GPU. To reduce storage consumption, a separate copy of the inference script was used in which trajectory saving was disabled; model weights, conditioning inputs and molecular sampling were unchanged. The complete pre-diversity output was exported as samples_protein_all.sdf and used for primary benchmarking, while FLOWR's diversity-filtered output was retained separately. Intermediate trajectory and PyTorch files were removed after successful SDF conversion. Targets producing 100 structures were classified as complete, those producing fewer as partial and those lacking usable structures as failed. Generated molecules were standardized using RDKit and evaluated using the common validity, uniqueness, docking, pose-quality, physicochemical, ADME and scalability pipelines.

FLOWR generated 16,047 molecules with median QED and normalized SA values of 0.51 and 0.70 and a median Vina score of −7.57 kcal $mol^{-1}$. The QED, SA and docking criteria were met by 22.2%, 58.9% and 56.1% of molecules, respectively, with 11.2% satisfying all three requirements. FLOWR ADME profiling achieved complete median Lipinski compliance and strong absorption and excretion performance, but its overall ADME soft-pass rate was more moderate at 89.0%. The metabolism pass rate was 54.0%, reflecting appreciable predicted inhibition of CYP2C9, CYP2C19 and CYP3A4. P-gp inhibition was also predicted for 40.0% of molecules per target, suggesting that transporter and metabolic liabilities constitute the principal limitations of the FLOWR-generated set. FLOWR showed a markedly different toxicity profile from the other methods. The analysis comprised 16,386 molecules across 164 targets, of which 89.9% were negative across all 18 endpoints. Only 1.2% of molecules were positive for at least six endpoints, and 3.0% were positive for four or more. At least one general-toxicity liability was predicted for 9.2% of molecules, with a mean endpoint-positive rate of 3.1%. Nuclear-receptor and stress-response liabilities were similarly uncommon, affecting 1.4% and 4.0% of molecules, respectively. These predictions identify FLOWR as producing the lowest computational toxicity burden in this analysis, although the unusually favorable profile should be interpreted alongside its distinct generated chemical-property distribution and applicability domain.

#### Supplementary Note S7. TargetDiff implementation and benchmarking protocol

TargetDiff was included as a pocket-conditioned three-dimensional molecular generation model. It uses an SE(3)-equivariant diffusion process to jointly generate ligand atom types and coordinates within a protein-binding pocket. For each of the 176 benchmark systems, the standardized receptor structure was supplied as the protein context, and the crystallographic ligand was used only to identify the binding pocket. Generation was performed using the official pretrained model and inference workflow, with 100 molecules requested per target and a fixed random seed. Generated structures were exported in SDF format and standardized using RDKit. Invalid structures and duplicate canonical SMILES were removed, and up to 100 valid unique molecules per target were retained. Target completion, runtime, memory consumption, GPU utilization, output size and throughput were recorded using the common scalability workflow. TargetDiff was evaluated as a receptor-pocket-conditioned de novo generator. TargetDiff paper and implementation.

Among 14,872 molecules, TargetDiff showed median QED and normalized SA values of 0.46 and 0.57, with 15.8% and 23.2% meeting the respective thresholds. Although 72.6% achieved Vina scores ≤ −7 kcal $mol^{-1}$ (median, −8.49), only 4.1% satisfied all three criteria, indicating strong predicted docking but limited joint drug-likeness and synthesizability. TargetDiff ADME profiling showed a broadly favorable predicted ADME profile, with a median target-level soft-pass rate of 94.3%. Absorption, distribution and excretion pass rates were 96.7%, 88.2% and 95.9%, respectively, while the metabolism pass rate reached 82.1%. The model also showed high Lipinski compliance (97.8%) and comparatively low predicted inhibition of individual CYP isoforms, although P-gp inhibition was predicted for 23.7% of molecules. The TargetDiff toxicity analysis included 14,779 molecules from 174 targets. Of these, 28.9% had no predicted toxicity liability, whereas 7.7% were positive for at least six endpoints. At the mechanistic-class level, 24.5% of general-toxicity endpoints were positive on average, and 69.6% of molecules were positive for at least one general-toxicity endpoint. Nuclear-receptor liabilities were comparatively limited, with a mean endpoint-positive rate of 1.8% and 10.1% of molecules positive for at least one nuclear-receptor endpoint. Stress-response liabilities were more frequent, with corresponding rates of 8.5% and 25.7%. Thus, TargetDiff produced a comparatively large liability-free fraction, although a smaller subgroup exhibited substantial multi-endpoint burden.

**Supplementary Note S8. DrugFlow implementation and benchmarking protocol**

DrugFlow was included as a pocket-conditioned three-dimensional molecular generation model that combines continuous flow matching for atomic coordinates with discrete probabilistic transitions for molecular composition. The model directly incorporates the protein-binding environment and was therefore evaluated as a structure-based de novo generation method. For each benchmark target, the receptor structure and ligand-defined binding pocket were prepared using the standardized input pipeline, and 100 candidate molecules were requested using the same checkpoint and generation settings. Molecular structures were exported in SDF format and processed using the common RDKit validity, canonicalization and deduplication workflow. Up to 100 valid unique molecules were retained per target. DrugFlow outputs were subjected to the same docking, physicochemical, diversity, ADME, toxicity and scalability analyses applied to the other models. Because DrugFlow also provides uncertainty-related outputs, these were retained when available but were not treated as experimentally calibrated confidence estimates. DrugFlow study.

DrugFlow generated 12,018 molecules with median QED, normalized SA and Vina values of 0.53, 0.70 and −8.67 kcal $mol^{-1}$, respectively. Although only 24.1% passed the QED threshold, 60.1% passed the SA threshold and 75.4% met the docking criterion, resulting in a joint-pass rate of 13.4%. DrugFlow ADME profiling showed high absorption (99.1%) and excretion (97.9%) pass rates, resulting in an overall median ADME soft-pass rate of 92.0%. Lipinski compliance was also high at 97.4%, although P-gp inhibition occurred in a median of 31.5% of molecules per target. The metabolism pass rate was more moderate at 63.1%, with CYP3A4 representing the most prominent predicted CYP liability. DrugFlow toxicity analysis included 17,601 molecules from 163 targets. The positive-endpoint counts were broadly distributed: 18.1% of molecules had no predicted liability, while 20.6%, 19.1% and 14.8% carried one, two and three positive endpoints, respectively. A further 17.3% were positive for four or five endpoints and 10.1% for at least six. General-toxicity endpoints had a mean positive rate of 29.4%, with 80.6% of molecules positive for at least one such endpoint. Nuclear-receptor activity remained comparatively limited, whereas 28.5% of molecules were positive for at least one stress-response endpoint. These results indicate heterogeneous DrugFlow output, containing both liability-free molecules and a substantial high-burden subset.

**Supplementary Note S9. FlowMol implementation and benchmarking protocol**

FlowMol was included as an unconditional three-dimensional molecular generation baseline. The method uses multimodal flow matching to jointly generate atomic coordinates, atom identities, charges and bond information, but it does not directly condition generation on a protein receptor or binding pocket. Consequently, neither the receptor coordinates nor the reference ligand was supplied during molecular sampling. Independent sampling jobs were used to generate the requested molecular sets, after which the resulting compounds were evaluated against the benchmark targets using the common docking and property-analysis pipeline. One hundred molecules were requested for each target-indexed run, and invalid or duplicate structures were removed using RDKit before retaining up to 100 valid unique molecules. FlowMol therefore served as a target-agnostic control for determining the added value of explicit pocket conditioning. The exact FlowMol release should be reported because its successive versions use different treatments of continuous and categorical variables. Official FlowMol implementation.

FlowMol showed the strongest overall chemical-property profile, with median QED and normalized SA values of 0.75 and 0.86. Of 17,183 molecules, 62.9% passed the QED criterion, 97.8% passed the SA criterion and 57.1% achieved Vina scores ≤ −7 kcal $mol^{-1}$, yielding the highest joint-pass rate of 34.7%. FlowMol ADME profiling generated molecules with complete median Lipinski compliance, high intestinal absorption and bioavailability, and comparatively favorable Caco-2 permeability. However, its overall ADME soft-pass rate was lower at 79.0%, largely because only 40.0% of molecules per target passed the metabolism criterion. Predicted inhibition was particularly prevalent for CYP1A2 and CYP2C19, and the molecules also showed relatively high plasma protein binding and hepatocyte clearance. FlowMol toxicity profiling comprised 17,498 molecules across 175 targets. Only 4.4% of molecules were negative across all 18 endpoints, indicating that 95.6% carried at least one predicted liability; however, only 6.0% were positive for at least six endpoints. Most molecules were concentrated in the one- and two-positive-endpoint categories, representing 29.7% and 29.5% of the collection, respectively. At least one general-toxicity liability was predicted for 95.3% of molecules, although the mean positive rate across individual general-toxicity endpoints was 29.9%. Nuclear-receptor and stress-response liabilities affected 20.4% and 25.1% of molecules, respectively. FlowMol therefore produced broadly distributed but generally low-to-moderate per-molecule toxicity burdens.

**Supplementary Note S10. MolJO implementation and benchmarking protocol**

MolJO was included as a structure-based molecular optimization model rather than an unconstrained de novo generator. It operates in a continuous and differentiable space derived from Bayesian flow networks and applies gradient-based guidance jointly to atomic coordinates and discrete atom identities. For every benchmark system, the standardized receptor pocket and corresponding reference ligand were used to initialize target-specific molecular optimization. The official pretrained checkpoint and common optimization settings were applied across targets, with up to 100 optimized candidates requested per system. Generated structures were standardized and deduplicated using canonical isomeric SMILES, and a maximum of 100 RDKit-valid unique molecules was retained. MolJO was evaluated for scaffold modification, molecular-property improvement and pocket compatibility, but its results were interpreted separately from fully de novo methods because generation begins from an existing molecular structure. Docking, QED, normalized synthetic accessibility, diversity, ADME, toxicity and computational efficiency were evaluated using the unified downstream pipeline. Official MolJO implementation.

MolJO exhibited strong predicted docking, with a median Vina score of −9.60 kcal $mol^{-1}$ and 78.9% of 16,706 molecules meeting the docking criterion. Median QED and normalized SA were 0.54 and 0.71, respectively, and 11.3% of molecules passed the combined QED–SA–Vina filter. MolJO ADME profiling showed high predicted absorption, bioavailability and Lipinski compliance but the lowest overall metabolic performance. Only 21.0% of molecules per target passed the metabolism criterion, and predicted inhibition was frequent for CYP1A2, CYP2C9 and particularly CYP2C19. MolJO also showed the lowest predicted aqueous solubility, the highest plasma protein binding and frequent P-gp inhibition, resulting in an overall ADME soft-pass rate of 86.0% despite favorable absorption and excretion scores. MolJO produced the highest multi-endpoint toxicity burden among the individual model outputs. The analysis included 17,383 molecules across 175 targets. Only 3.3% of molecules were negative across all endpoints, whereas 44.6% were positive for at least six endpoints and an additional 21.9% were positive for four or five. General-toxicity endpoints had a mean positive rate of 44.4%, and 96.6% of molecules were positive for at least one endpoint in this class. Nuclear-receptor liabilities were also more prevalent than for the other models, with a mean endpoint-positive rate of 9.9% and 47.3% of molecules positive for at least one endpoint. Stress-response liabilities were particularly pronounced, with corresponding values of 32.0% and 70.7%. MolJO therefore generated the most concentrated high-liability profile, emphasizing the need for stringent toxicity filtering during subsequent candidate prioritization.

**Supplementary Note S11. PFM implementation and benchmarking protocol**

Perturbed Flow Matching (PFM) was included as a target-conditioned structure-based molecular generation model. PFM modifies conventional flow matching by introducing a perturbed conditional probability path intended to reduce the number of sampling steps required for generating pocket-compatible three-dimensional molecules. For each benchmark target, the receptor structure was provided as the conditioning input, whereas the bound ligand was used to define the binding-pocket region. The same pretrained checkpoint, integration settings, molecular-count target and random-seed policy were applied across all 176 systems. One hundred molecules were requested per target, and outputs generated were converted to standardized SDF and canonical-SMILES representations. Invalid structures and duplicate molecules were removed, and up to 100 valid unique candidates were retained. PFM was assessed using the same validity, docking, physicochemical, ADME, toxicity and scalability endpoints, allowing its sampling efficiency to be compared with diffusion- and flow-based pocket-conditioned generators. PFM paper.

PFM produced the most favorable median docking score (−10.17 kcal $mol^{-1}$), with 80.7% of its 16,235 molecules meeting the Vina threshold. However, median QED and normalized SA were comparatively low at 0.49 and 0.53, and only 2.4% passed all three criteria, demonstrating a pronounced trade-off between predicted binding and chemical developability. PFM ADME profiling achieved the highest median overall ADME soft-pass rate among the evaluated models (97.0%), supported by high absorption (98.0%), metabolism (88.5%) and complete excretion pass rates. Individual CYP inhibition rates were generally low, and the median CYP burden was zero. Nevertheless, PFM molecules showed relatively poor predicted aqueous solubility, high plasma protein binding and frequent P-gp inhibition (55.7%), identifying potential exposure-related liabilities that are not fully captured by the composite soft-pass score. PFM profiling comprised 16,430 molecules across 173 targets. Approximately 26.6% of molecules had no predicted positive endpoint, whereas 6.7% were positive for at least six endpoints. General-toxicity liabilities were less prevalent than for several other models, with a mean endpoint-positive rate of 17.4% and 62.5% of molecules carrying at least one general-toxicity liability. Nuclear-receptor activity was predicted for 24.3% of molecules. Stress-response liabilities were comparatively prominent, with a mean endpoint-positive rate of 14.7% and 45.7% of molecules positive for at least one stress-response endpoint. PFM therefore showed a relatively large liability-free fraction but a distinct enrichment for cellular stress-response predictions.

**Supplementary Note S12. SynFormer implementation and benchmarking protocol**

SynFormer was included as a synthesis-aware molecular generation model. Rather than directly generating unconstrained molecular graphs, SynFormer constructs molecules through reaction pathways composed of predefined reaction templates and purchasable building blocks, thereby restricting its outputs to the model-defined synthesizable chemical space. The receptor structure was not directly supplied during generation. For each benchmark system, the corresponding reference ligand was used as the molecular query or optimization seed, and candidate synthetic pathways were sampled using the official pretrained model and search workflow. Up to 100 product molecules were requested per target. Final products were extracted from successful pathways, standardized using RDKit and deduplicated by canonical isomeric SMILES. Candidates lacking a complete molecular product or valid reaction pathway were classified as unsuccessful outputs. Generated molecules were subsequently evaluated through the same docking, QED, normalized SA, diversity, ADME and toxicity pipeline. SynFormer was interpreted as a synthesis-constrained ligand-optimization baseline rather than a receptor-pocket-conditioned generator. Official SynFormer implementation.

SynFormer generated 14,661 molecules with a low median QED of 0.33 but a more favorable median normalized SA of 0.70. Although 59.1% passed the SA criterion and 64.7% achieved Vina scores ≤ −7 kcal $mol^{-1}$, only 13.0% met the QED threshold and 8.8% passed the combined filter. SynFormer ADME profiling showed complete median absorption and near-complete excretion pass rates, producing an overall ADME soft-pass rate of 92.0%. However, Lipinski compliance was lower and more target dependent (median, 81.0%), and only 45.5% of molecules passed the metabolism criterion. CYP3A4 was the dominant predicted metabolic liability, while comparatively low aqueous solubility and high plasma protein binding were also observed. SynFormer profiling included 14,603 molecules across 164 targets. Approximately 12.4% of molecules had no positive endpoint, whereas 7.8% were positive for at least six endpoints. The 4–5 endpoint category was the largest, accounting for 22.9% of molecules, and 30.7% were positive for at least four endpoints. General-toxicity liabilities occurred in 86.6% of molecules, with a mean endpoint-positive rate of 29.7%. At least one nuclear-receptor and stress-response liability was predicted for 16.8% and 39.1% of molecules, respectively. SynFormer therefore showed a broad general-toxicity burden accompanied by appreciable enrichment for multi-endpoint and stress-response predictions.

**Supplementary Note S13. PrexSyn implementation and benchmarking protocol**

PrexSyn was included as a programmable synthesis-aware molecular generation model. It uses a decoder-only transformer to autoregressively generate postfix representations of synthetic pathways constructed from reaction rules and purchasable building blocks. Generation can be conditioned on molecular descriptors, allowing exploration of synthesizable chemical space while directing outputs toward predefined property ranges. In this benchmark, the standardized reference ligand and its calculated molecular descriptors were used to establish target-specific generation or reconstruction conditions; receptor coordinates were not supplied directly to the model. Up to 100 final product molecules were requested for each system. Products from completed pathways were extracted, canonicalized and filtered using RDKit, and duplicate structures were removed before retaining a maximum of 100 valid unique molecules. PrexSyn outputs were docked into the corresponding protein pockets and analyzed using the common molecular-property, diversity, ADME, toxicity and scalability workflow. Its performance was interpreted as synthesis- and property-guided ligand generation rather than direct structure-based pocket generation. PrexSyn paper and documentation.

PrexSyn generated 13,910 molecules with a median QED of 0.36, normalized SA of 0.72 and Vina score of −8.09 kcal $mol^{-1}$. Although 63.5% passed the SA criterion and 70.0% met the docking threshold, only 11.7% achieved QED ≥ 0.70, limiting the joint-pass rate to 7.2%. PrexSyn ADME profiling showed complete median absorption and excretion pass rates and an overall ADME soft-pass rate of 91.0%. Lipinski compliance was moderately reduced to 88.0%, while the metabolism pass rate was 56.0%. Frequent predicted P-gp inhibition (53.0%) and CYP3A4 inhibition, together with relatively low aqueous solubility, suggest that transporter and metabolic optimization would be required for many PrexSyn-generated candidates. PrexSyn toxicity profiling included 13,804 molecules across 163 targets. Approximately 12.2% of molecules were negative across all endpoints, whereas 9.4% were positive for at least six endpoints. The largest burden category was four or five positive endpoints, representing 21.4% of molecules; collectively, 30.8% were positive for at least four endpoints. General-toxicity liabilities occurred in 86.9% of molecules, with a mean endpoint-positive rate of 30.0%. At least one nuclear-receptor liability was predicted for 18.8% of molecules, whereas 39.5% carried at least one stress-response liability. PrexSyn therefore showed a broad general-toxicity profile together with a substantial multi-endpoint and stress-response burden.

**Supplementary Note S14. ReaSyn implementation and benchmarking protocol**

ReaSyn was included as an iterative synthesis-pathway refinement model. The framework represents multistep synthesis using a chain-of-reaction notation and projects input molecules toward chemically related products that can be assembled from available building blocks and supported reaction transformations. For each benchmark system, the crystallographic or reference ligand was supplied as the starting molecular query, whereas the receptor structure was reserved for subsequent docking and evaluation. Candidate pathways were generated and iteratively refined using the official pretrained model and search procedure. Final products from valid completed pathways were extracted, standardized with RDKit and deduplicated using canonical isomeric SMILES. Up to 100 valid unique molecules were retained per target when available. ReaSyn was evaluated as a reference-ligand-conditioned synthesizable-analogue generator; accordingly, target specificity was assessed only after docking the generated products into the corresponding protein pocket. Reaction steps, building blocks and pathway identifiers were retained to support synthetic traceability. Official ReaSyn implementation.

ReaSyn showed a median QED of 0.31 and median normalized SA of 0.73 across 8,048 molecules. The SA and docking criteria were met by 70.2% and 65.5% of molecules, respectively, but the lower QED pass rate of 14.9% restricted the joint-pass fraction to 8.7%. ReaSyn ADME profiling generated molecules with high predicted absorption and excretion performance, yielding a median overall ADME soft-pass rate of 92.6%. In contrast, its Lipinski pass rate was only 67.0%, the lowest among the evaluated models, indicating broader physicochemical-rule violations. The metabolism pass rate was 55.4%, with CYP3A4 and P-gp inhibition representing the most prominent predicted liabilities; BBB penetration was also lower than for most other models. The ReaSyn toxicity analysis included 12,094 molecules across 151 targets. Only 9.2% of molecules were negative across all endpoints, whereas 7.7% were positive for at least six. A further 21.8% were positive for four or five endpoints, producing a combined high-burden fraction of 29.5%. General-toxicity liabilities were present in 90.2% of molecules, with a mean endpoint-positive rate of 30.0%. Nuclear-receptor liabilities affected 15.7% of molecules, whereas stress-response liabilities affected 39.0%. ReaSyn therefore produced toxicity profiles like those of SynFormer, characterized by widespread general-toxicity predictions and a notable cellular stress-response burden.

##### Supplementary Note S15. REINVENT4 implementation and benchmarking protocol

REINVENT4 was included as a two-dimensional, reference-ligand-conditioned molecular generation and optimization baseline. The model generates SMILES using recurrent or transformer-based architectures and supports transfer learning, reinforcement learning, scaffold hopping and Mol2Mol-style analogue generation. It does not directly use receptor coordinates unless receptor-dependent quantities are explicitly introduced as scoring components. In this benchmark, the crystallographic ligand associated with each target was used as the reference molecular seed, and candidate analogues were generated using a consistent Mol2Mol configuration and sampling policy. One hundred molecules were requested per target. Generated SMILES were parsed, standardized and canonicalized using RDKit; invalid outputs and duplicate canonical structures were removed, and up to 100 valid unique molecules were retained. Three-dimensional conformers were subsequently generated for docking and pose evaluation. REINVENT4 was therefore treated as a ligand-guided analogue-generation baseline, with target compatibility determined through the same downstream docking, physicochemical, ADME and toxicity analyses applied to all models. Official REINVENT4 implementation

REINVENT4 generated 10,866 molecules with median QED, normalized SA and Vina values of 0.55, 0.77 and −8.22 kcal $mol^{-1}$. Its high SA pass rate of 83.0% and docking pass rate of 72.6% compensated partly for a QED pass rate of 24.8%, producing a joint-pass rate of 17.5%. REINVENT4 ADME profiling achieved an overall median ADME soft-pass rate of 95.0%, together with complete Lipinski, absorption and excretion pass rates and the highest distribution pass rate (92.0%). Its metabolism pass rate was comparatively lower at 59.0%, largely reflecting predicted CYP3A4 inhibition. Thus, the model combined favorable predicted physicochemical and exposure-related characteristics with a remaining CYP-mediated interaction liability. REINVENT4 profiling comprised 11,137 molecules across 114 targets. Approximately 14.1% of molecules had no predicted liability, whereas 8.8% were positive for at least six endpoints. The remaining molecules were distributed relatively evenly across the one-, two-, three- and four-to-five-endpoint categories. General-toxicity predictions occurred in 83.9% of molecules, with a mean endpoint-positive rate of 28.6%. Nuclear-receptor activity was predicted for 26.0% of molecules, with a mean endpoint-positive rate of 5.2%. Stress-response liabilities affected 34.5% of molecules and had a mean endpoint-positive rate of 11.0%. REINVENT4 therefore displayed a heterogeneous burden with substantial general-toxicity and moderate stress-response predictions.

#### Supplementary Methods S16: ADME profiling

ADME predictions for RDKit-valid molecules were obtained using ADMET-AI and linked to the corresponding generative model and protein target through canonical SMILES. Binary endpoints, including human intestinal absorption, bioavailability, P-glycoprotein inhibition, blood–brain barrier penetration and inhibition of five major cytochrome P450 isoforms, were classified using a probability threshold of 0.5. Continuous endpoints comprised aqueous solubility, Caco-2 permeability, plasma protein binding, volume of distribution, hepatocyte and microsomal clearance, and half-life. Lipinski Rule-of-Five descriptors were calculated with RDKit, with molecules considered compliant when they violated no more than one of the molecular-weight, logP, hydrogen-bond donor or hydrogen-bond acceptor criteria. To control for unequal molecule counts, endpoint values were first summarized within each model–target combination and subsequently aggregated across targets using the median and interquartile range.

#### Supplementary Methods S17: Toxicity profiling

ADMET-AI predictions generated for each model were processed using a standardized, model-independent workflow. Prediction CSV files were identified within each model-specific directory, concatenated and annotated with their source file. Duplicate records were removed using the available combination of molecule identifier and SMILES, retaining the first occurrence. Endpoint values were converted to numeric format and missing or non-numeric predictions were retained as missing values rather than interpreted as negative predictions. The toxicity analysis included 18 ADMET-AI classification endpoints grouped into three mechanistic classes: general toxicity, comprising Ames mutagenicity, carcinogenicity, clinical toxicity, drug-induced liver injury (DILI), hERG liability and skin reaction; nuclear-receptor activity, comprising androgen receptor ligand-binding domain (AR-LBD), androgen receptor (AR), aryl hydrocarbon receptor (AhR), aromatase, estrogen receptor ligand-binding domain (ER-LBD), estrogen receptor (ER) and peroxisome proliferator-activated receptor-γ (PPARγ); and cellular stress response, comprising antioxidant-response element (ARE), ATAD5, heat-shock response element (HSE), mitochondrial membrane potential (MMP) and p53. ADME and cytochrome P450 interaction endpoints were standardized separately and were not included in the toxicity-only figures.

All classification outputs were treated as predicted probabilities, and an endpoint was classified as positive when its predicted probability was greater than or equal to 0.50. For each endpoint, the number of non-missing predictions, mean, median, standard deviation, minimum, maximum, 5th, 25th, 75th and 95th percentiles, and predicted-positive frequency were calculated. These probabilities represent computational liability predictions and were not interpreted as experimentally confirmed toxicities. A molecule-level toxicity burden was calculated as the number of the 18 endpoints classified as positive. Additional molecule-level variables included the fraction of available endpoints classified as positive, the presence of at least one positive endpoint, mean toxicity probability, maximum toxicity probability and the endpoint producing the maximum predicted probability. Positive-endpoint counts were grouped into six categories: 0, 1, 2, 3, 4–5 and ≥6 predicted liabilities. Within each mechanistic class, two complementary quantities were calculated: the mean positive rate across the constituent endpoints and the percentage of molecules positive for at least one endpoint in that class. Where target identifiers were available, the mean probability, positive-endpoint count and fraction of molecules with at least one predicted liability were also summarized at the target level. The 200 molecules with the largest positive-endpoint counts, ranked secondarily by maximum toxicity probability, were retained as a high-risk candidate table. For each model, the supplementary toxicity figure contained four panels. Panel a shows the predicted positive prevalence for all 18 endpoints. Panel b shows probability distributions for the six general-toxicity endpoints; boxes represent the interquartile range with the median indicated by the internal line, whiskers follow the default 1.5× interquartile-range definition, outliers are omitted from visualization, and the dashed line indicates the 0.50 classification threshold. Panel c reports the distribution of per-molecule positive-endpoint counts. Panel d compares the mean endpoint-positive rate with the percentage of molecules positive for at least one endpoint within each mechanistic class.

**Supplementary Note S18: State Aware Functional Classifier (SAFC)**

**Introduction.** Conventional structure-based virtual screening frequently evaluates ligand binding against a single receptor structure and uses docking score as a proxy for biological activity. However, favorable binding alone does not necessarily indicate that a compound will produce the intended functional effect. Functional activity may depend on compatibility with specific receptor states and on whether the associated protein–ligand interaction pattern remains reproducible across the conformational variability of the binding pocket. The state-aware functional classifier (SAFC) was developed to address this limitation by integrating predefined receptor functional states, molecular-dynamics-derived conformational ensembles, representative-conformation selection, ensemble docking, complex-level protein–ligand representations, and target-specific transfer learning. Rather than treating docking score as the final prediction endpoint, SAFC estimates functional activity from the combined ligand and receptor-conditioned interaction evidence observed across multiple representative receptor conformations.

This supplementary analysis describes the methodological basis of SAFC and evaluates its reliability using model-selection, cross-target discrimination, feature-ablation, threshold-sensitivity, and error analyses on ZAK and norepinephrine transporter datasets.

**Methods**

The target protein was represented by a set of predefined functional states

$$\mathcal{S} = \{s_1, s_2, \dots, s_M\},$$

with each state represented by a set of selected receptor conformations

$$\mathcal{R}_s = \{R_{s,1}, R_{s,2}, \dots, R_{s,K_s}\}.$$

For ligand $L_i$, docking against receptor conformation $R_{s,k}$ generated a protein–ligand complex

$$C_{i,s,k} = \mathcal{D}(L_i, R_{s,k}),$$

where $\mathcal{D}$ denotes the docking operation. Each docked complex was transformed into a complex-level representation

$$\mathbf{x}_{i,s,k} = \Phi(C_{i,s,k}) = \left[\mathbf{f}_i^{\text{lig}} \| \mathbf{f}_{i,s,k}^{\text{contact}} \| \mathbf{f}_{i,s,k}^{\text{geom}} \| \mathbf{f}_{i,s,k}^{\text{dock}}\right],$$

where $\Phi$ denotes the complex-representation function and $\|$ denotes vector concatenation. The representation combines intrinsic ligand information with receptor-conditioned contact, geometric, and docking-derived evidence. Consequently, chemically similar ligands may receive different predictions when they form different interaction patterns within the receptor ensemble, while the same ligand may produce distinct representations in different receptor states.

Complex representations were first aggregated within each functional state:

$$\mathbf{z}_{i,s} = \frac{1}{K_s}\sum_{k=1}^{K_s} \mathbf{x}_{i,s,k}.$$

The resulting state-specific representations were concatenated in a fixed state order:

$$\mathbf{z}_i = \mathbf{z}_{i,s_1} \| \mathbf{z}_{i,s_2} \| \cdots \| \mathbf{z}_{i,s_M}.$$

The target-specific classifier then estimated

$$\hat{p}_i = f_\theta(\mathbf{z}_i),$$

where $\hat{p}_i$ is defined as the SAFC score. The continuous score was retained for candidate ranking and threshold-dependent prioritization.

The complete SAFC workflow, including receptor-ensemble generation, activity-data curation, ensemble docking, complex representation, and functional classification, is summarized in Supplementary Figure 37.

Experimentally measured activity data were collected from target-specific ChEMBL records. The principal analyses were restricted to IC50 measurements to reduce heterogeneity associated with combining non-equivalent biochemical endpoints such as IC50, Ki, and Kd. Reported concentrations were converted to molar units and transformed as

$$\text{pActivity} = -\log_{10}(\text{Activity in mol/L}).$$

Molecular structures were standardized before analysis. Disconnected minor fragments and counterions were removed, and repeated activity measurements associated with the same standardized compound were summarized using the median:

$$\widetilde{pA}_i = \mathrm{median}\left(pA_{i,1}, pA_{i,2}, \ldots, pA_{i,n_i}\right).$$

High-confidence binary labels were assigned using two activity thresholds:

$$y_i = \begin{cases} 1, & \widetilde{pA}_i \geq T_{\mathrm{active}}, \\ 0, & \widetilde{pA}_i \leq T_{\mathrm{inactive}}. \end{cases}$$

Compounds satisfying

$$T_{\mathrm{inactive}} < \widetilde{pA}_i < T_{\mathrm{active}}$$

were excluded from supervised training and primary evaluation. This strategy reduced uncertainty associated with assay variation and prevented small numerical differences near a single activity boundary from generating opposing labels. The activity thresholds used to construct experimental labels were distinct from the probability threshold subsequently applied to model predictions.

Proteins sample multiple conformational microstates rather than remaining in a single static structure. Even within one broadly defined functional state, variations in side-chain orientation, loop position, pocket volume, solvent exposure, and local geometry may alter ligand placement and protein–ligand contact formation. Evaluating each ligand against a receptor ensemble therefore reduces dependence on an individual experimental structure and tests whether predicted interaction patterns remain reproducible across plausible pocket geometries.

Candidate receptor conformations were extracted from the equilibrated or structurally stable region of each molecular dynamics trajectory. Periodic-boundary artifacts were corrected before trajectory alignment and structural analysis, and frames were sampled at defined temporal intervals to reduce redundancy among adjacent and highly autocorrelated snapshots. Target-specific structural descriptors were used to verify that retained conformations remained consistent with their assigned functional state.

Representative conformations were selected according to structural variation in the ligand-binding region rather than global protein motion. For two aligned candidate conformations $R_{s,a}$ and $R_{s,b}$, the pocket-focused structural distance was defined as

$$d_s\left(R_{s,a}, R_{s,b}\right) = \sqrt{\frac{1}{|\mathcal{P}_s|}\sum_{r\in\mathcal{P}_s} \|\mathbf{r}_{a,r} - \mathbf{r}_{b,r}\|_2^2},$$

where $\mathcal{P}_s$ denotes the predefined set of binding-site and state-sensitive residues. This metric emphasizes local structural differences capable of affecting steric complementarity, hydrogen-bond geometry, and other protein–ligand interactions.

The first representative was selected as the ensemble medoid:

$$\tilde{R}_{s,1} = \arg\min_{R_{s,a}\in\mathcal{C}_s} \frac{1}{N_s - 1}\sum_{\substack{b=1\\ b\neq a}}^{N_s} d_s\left(R_{s,a}, R_{s,b}\right).$$

Because the medoid is actual molecular dynamics snapshot rather than an arithmetic coordinate average, it can be used directly for receptor preparation and docking. Its central position represents the geometrically dense region of the sampled pocket ensemble.

Additional representatives were selected through a greedy maximum–minimum-distance procedure:

$$\tilde{R}_{s,t} = \arg \max_{\substack{R\in\mathcal{C}_s\\ R\notin\mathcal{A}_{s,t-1}}} \min_{\tilde{R}\in\mathcal{A}_{s,t-1}} d_s\left(R, \tilde{R}\right).$$

The medoid captures the central pocket geometry, while subsequent selections extend coverage toward conformational regions that are not represented by the structures already selected. Selection was terminated when the intended number of representatives was reached or when all remaining candidate conformations were within the predefined structural-distance threshold.

For each candidate conformation, its distance to the nearest selected representative was defined as

$$e_s(R) = \min_{\tilde{R} \in \mathcal{A}_s} d_s\big(R, \tilde{R}\big).$$

Structural coverage at tolerance $\delta$ was calculated as

$$\mathrm{Coverage}_s(\delta) = \frac{1}{N_s} \sum_{R \in \mathcal{C}_s} \mathbb{I}\,[e_s(R) \leq \delta],$$

where $\mathbb{I}$ denotes the indicator function. The selected receptor subset was therefore interpreted as a structural-coverage representation of the pocket conformations sampled during the analyzed trajectory, rather than as an exhaustive description of the complete thermodynamic landscape.

Each standardized ligand was docked independently against every selected receptor conformation. Docking was used to generate plausible protein–ligand complexes, while docking scores were retained as auxiliary features rather than being interpreted as direct functional predictions. This allowed SAFC to distinguish complexes with similar docking scores but different contact or geometric arrangements.

State-wise averaging prevented the ligand-level prediction from being determined by a single unusually favorable or unfavorable pose. Instead, it emphasized interaction features reproduced across structurally distinct conformations belonging to the same receptor state. Concatenating the resulting state-level representations preserved the functional-state origin of the structural evidence and allowed the downstream classifier to learn differential relationships between receptor states.

The transfer-learning classifier mapped the ligand-level representation to a functional probability through nonlinear transformations followed by a sigmoid output. Model parameters were optimized using binary cross-entropy:

$$\mathcal{L}_{\mathrm{BCE}} = -\frac{1}{N} \sum_{i=1}^{N} [y_i \log \hat{p}_i + (1 - y_i) \log(1 - \hat{p}_i)].$$

Moderate resampling was applied only to the training subset to reduce majority-class dominance. Validation and test subsets retained their original class distributions so that the reported metrics reflected performance under the observed data distribution.

For probability threshold $\tau$, the predicted class was defined as

$$\hat{y}_i(\tau) = \begin{cases} 1, & \hat{p}_i \geq \tau, \\ 0, & \hat{p}_i < \tau. \end{cases}$$

A threshold of 0.5 was used for standard binary classification, while additional thresholds were evaluated to characterize the trade-off between functional-compound recovery and high-confidence candidate selection.

**Results.** The ZAK activity dataset contained substantially more functional than non-functional compounds. Model configurations were therefore evaluated using specificity and balanced accuracy in addition to accuracy, precision, recall, and F1 score. Balanced accuracy was defined as

$$\text{Balanced Accuracy} = \frac{\text{Sensitivity} + \text{Specificity}}{2},$$

which assigns equal importance to recognition of the functional and non-functional classes.

The standard BCE model achieved high accuracy, recall, and F1 score, but its specificity was 0.0000 and its balanced accuracy was 0.5000, indicating that its apparent performance was driven primarily by prediction of the majority functional class. Ensembling modestly increased specificity to 0.1250 but did not sufficiently correct this bias. BCE with moderate resampling produced the most balanced behavior, achieving precision and recall of 0.9322, specificity of 0.5000, and balanced accuracy of 0.7161. The focal-loss model achieved higher nominal accuracy and F1 score but retained limited specificity. Combining focal loss with resampling produced the opposite failure mode, increasing specificity to 0.8750 while reducing recall to 0.1864. BCE with moderate resampling was therefore selected as the primary training configuration because it provided the most useful compromise between recovery of functional compounds and exclusion of non-functional candidates (Table 1A).

SAFC achieved ROC-AUC values of 0.89 for ZAK and 0.92 for NET. These results demonstrate that the continuous SAFC score retained useful ranking information across two structurally and mechanistically distinct target systems. The cross-target results support the transferability of the state-aware interaction framework, while also indicating that target-specific operating thresholds remain necessary. The NET feature-ablation analysis examined whether model performance was driven exclusively by ligand chemical structure or also benefited from receptor-conditioned structural information (Table 1B). Ligand fingerprints provided a strong baseline, with a balanced accuracy of 0.8417. Adding docking-derived score and geometric descriptors increased balanced accuracy to 0.8527 and improved both recall and specificity, indicating that structural features contributed information beyond ligand identity alone.

Contact descriptors did not improve performance when combined with ligand features without score and geometric information, suggesting that isolated contacts may be ambiguous without their associated pose context. The complete representation combining ligand, score/geometric, and contact features achieved the highest balanced accuracy of 0.8532, together with the highest specificity and precision. Score and geometric descriptors therefore provided the clearest independent structural contribution, whereas contact features were most informative as complementary evidence within the complete representation.

Threshold-dependent evaluation showed that the continuous SAFC score could support different candidate-selection objectives (Table 1C). At a NET threshold of 0.61, SAFC achieved precision of 0.98, specificity of 0.90, recall of 0.74, and balanced accuracy of 0.82. Increasing the threshold to 0.72 raised specificity to 0.95 but reduced recall to 0.50. At the most stringent threshold of 0.78, specificity reached 0.98 while recall decreased to 0.34. The monotonic increase in specificity and corresponding decrease in recall demonstrate a coherent candidate-enrichment trade-off. A threshold near 0.61 therefore provides a balanced high-confidence operating point for NET, whereas more stringent thresholds may be applied when candidate purity is prioritized over broad recovery. These thresholds represent target-specific operating points rather than universal cutoffs.

False-positive predictions were concentrated primarily among experimentally non-functional compounds with active-like chemotypes. These errors occurred in the more challenging region of chemical space where structurally similar molecules exhibit divergent experimental activities. False-negative predictions may reflect failure to recover the relevant docking pose, incomplete representation of an important receptor microstate, or functional mechanisms not fully described by the selected pocket-level structural features. The concentration of errors among chemically or mechanistically ambiguous compounds suggests that model errors were not distributed indiscriminately across unrelated chemical space.

The trained ZAK model was subsequently applied to AI-generated inhibitor candidates. Each candidate was evaluated across the selected receptor conformations and assigned a continuous SAFC score. The score was used both for binary functional prioritization and for ranking candidates according to model confidence. High-scoring compounds were considered more consistent with the chemical and receptor-conditioned interaction patterns of experimentally functional molecules. These outputs represent computational prioritization rather than direct experimental confirmation.

**Discussion.** The present results support SAFC as a state-aware functional-prioritization framework rather than a conventional docking-score classifier. The ZAK model-selection analysis illustrates why class-sensitive metrics are necessary for functional datasets with substantial label imbalance. Although several configurations achieved high nominal accuracy and F1 scores, these values concealed poor recognition of non-functional compounds. Moderate resampling with binary cross-entropy produced the most useful balance between sensitivity and specificity, indicating that appropriate treatment of class imbalance was essential to the practical behavior of the classifier.

The NET ablation analysis further showed that the model did not rely exclusively on ligand chemical identity. Ligand fingerprints provided a strong baseline, but docking-derived score and geometric descriptors improved discrimination beyond this baseline. Contact descriptors were less informative in isolation but contributed complementary information when combined with ligand and pose-level features. This pattern is consistent with the design of SAFC: an individual contact may be ambiguous without its geometric context, whereas the integrated representation can distinguish interaction arrangements that are not apparent from ligand structure alone. The ROC-AUC values obtained for ZAK and NET indicate that the continuous SAFC score preserved informative ranking ability across two distinct target systems. In addition, the systematic increase in specificity and corresponding decrease in recall at stricter NET thresholds demonstrate that the score supports predictable operating-point selection. SAFC can therefore be used either to retain a broader set of potentially functional candidates or to identify a smaller, more highly enriched subset when downstream experimental capacity is limited.

The observed errors also define the current boundary of the method. False-positive predictions were concentrated primarily among inactive compounds with active-like chemotypes, while false negatives may reflect docking-pose failure, insufficient representation of relevant receptor microstates, or functional mechanisms not fully described by pocket-level structural features. These cases are intrinsically challenging because favorable binding patterns and chemical similarity do not always translate directly into the measured functional endpoint. SAFC reduces this ambiguity by incorporating receptor-state and conformational information, but it does not replace experimental functional validation.

Overall, SAFC provides a unified computational strategy for distinguishing potential functional activity from favorable binding alone. By integrating receptor functional states, MD-derived conformational diversity, ensemble docking, complex-level interaction representations, and target-specific activity supervision, the framework evaluates whether ligand–receptor compatibility is reproducible across biologically relevant structural contexts. The combined model-selection, cross-target, ablation, threshold, and error analyses provide complementary evidence that SAFC can reduce large molecular candidate sets to smaller subsets enriched for predicted functional activity.

| Parameter | Setting |
|---|---|
| Benchmark systems | 176 |
| AlignDockBench systems | 61 |
| PoseBusters systems | 40 |
| PLINDER systems | 75 |
| Requested molecules per target | 100 |
| Random seed | 2026 |
| Sampling-box size | 23 Å |
| Beam size | 300 |
| Maximum generation steps | 50 |
| Focal-site threshold | 0.50 |
| Position threshold | 0.25 |
| Element threshold | 0.30 |
| Atom-existence threshold | 0.60 |
| Bond threshold | 0.40 |
| GPU allocation | One NVIDIA A100 GPU |
| CPU allocation | One CPU core per process |
| Execution mode | Sequential target-wise generation |
| Retained outputs | SDF, SMILES, logs and configuration files |
| Intermediate .pt files | Disabled/removed |

**Supplementary Table 1.** Sampling configuration, benchmark composition, and computational resources used for molecular generation across 176 protein-ligand systems.

| Workflow | Generation paradigm | Target completion, n/176 (%) | Recorded runtime/target, min, median [Q1–Q3] | Observed peak VRAM, GiB |
|---|---|---|---|---|
| PFM | Perturbed flow matching | 173 (98.3) | 7.1 [5.0–9.9] | 17.9 |
| DrugFlow | Flow matching with discrete Markov bridges | 163 (92.6) | 5.2 [4.3–6.6] | 12.4 |
| FLOWR | Pocket-aware equivariant flow matching | 164 (93.2) | 1.0 [0.7–1.2] | 78.6 |
| FlowMol† | Unconditional flow matching | 175 (99.4) | 1.3 [1.2–1.3] | 7.5 |
| TargetDiff | Target-aware SE(3)-equivariant diffusion | 174 (98.9) | 27.2 [19.6–43.5] | 21.7 |
| ShEPhERD | SE(3)-equivariant joint diffusion | 165 (93.8) | 36.8 [26.9–65.1] | 65.4 |
| Pocket2Mol | Pocket-conditioned E(3)-equivariant autoregressive generation | 175 (99.4) | 12.5 [9.8–16.0] | 18.4 |
| MolJO | Gradient-guided Bayesian-flow optimization | 175 (99.4) | 43.4 [33.5–62.5] | 39.1 |
| REINVENT4 Mol2Mol | Sequence-based molecular optimization | 114 (64.8) | 2.4 [1.8–3.0] | NR |
| PrexSyn | Decoder-only synthesis transformer | 164 (93.2) | 0.9 [0.9–1.0] | 7.8‡ |
| ReaSyn | Chain-of-reaction pathway generation | 153 (86.9) | 1.9 [1.6–2.4] | NR |
| SynFormer | Synthesis transformer with diffusion-based building-block selection | 164 (93.2) | 0.7 [0.5–1.2] | 7.8‡ |

**Supplementary Table 2.** Target completion, runtime, and observed peak GPU memory across molecular generation workflows. All workflows were configured to generate 100 molecules per target. Target completion was determined from the corresponding workflow execution records and is reported over the full 176-target benchmark; non-completed targets remained in the denominator. PDB 5OHE was retained in the benchmark; for affected workflows, its Fe–N dative coordination bond caused an RDKit ligand-preprocessing failure. Runtime summarizes the available per-target timing records and was analyzed independently of target completion. Observed peak VRAM denotes the maximum recorded device-level GPU-memory use under the evaluated execution setting. †FlowMol is an unconditional, target-agnostic baseline. ‡PrexSyn and SynFormer VRAM values were obtained from run-wide rather than target-specific monitoring. NR, not recorded.

**Fig. S1. QED, SA, and AutoDock Vina score of PFM**

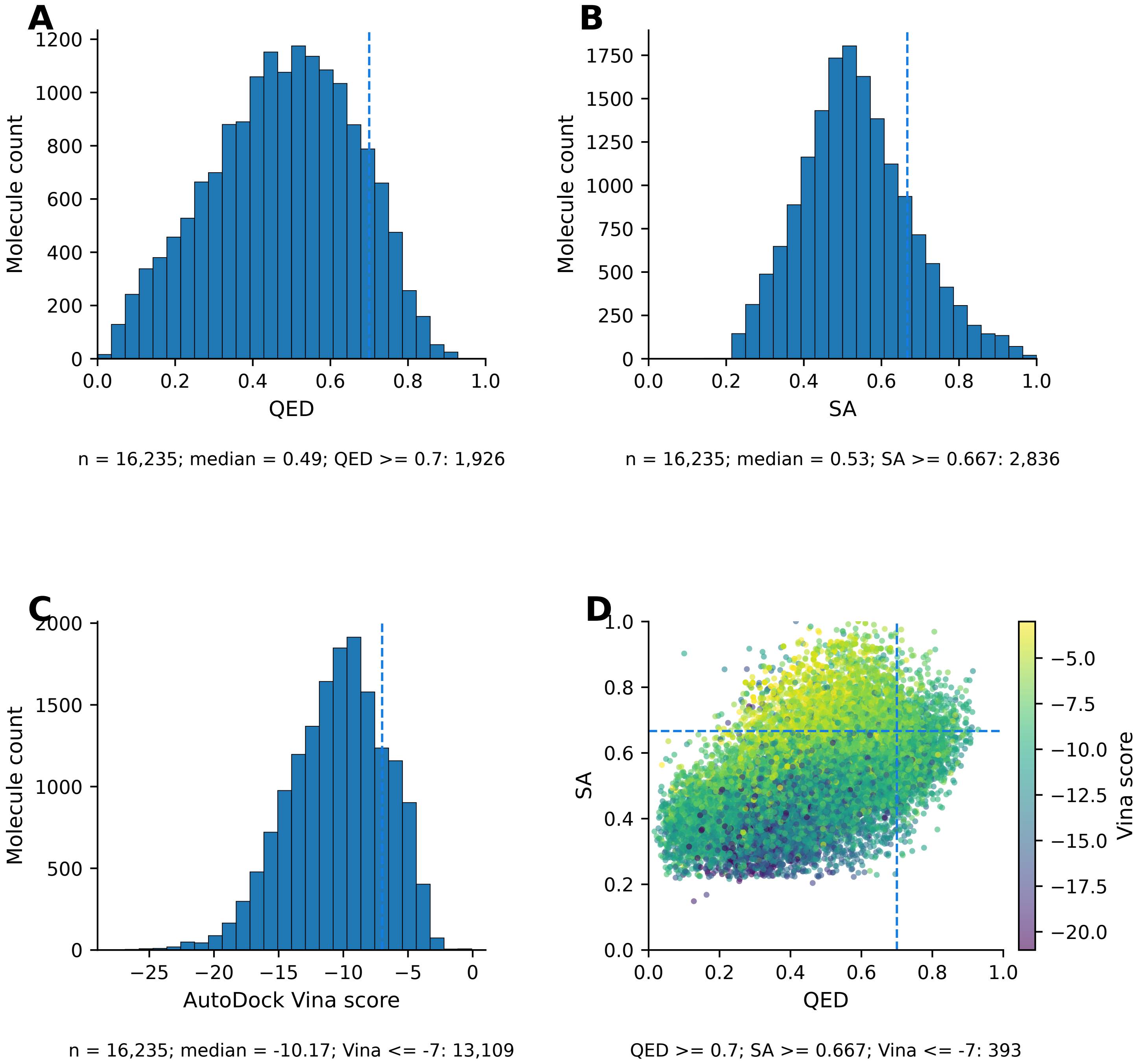

**Fig. S2. QED, SA, and AutoDock Vina score of MolJO**

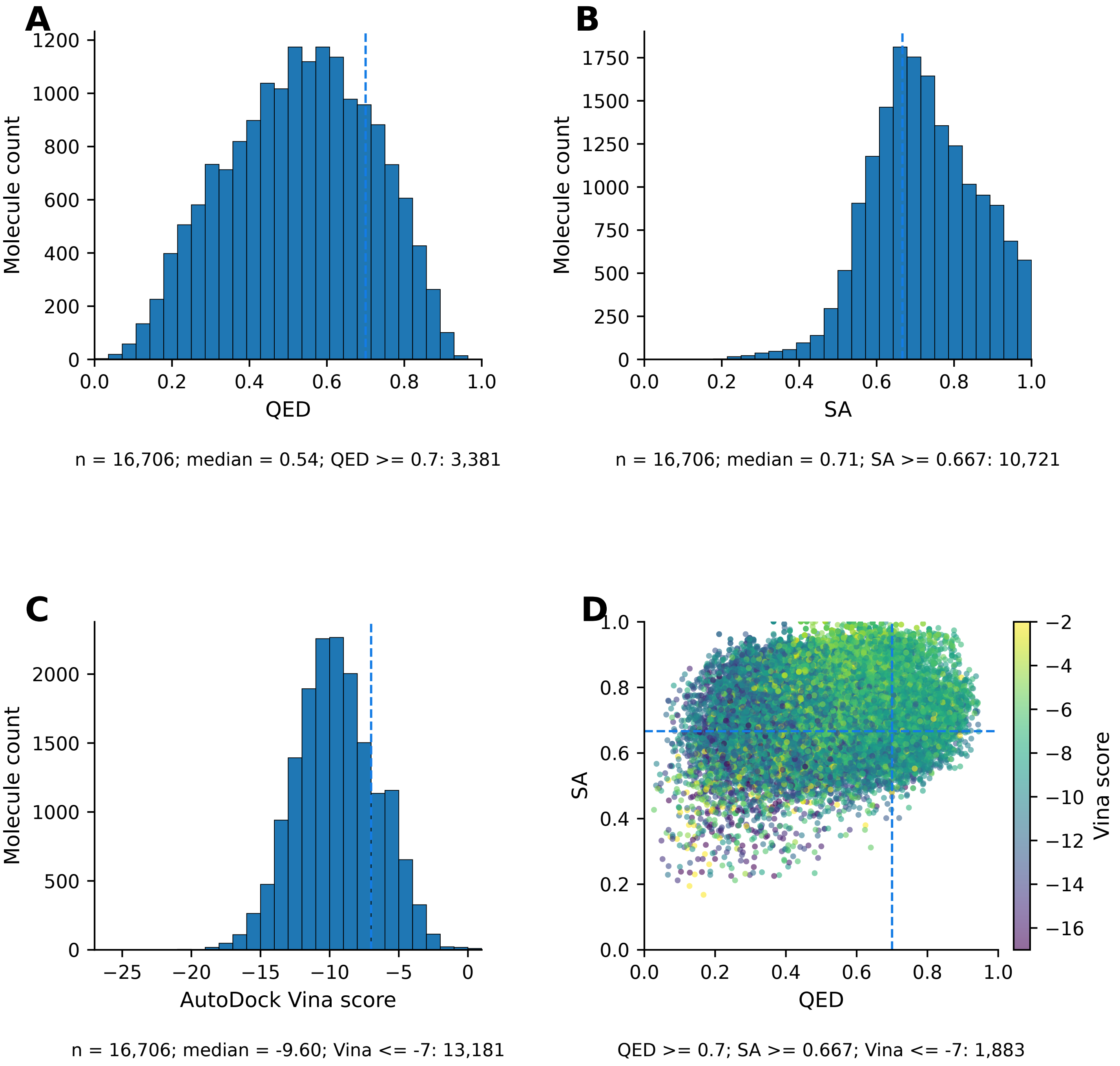

**Fig. S3. QED, SA, and AutoDock Vina score of DrugFlow**

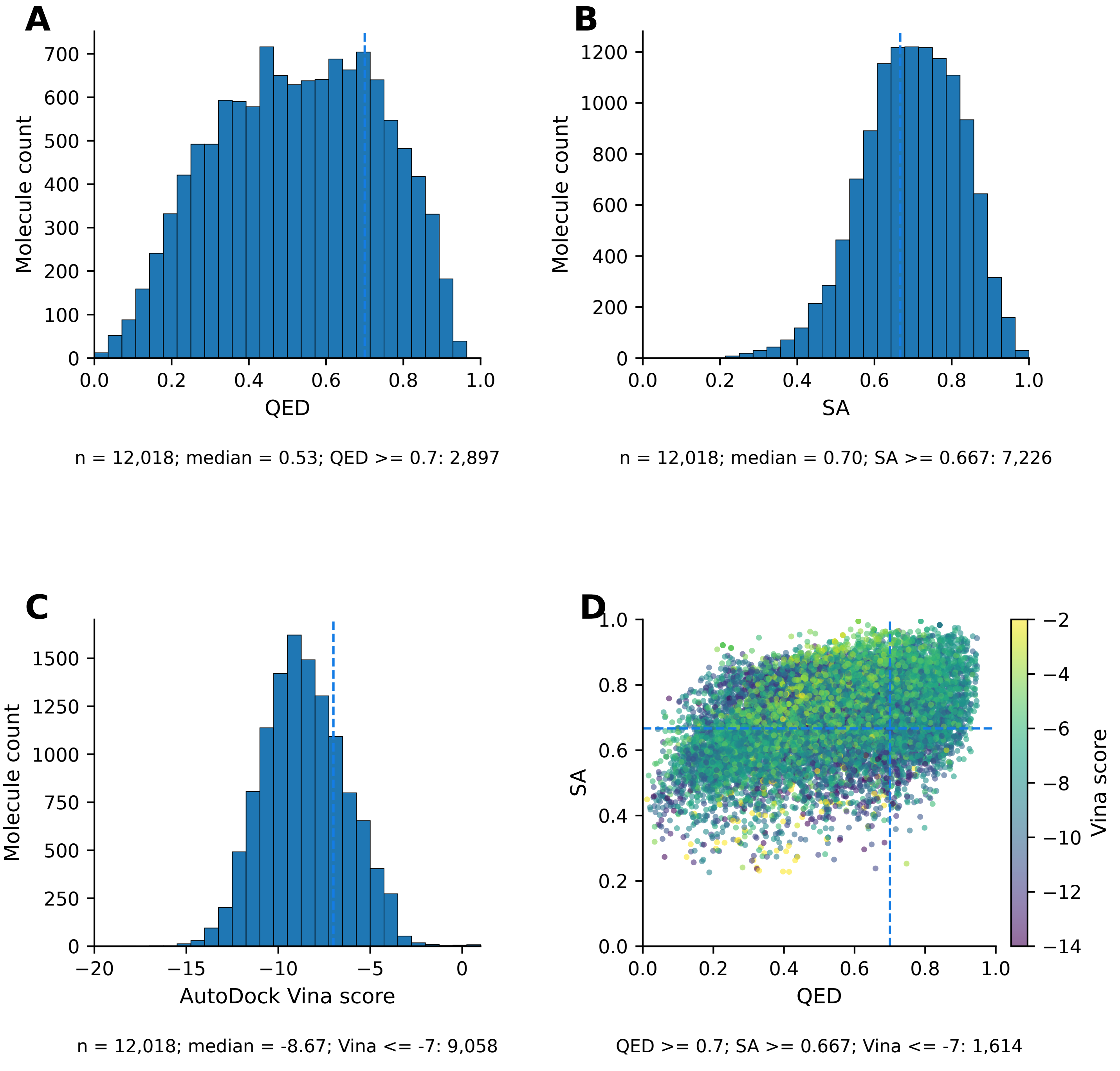

**Fig. S4. QED, SA, and AutoDock Vina score of FlowMol**

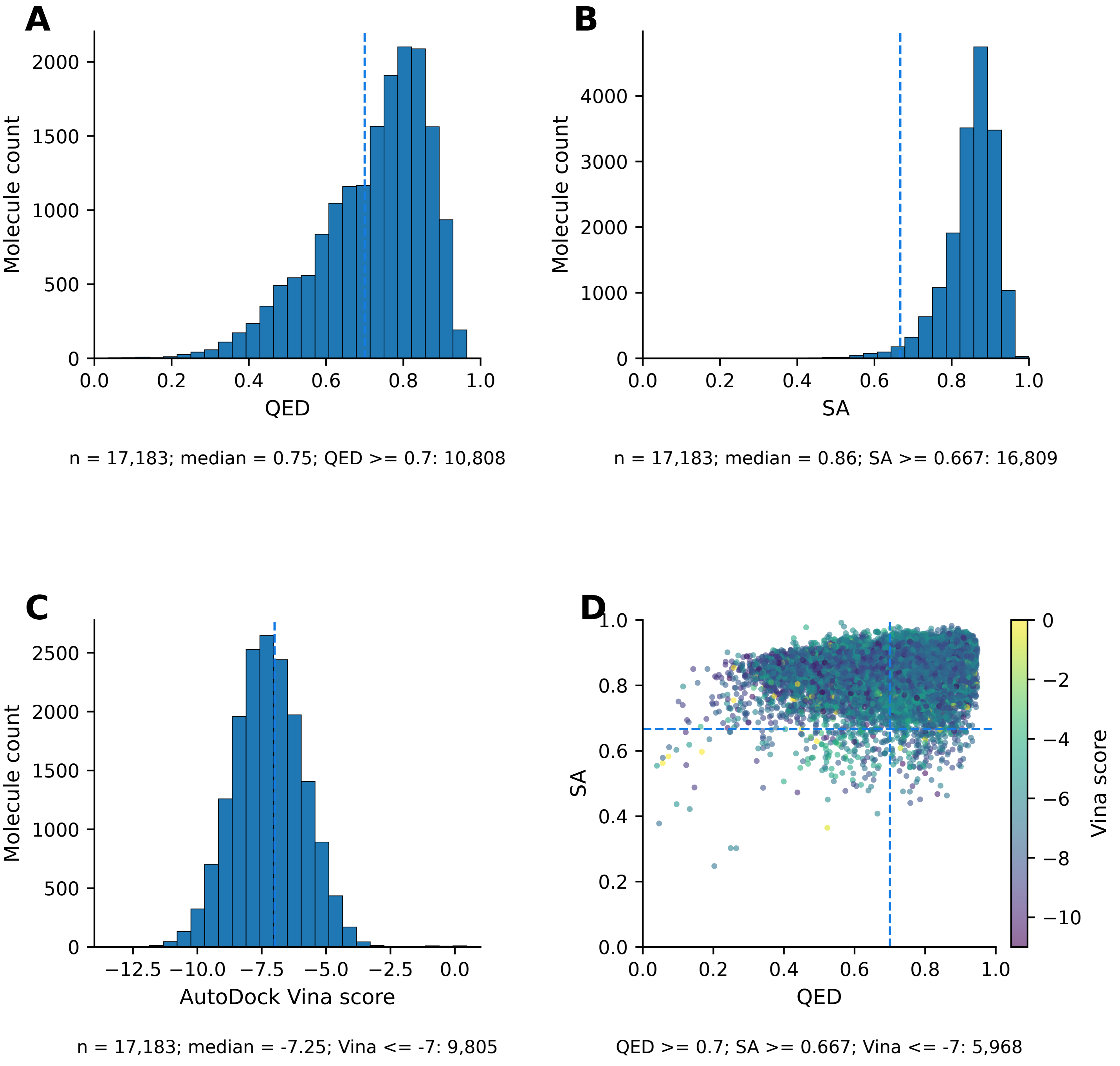

**Fig. S5. QED, SA, and AutoDock Vina score of Pocket2Mol**

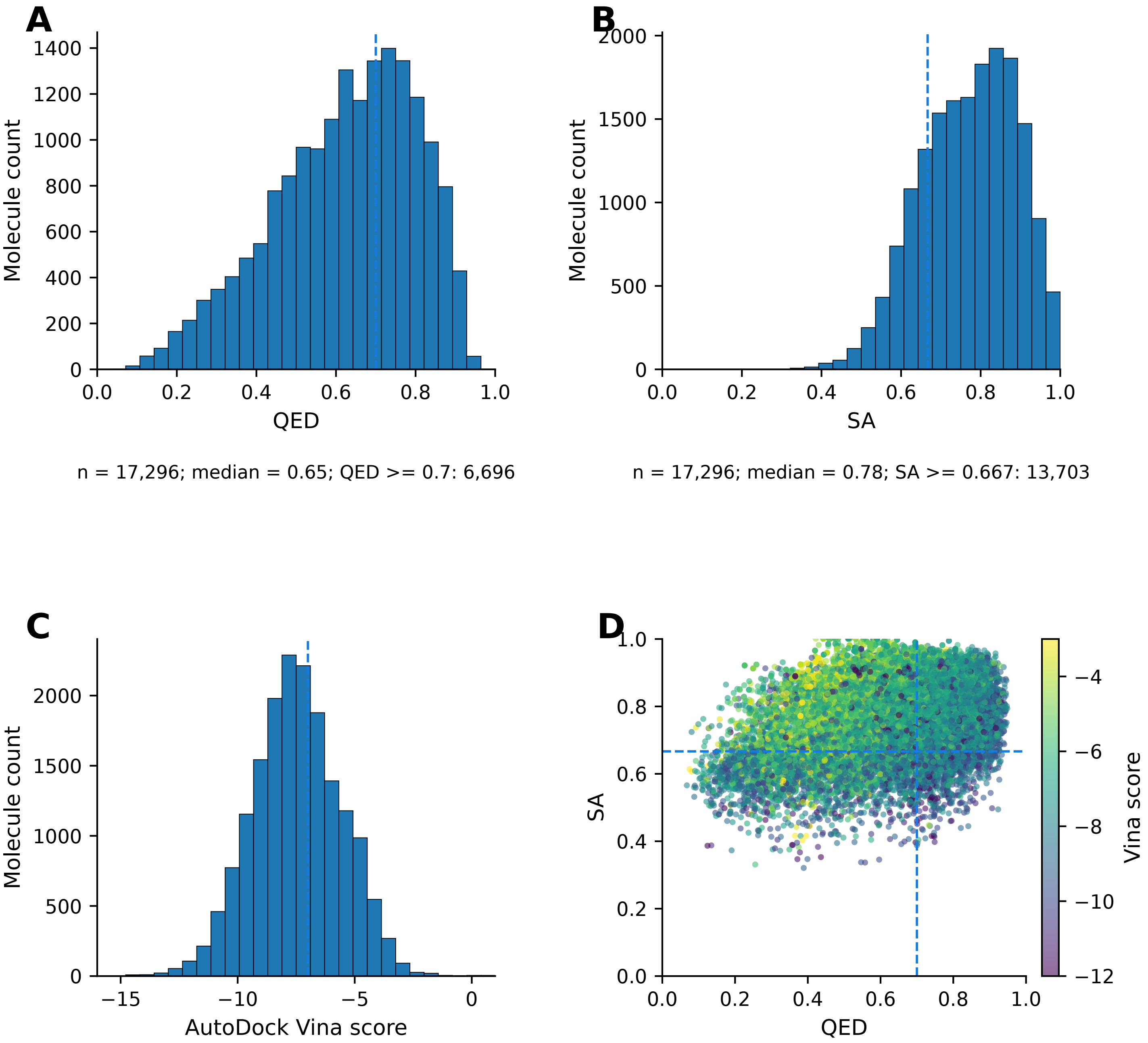


n = 17,296; median = 0.65; QED >= 0.7: 6,696

n = 17,296; median = 0.78; SA >= 0.667: 13,703

n = 17,296; median = -7.50; Vina <= -7: 10,441

QED >= 0.7; SA >= 0.667; Vina <= -7: 4,458

**Fig. S6. QED, SA, and AutoDock Vina score of ShEPhERD**

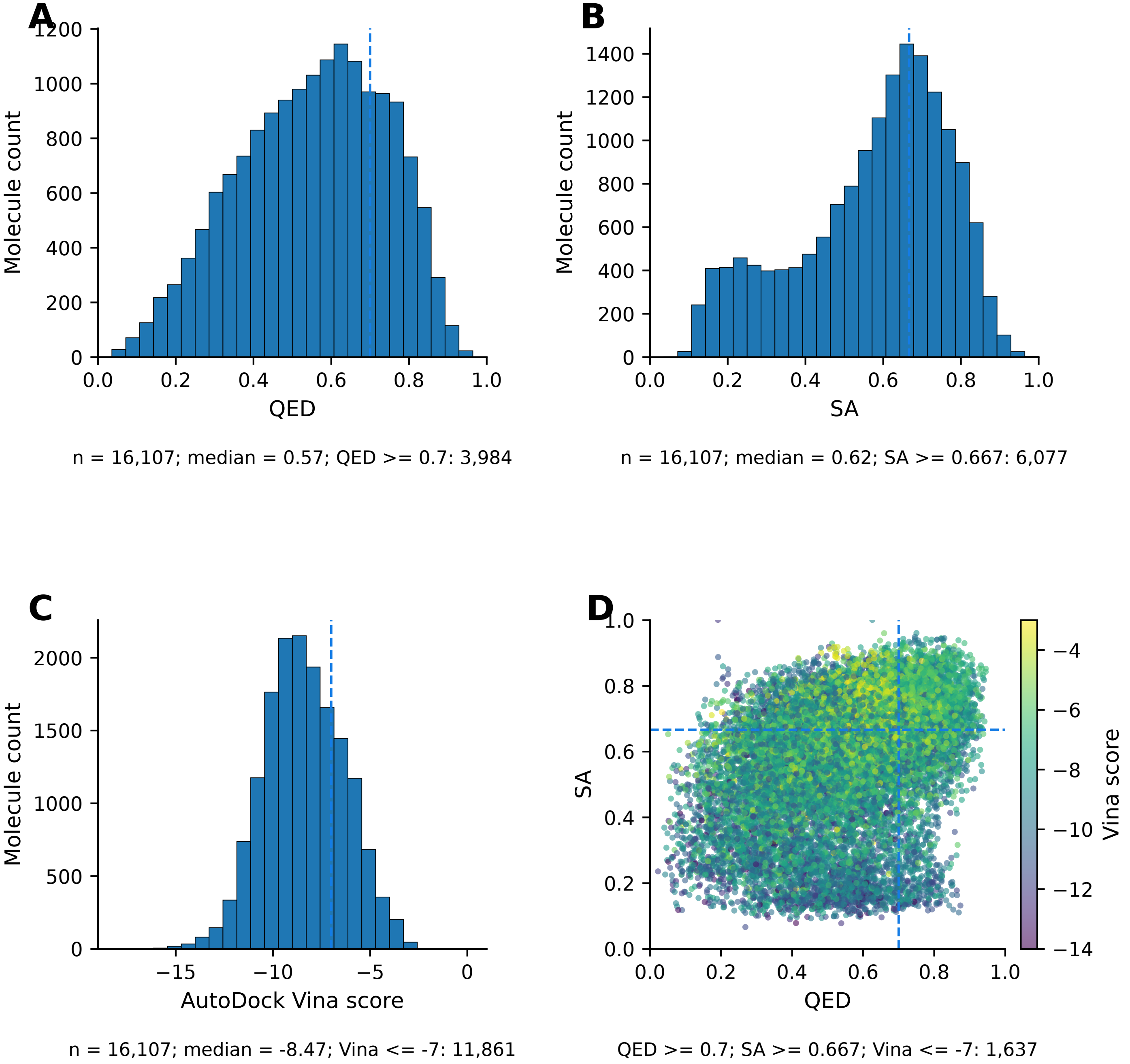

**Fig. S7. QED, SA, and AutoDock Vina score of REINVENT4**

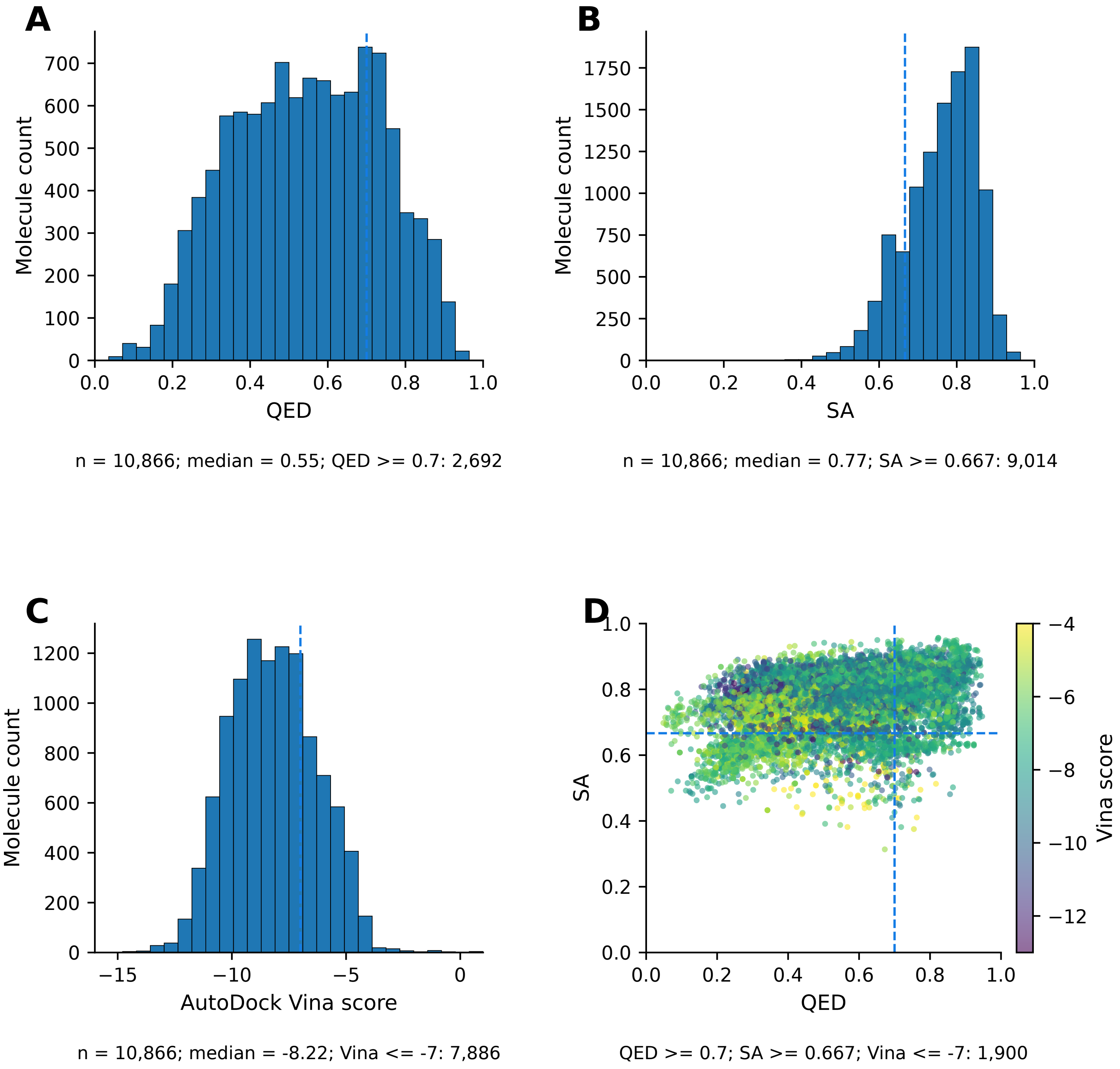

**Fig. S8. QED, SA, and AutoDock Vina score of ReaSyn**

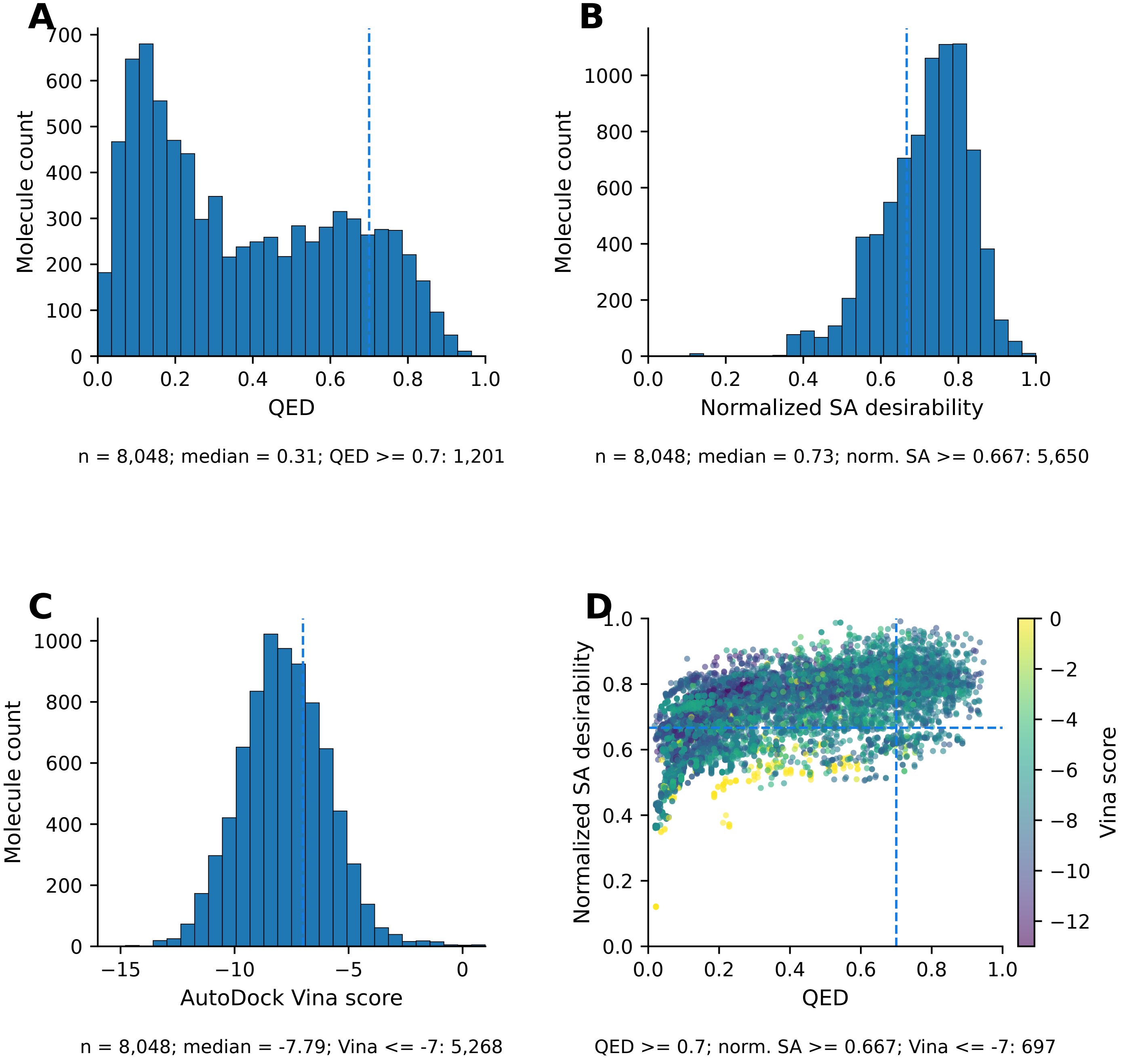

**Fig. S9. QED, SA, and AutoDock Vina score of SynFormer**

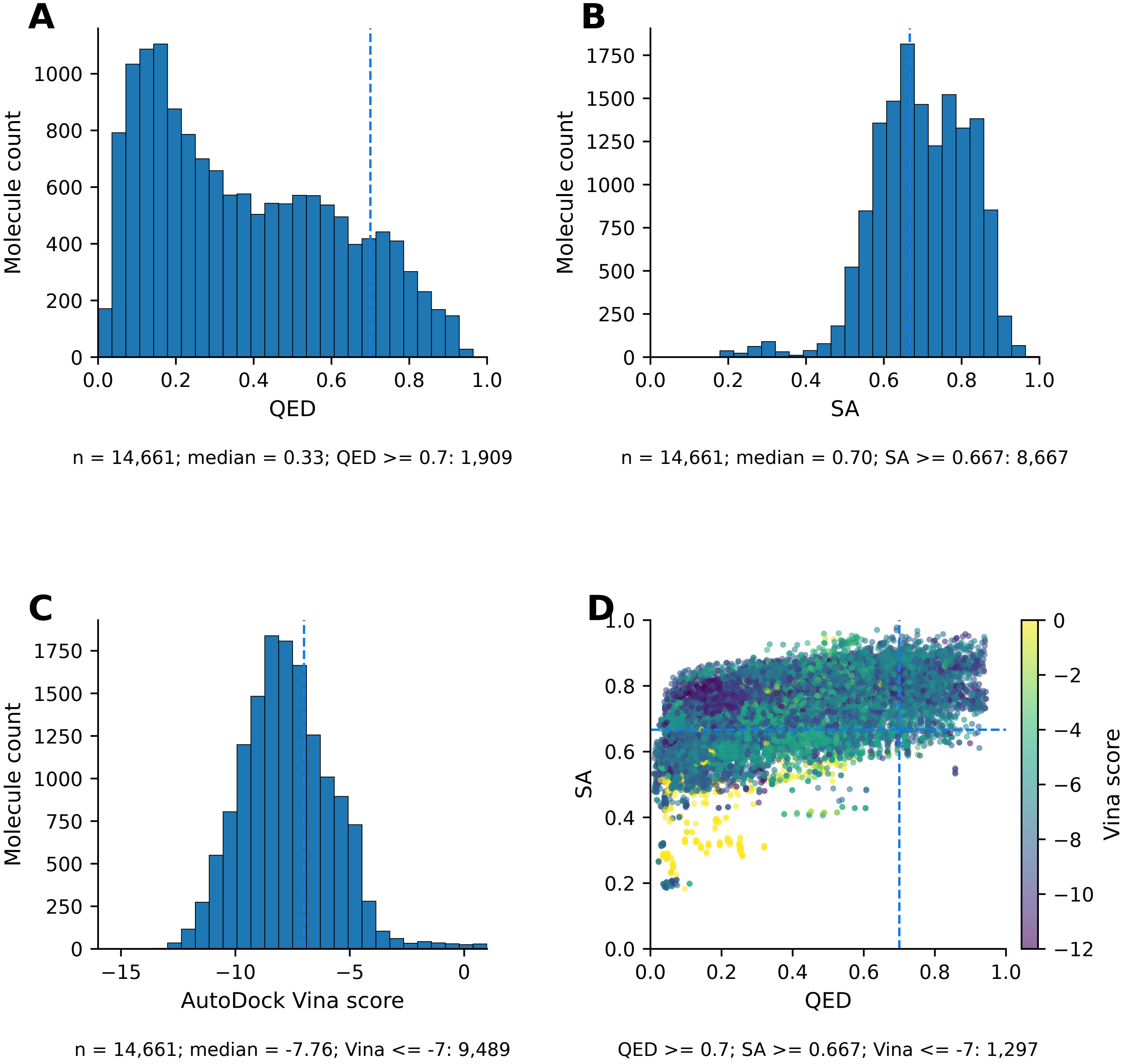

**Fig. S10. QED, SA, and AutoDock Vina score of PrexSyn**

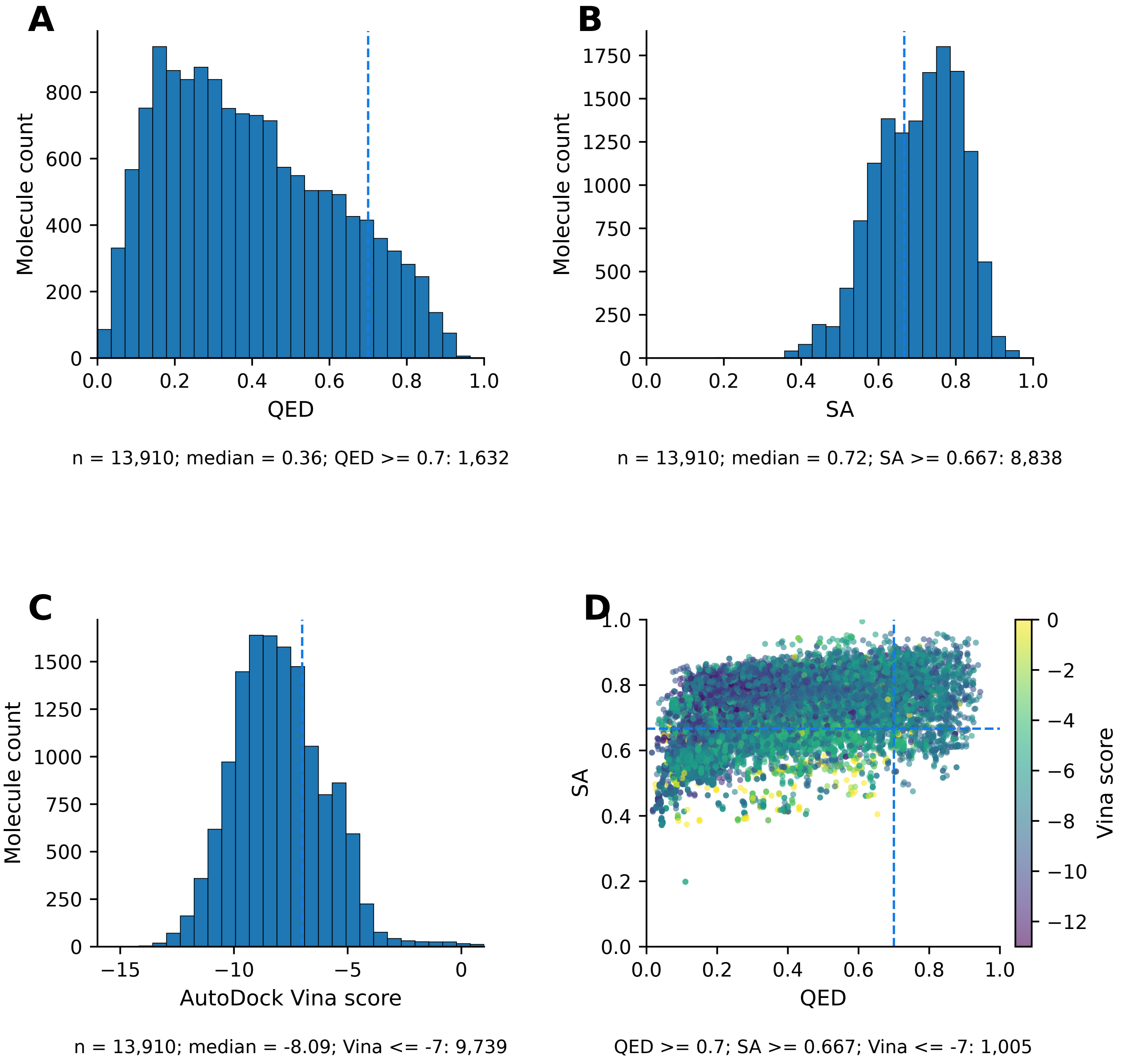

**Fig. S11. QED, SA, and AutoDock Vina score of TargetDiff**

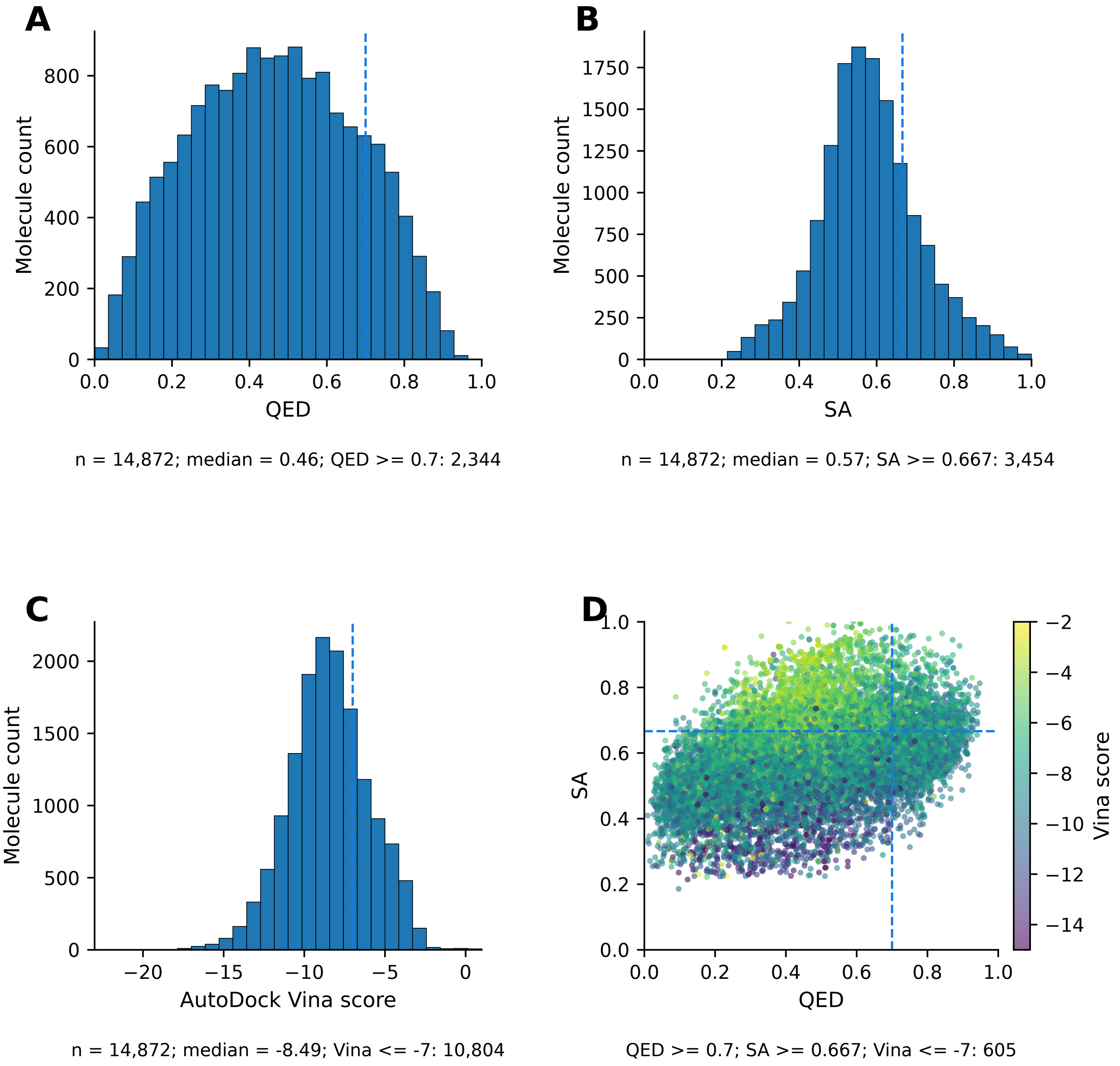

**Fig. S12. QED, SA, and AutoDock Vina score of FLOWR**

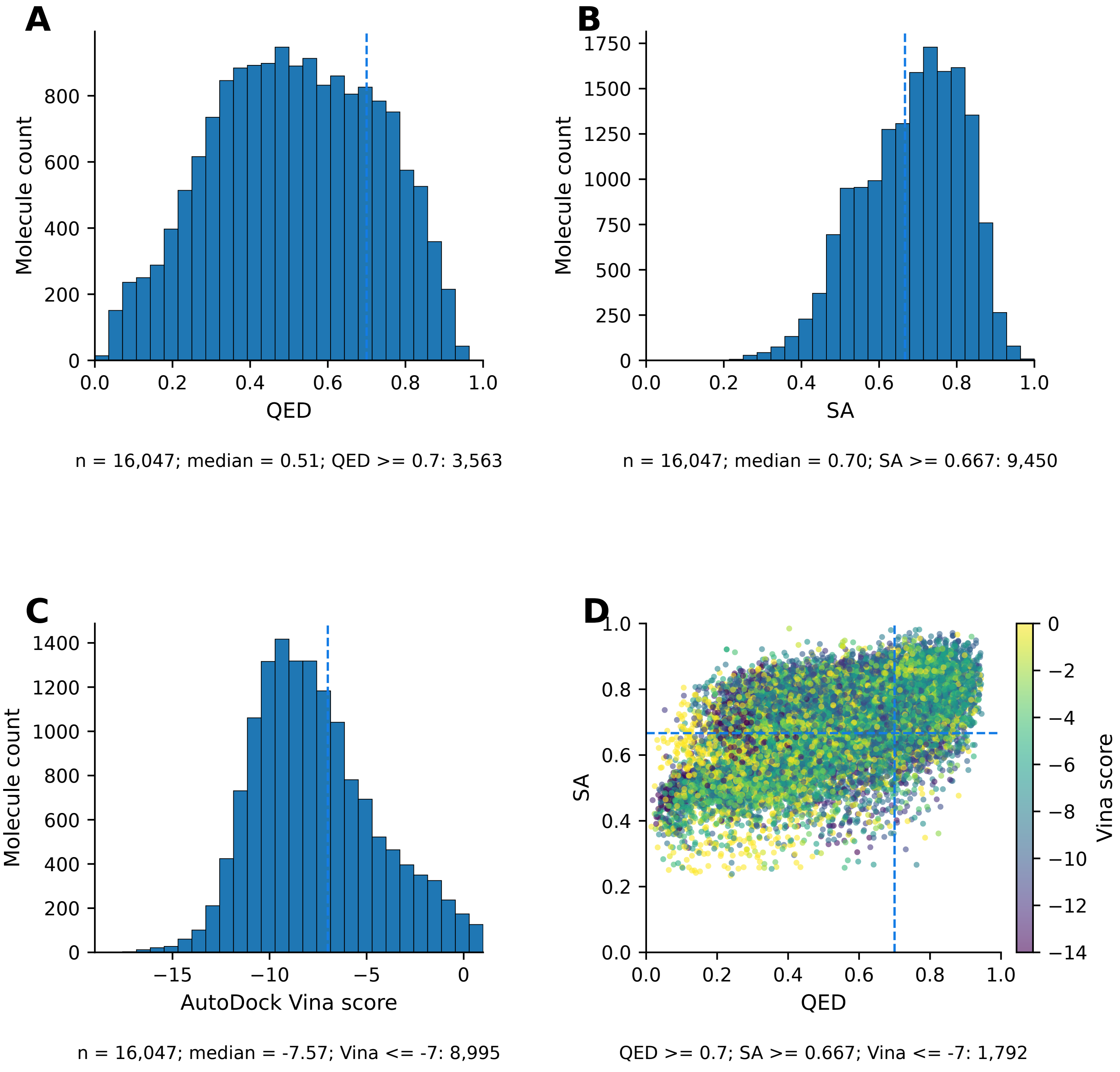

### Fig. S13. ADMET-AI ADME profiling of PFM-generated molecules

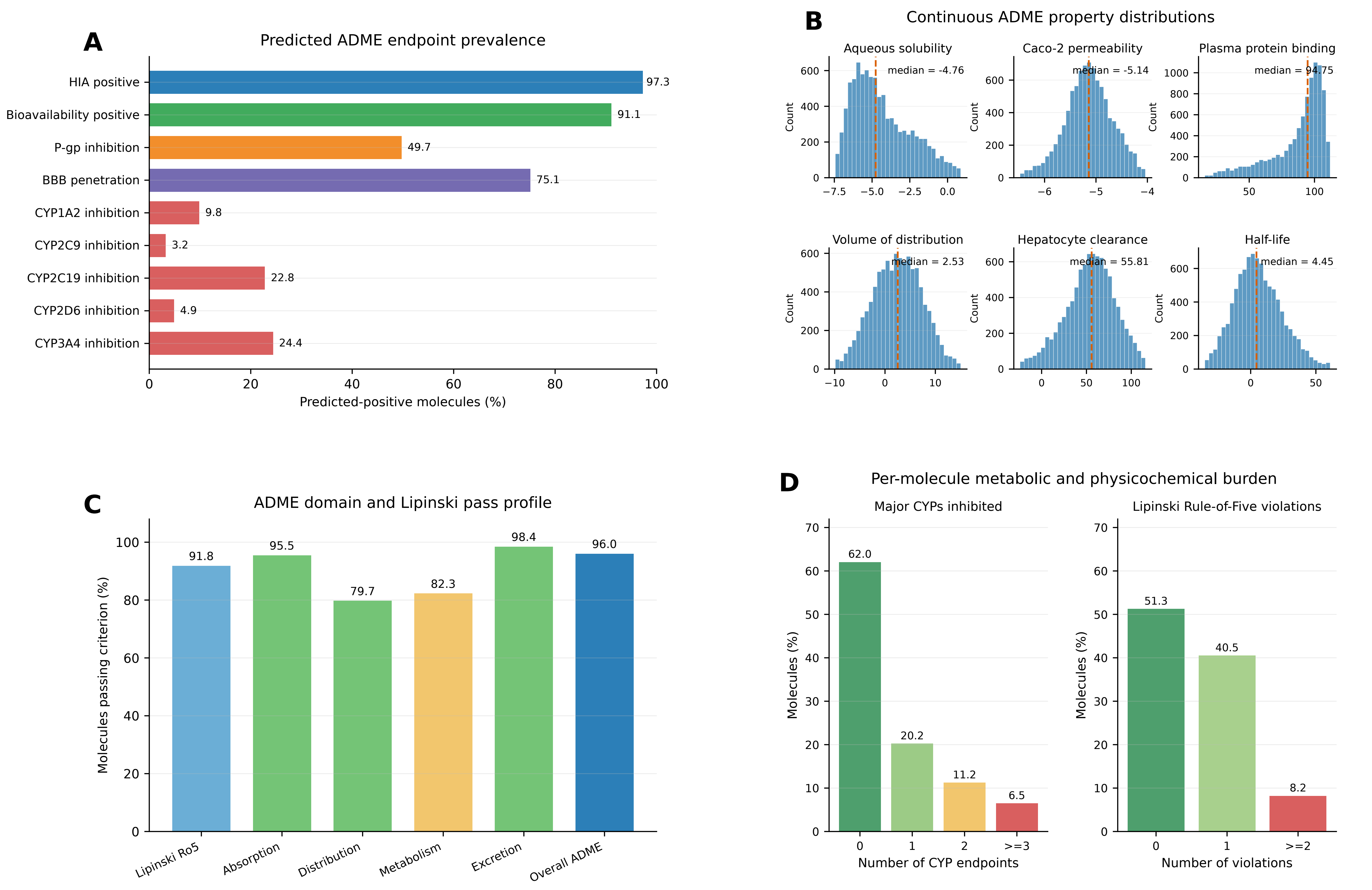


n = 9,449 molecule records (9,377 unique SMILES) across 97 targets; predicted-positive threshold = 0.5.

Distribution and excretion pass bands are comparative pooled 5th-95th percentile ranges.

**Fig. S14. ADMET-AI ADME profiling of MolJO-generated molecules**

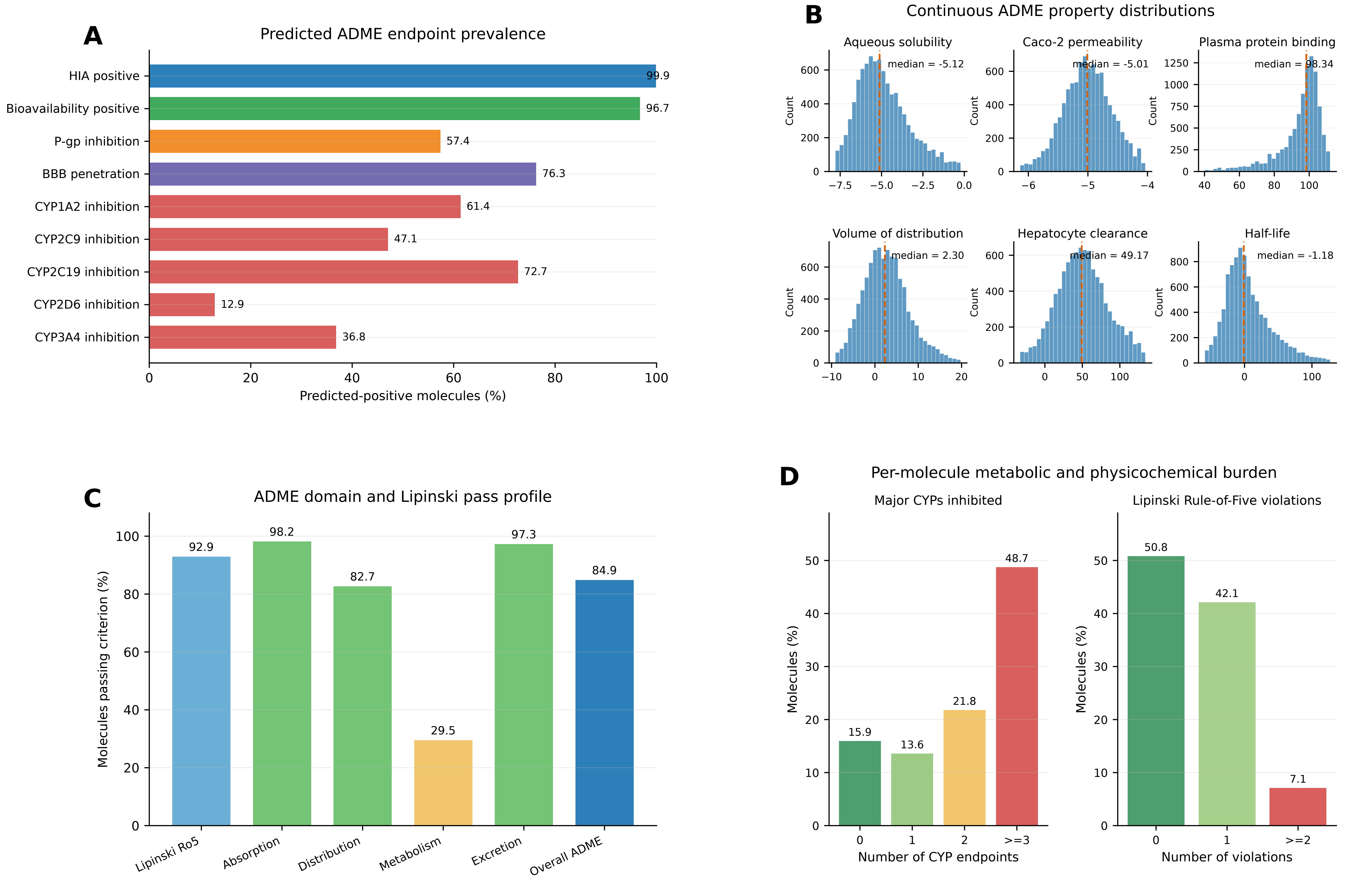


n = 9,689 molecule records (9,256 unique SMILES) across 97 targets; predicted-positive threshold = 0.5.

Distribution and excretion pass bands are comparative pooled 5th-95th percentile ranges.

**Fig. S15. ADMET-AI ADME profiling of DrugFlow-generated molecules**

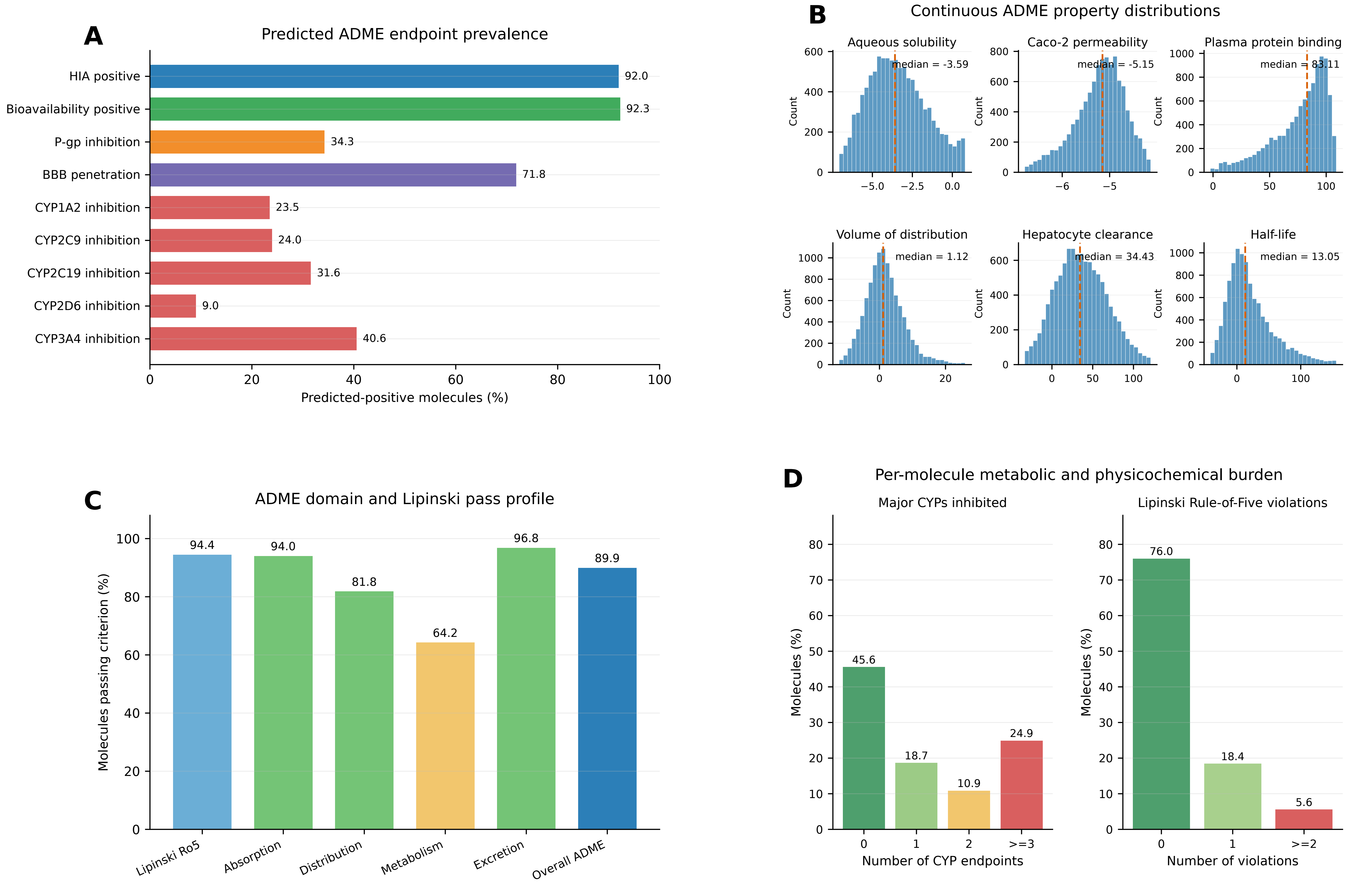


n = 10,641 molecule records (10,488 unique SMILES) across 97 targets; predicted-positive threshold = 0.5.

Distribution and excretion pass bands are comparative pooled 5th-95th percentile ranges.

### Fig. S16. ADMET-AI ADME profiling of FlowMol-generated molecules

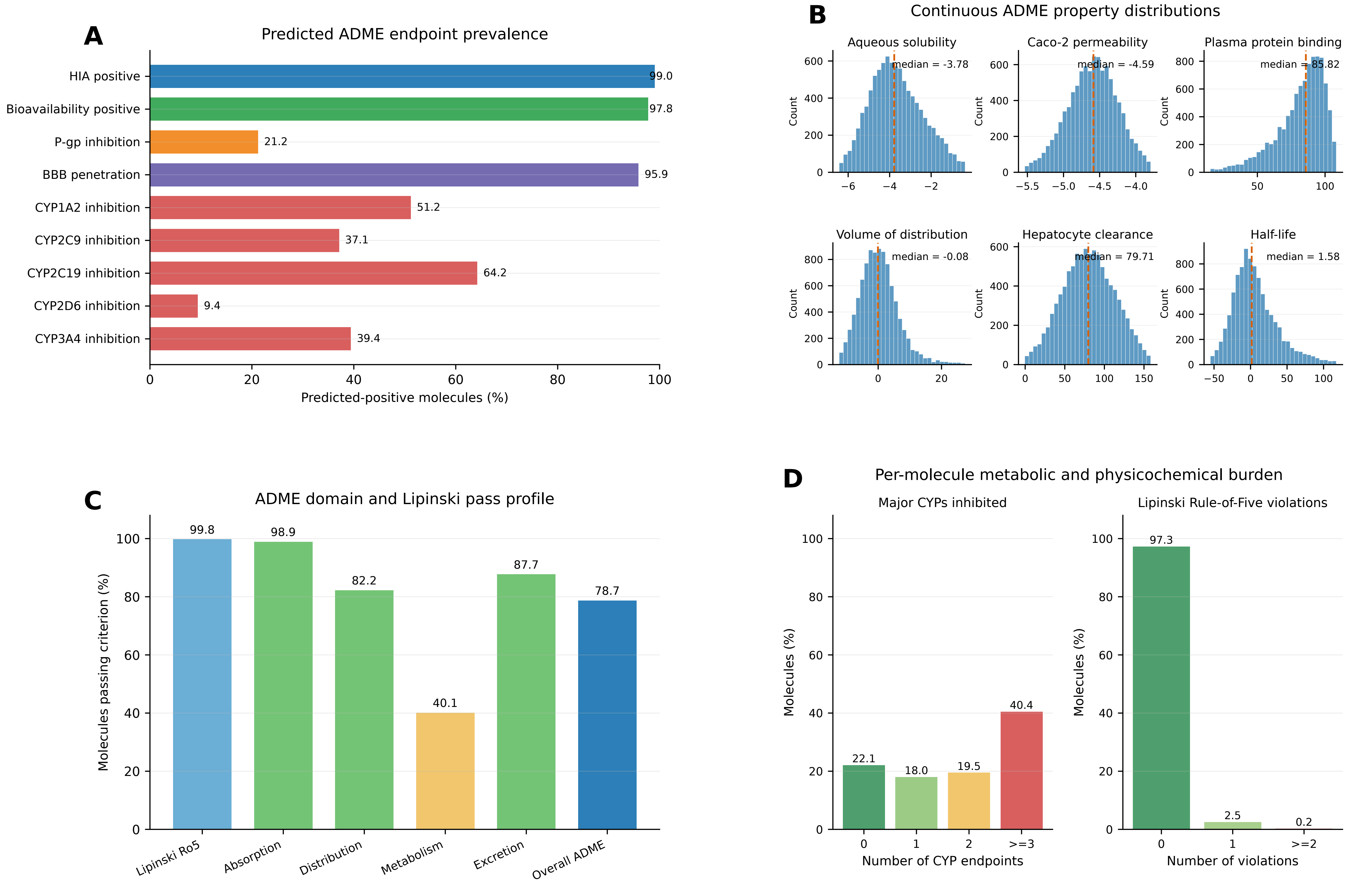


n = 9,699 molecule records (9,653 unique SMILES) across 97 targets; predicted-positive threshold = 0.5.

Distribution and excretion pass bands are comparative pooled 5th-95th percentile ranges.

### Fig. S17. ADMET-AI ADME profiling of Pocket2Mol-generated molecules

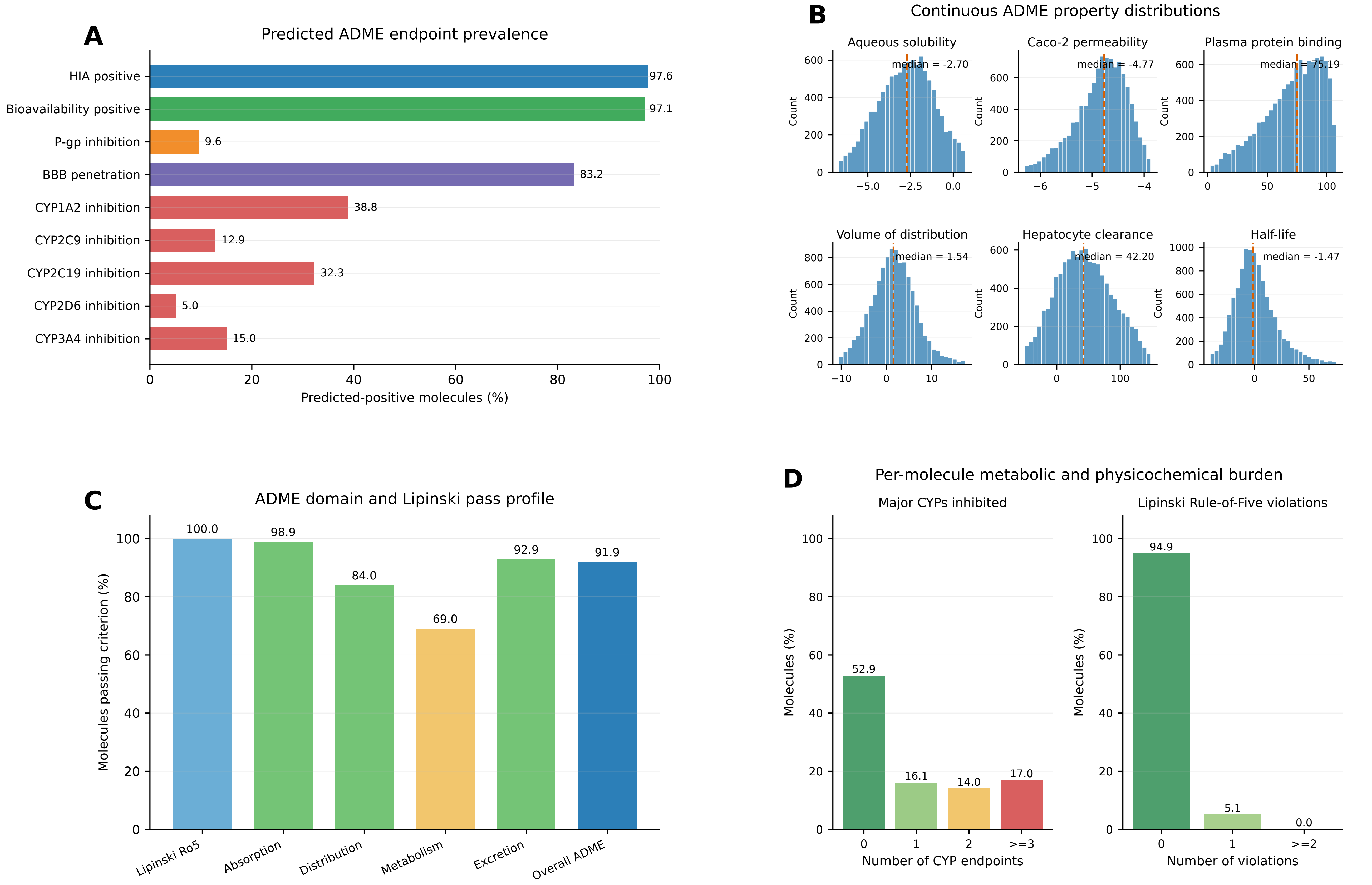


n = 10,749 molecule records (10,385 unique SMILES) across 97 targets; predicted-positive threshold = 0.5.

Distribution and excretion pass bands are comparative pooled 5th-95th percentile ranges.

### Fig. S18. ADMET-AI ADME profiling of ShEPhERD-generated molecules

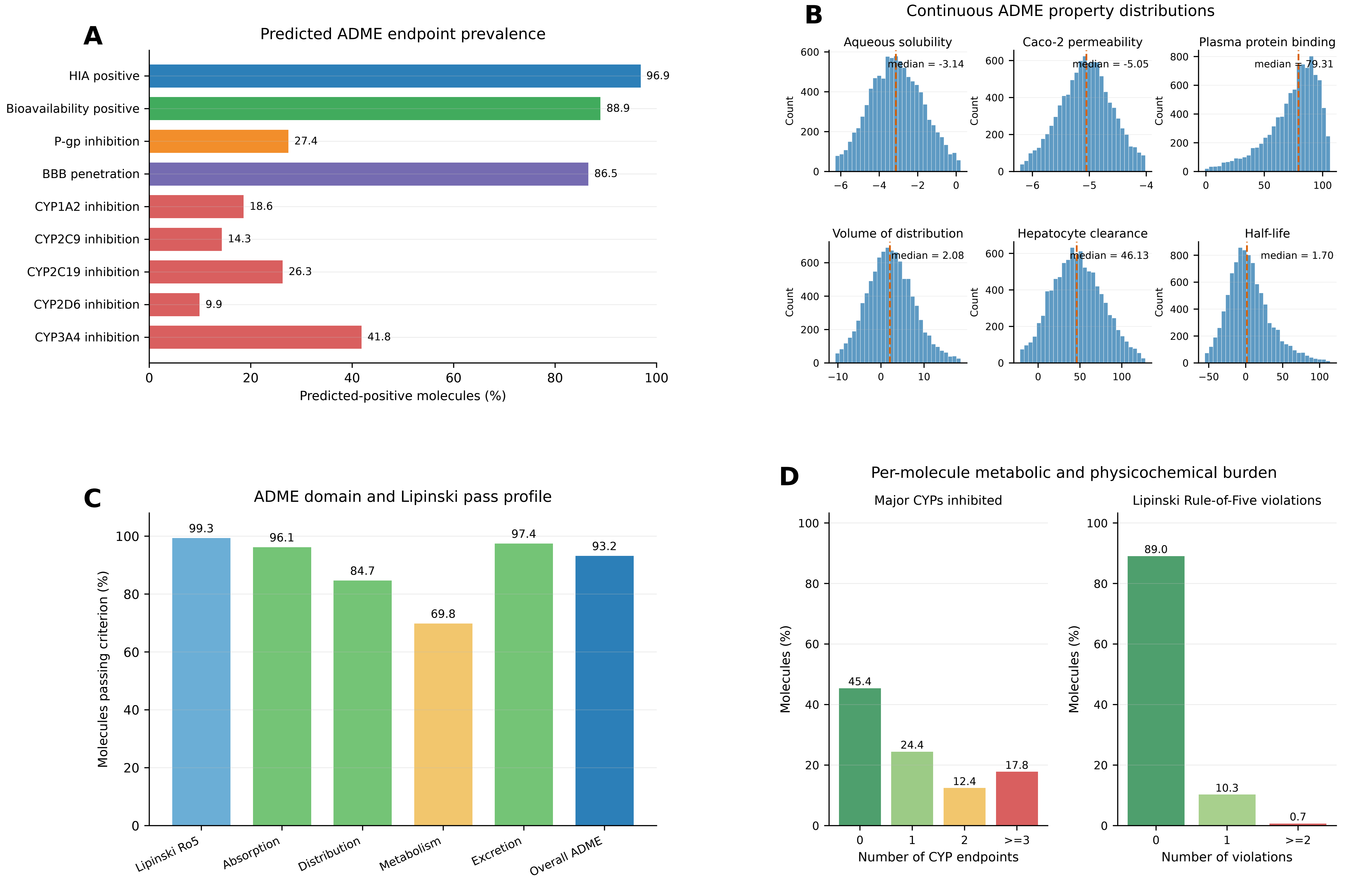


n = 9,609 molecule records (9,608 unique SMILES) across 97 targets; predicted-positive threshold = 0.5.

Distribution and excretion pass bands are comparative pooled 5th-95th percentile ranges.

**Fig. S19. ADMET-AI ADME profiling of REINVENT4-generated molecules**

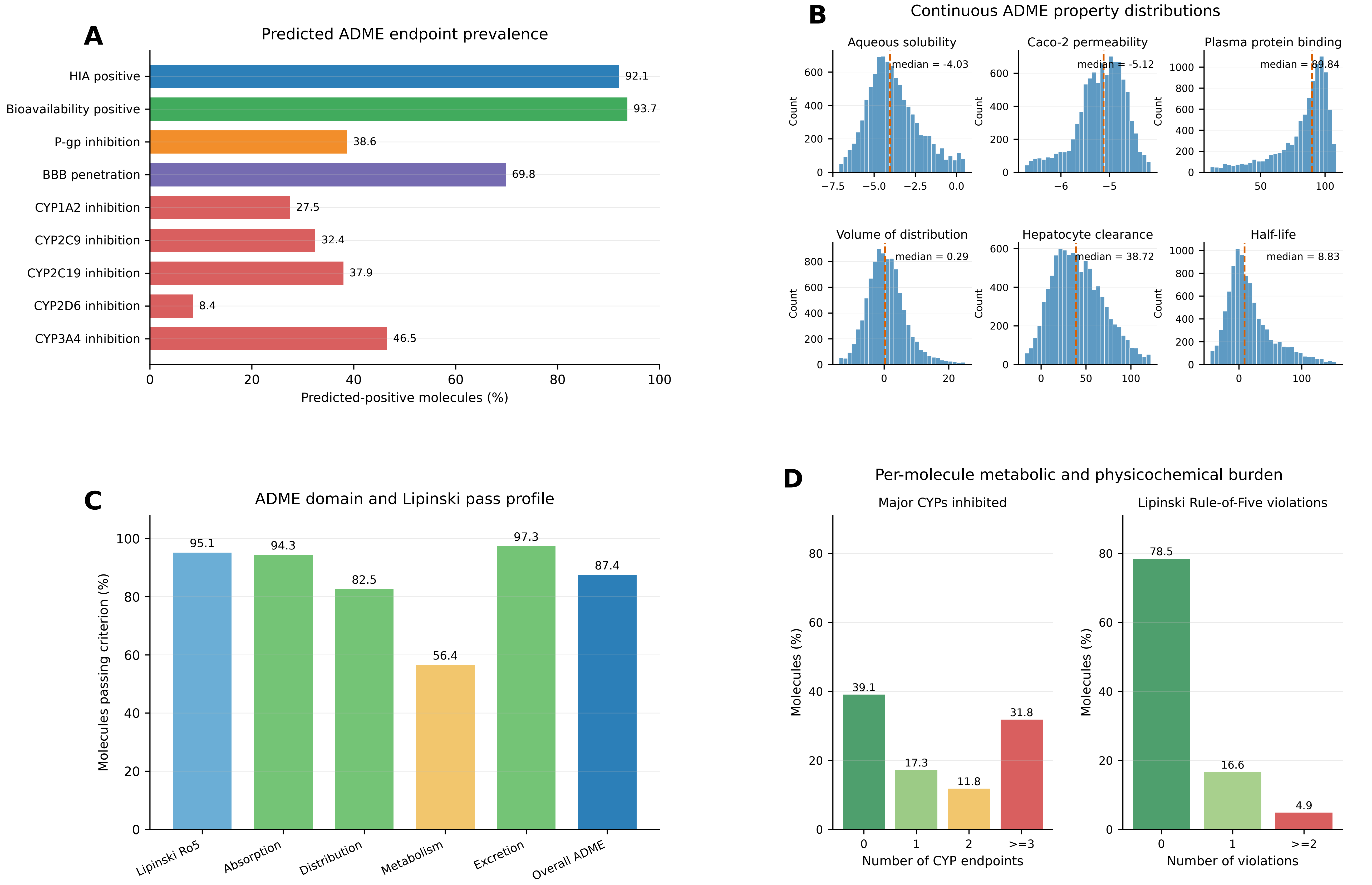


n = 9,499 molecule records (9,185 unique SMILES) across 97 targets; predicted-positive threshold = 0.5.

Distribution and excretion pass bands are comparative pooled 5th-95th percentile ranges.

### Fig. S20. ADMET-AI ADME profiling of ReaSyn-generated molecules

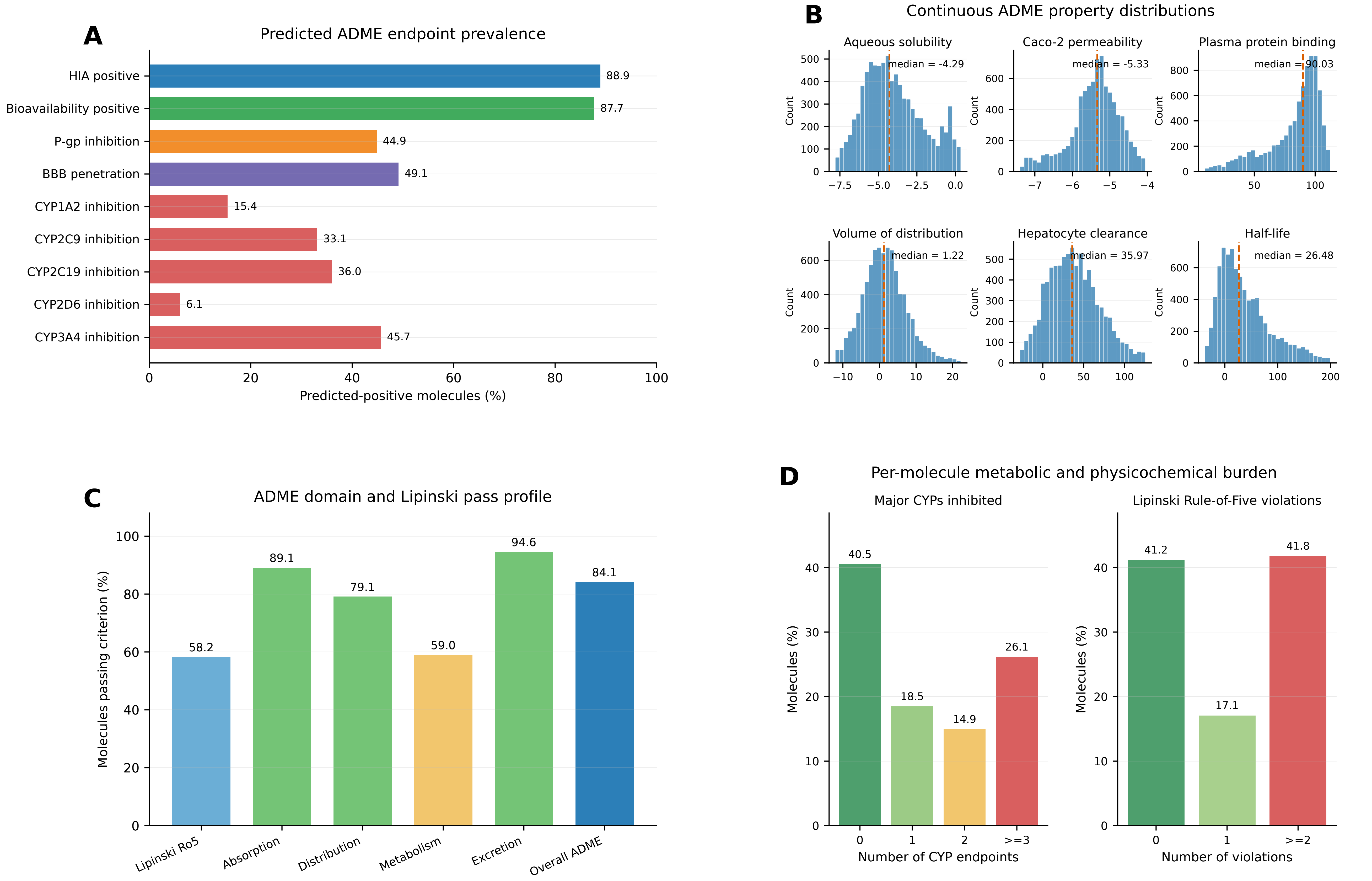


n = 8,526 molecule records (8,453 unique SMILES) across 97 targets; predicted-positive threshold = 0.5.

Distribution and excretion pass bands are comparative pooled 5th-95th percentile ranges.

### Fig. S21. ADMET-AI ADME profiling of SynFormer-generated molecules

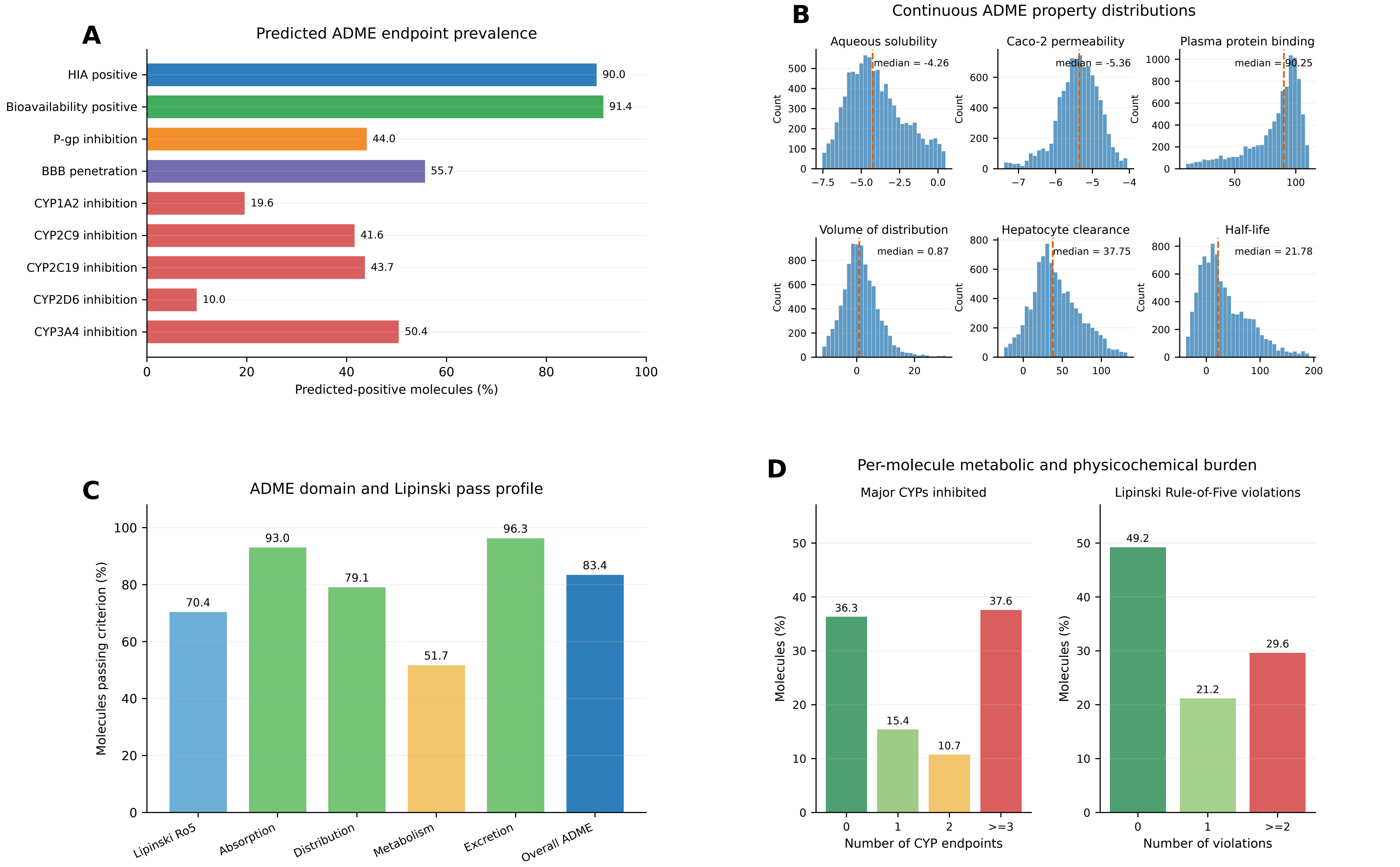


n = 9,105 molecule records (9,028 unique SMILES) across 97 targets; predicted-positive threshold = 0.5.

Distribution and excretion pass bands are comparative pooled 5th-95th percentile ranges.

### Fig. S22. ADMET-AI ADME profiling of PrexSyn-generated molecules

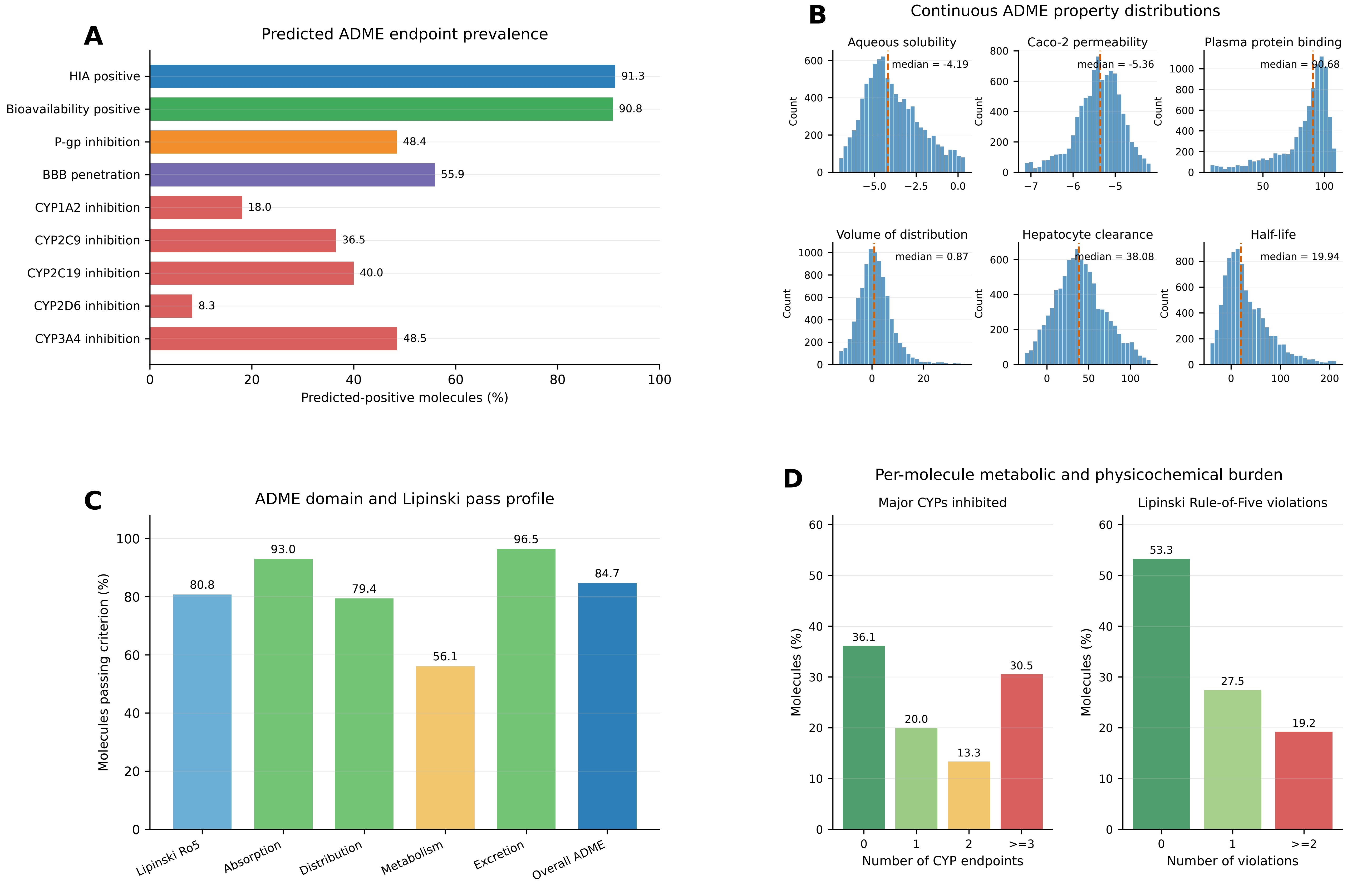


n = 9,065 molecule records (9,006 unique SMILES) across 97 targets; predicted-positive threshold = 0.5.

Distribution and excretion pass bands are comparative pooled 5th-95th percentile ranges.

### Fig. S23. ADMET-AI ADME profiling of TargetDiff-generated molecules

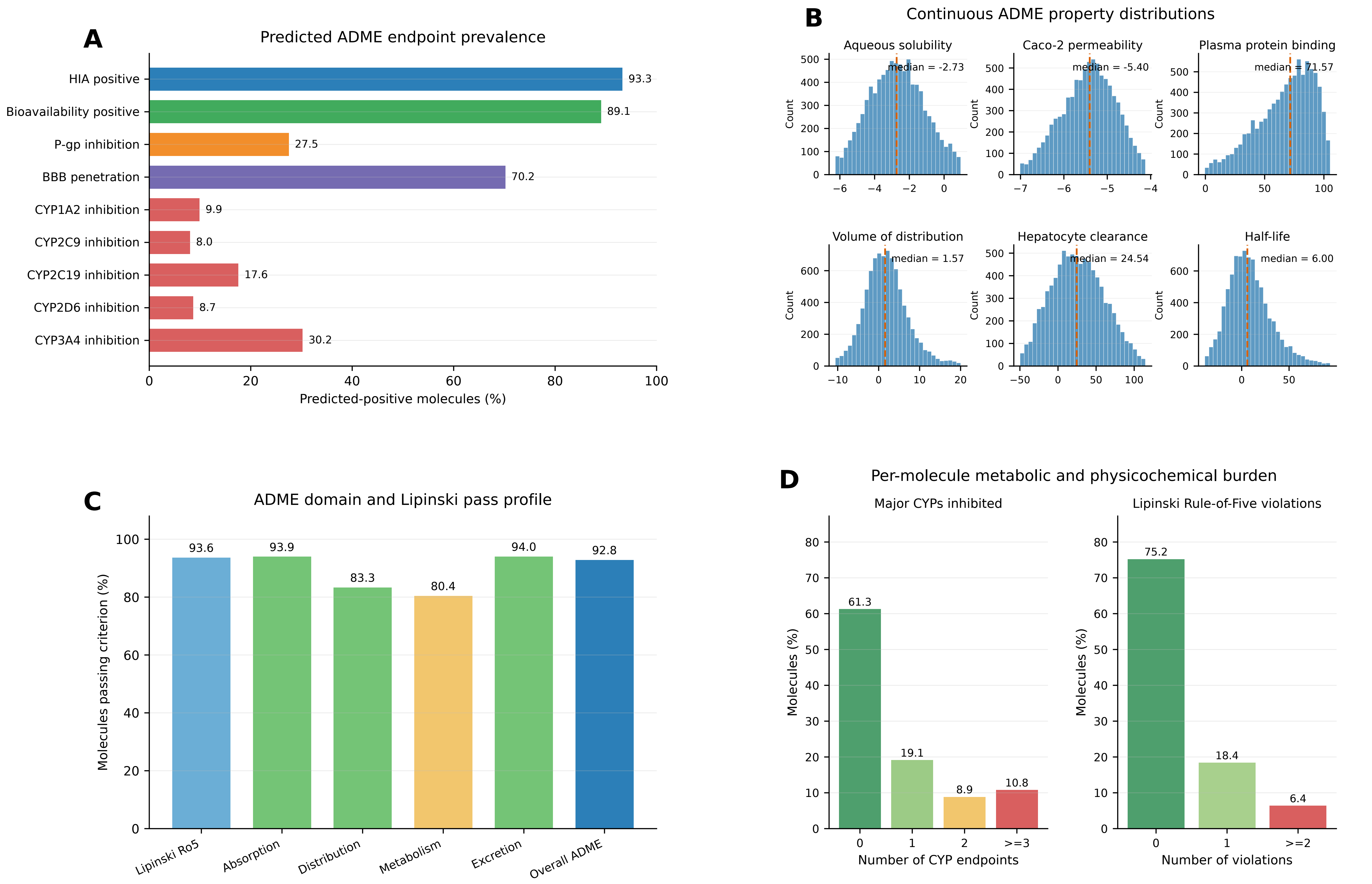


n = 8,700 molecule records (8,653 unique SMILES) across 97 targets; predicted-positive threshold = 0.5.

Distribution and excretion pass bands are comparative pooled 5th-95th percentile ranges.

### Fig. S24. ADMET-AI ADME profiling of FLOWR-generated molecules

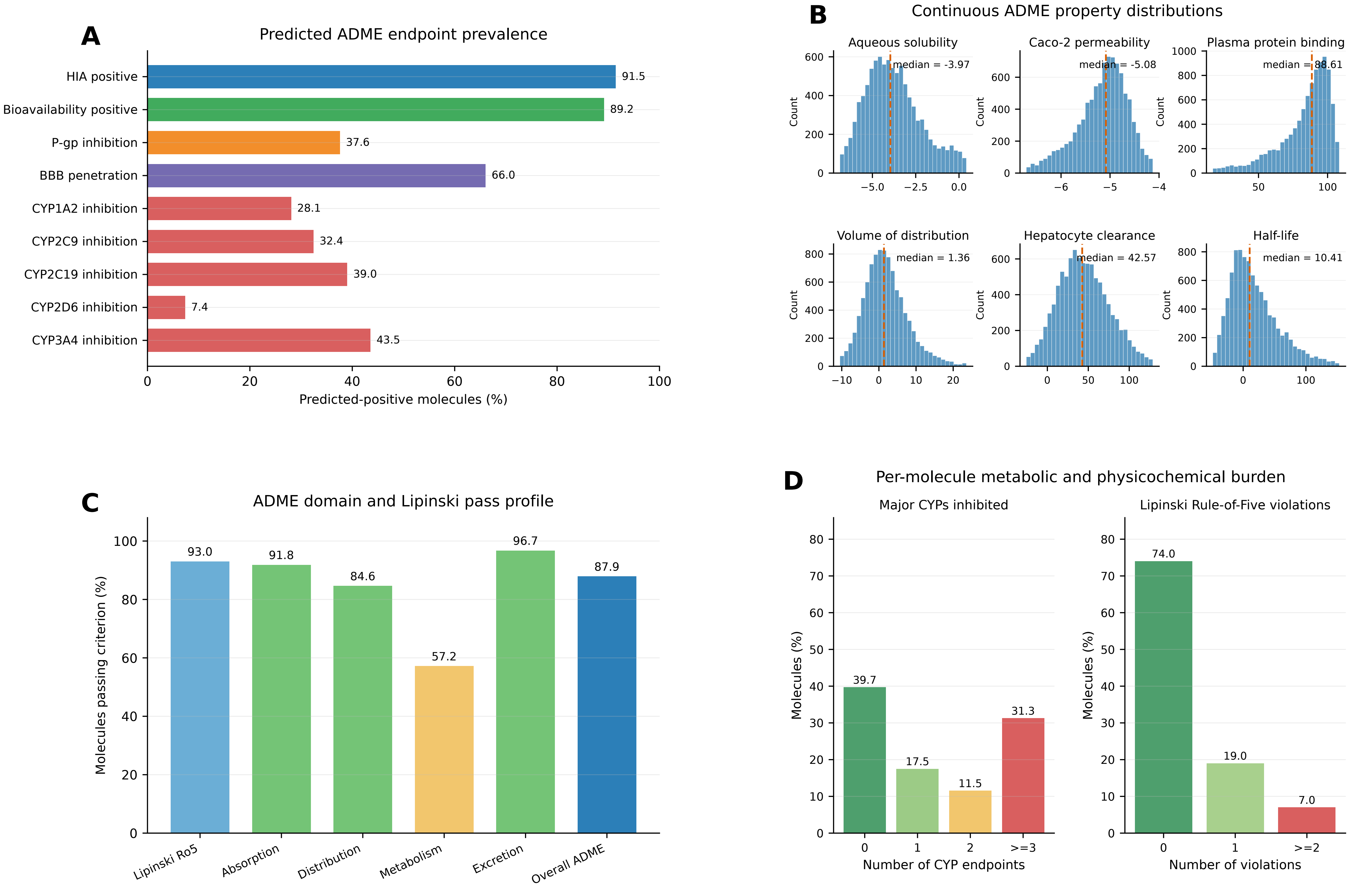


n = 9,695 molecule records (9,674 unique SMILES) across 97 targets; predicted-positive threshold = 0.5.

Distribution and excretion pass bands are comparative pooled 5th-95th percentile ranges.

**Fig. S25. ADMET-AI toxicity profiling of PFM-generated molecules**

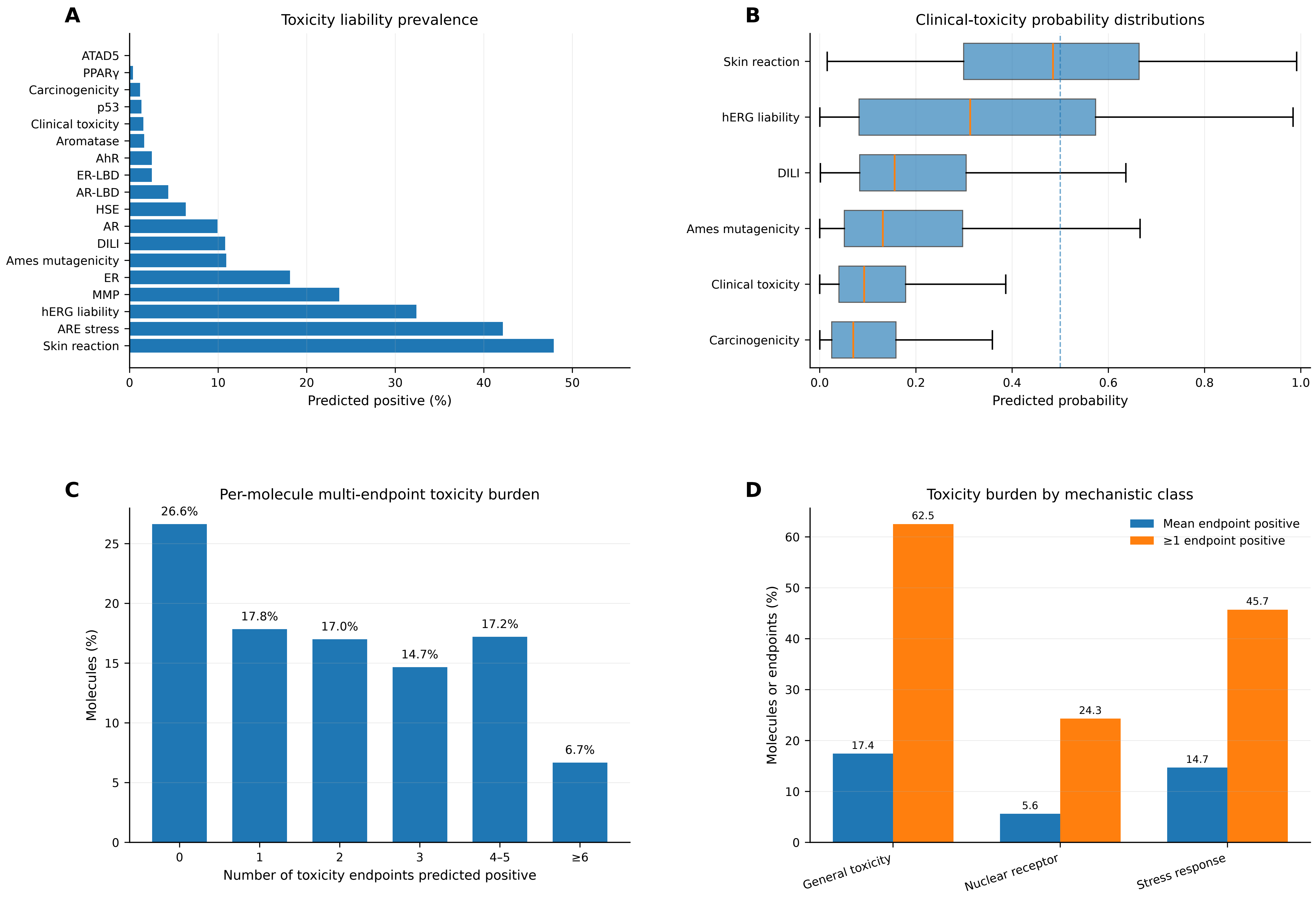


n = 16,430 molecules across 173 targets; predicted-positive threshold = 0.5.

### Fig. S26. ADMET-AI toxicity profiling of MolJO-generated molecules

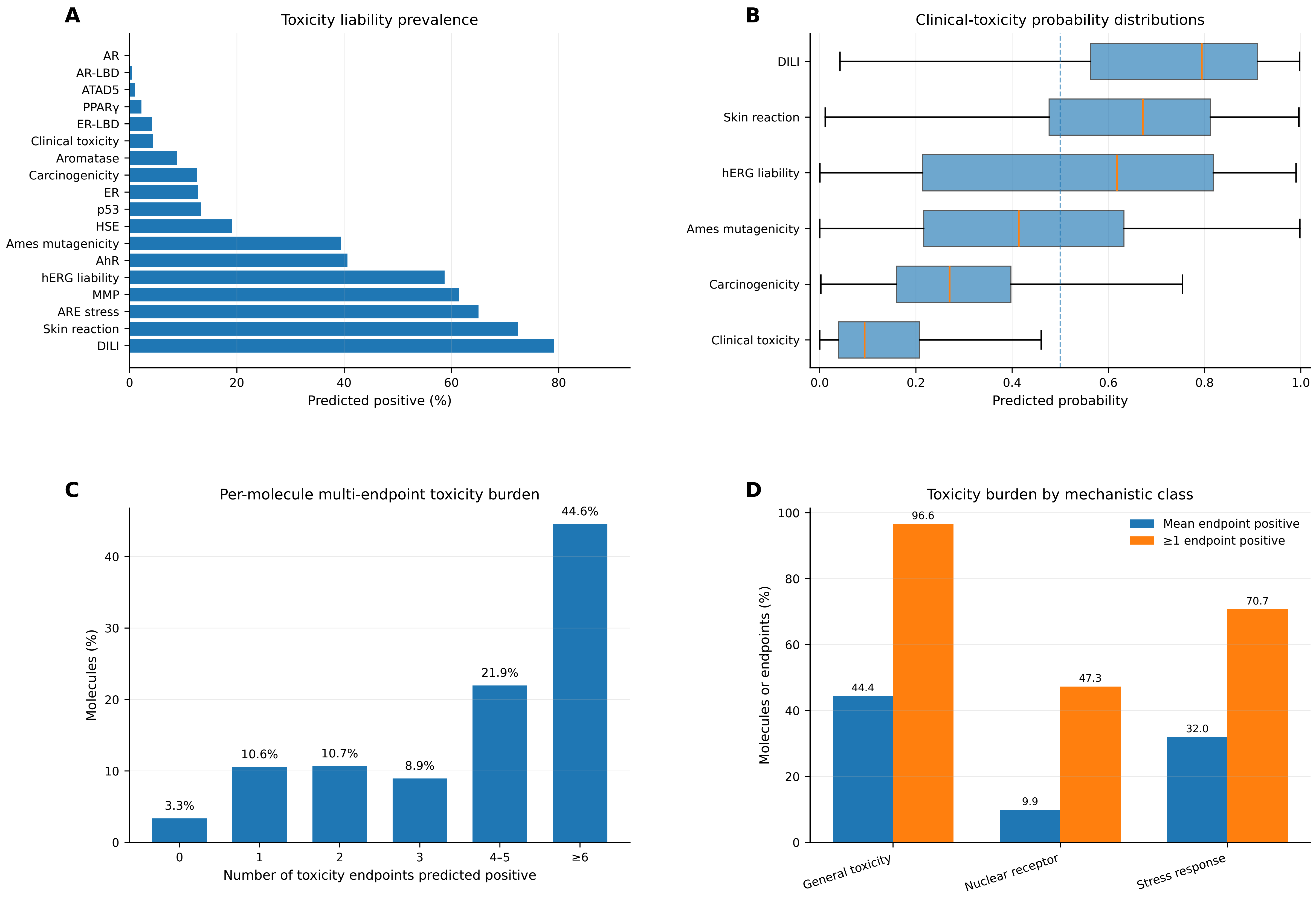


n = 17,383 molecules across 175 targets; predicted-positive threshold = 0.5.

#### Fig. S27. ADMET-AI toxicity profiling of DrugFlow-generated molecules

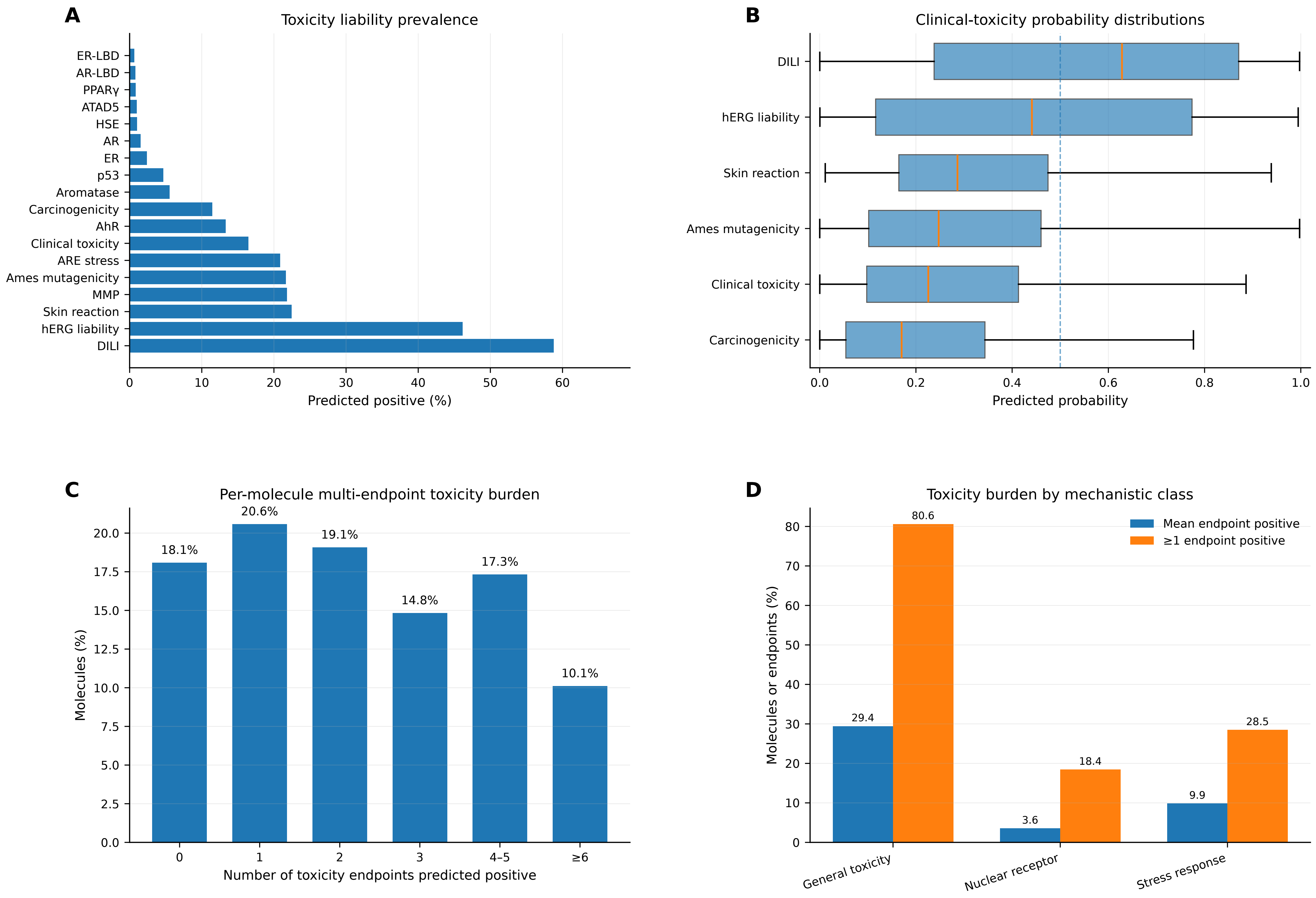


n = 17,601 molecules across 163 targets; predicted-positive threshold = 0.5.

### Fig. S28. ADMET-AI toxicity profiling of FlowMol-generated molecules

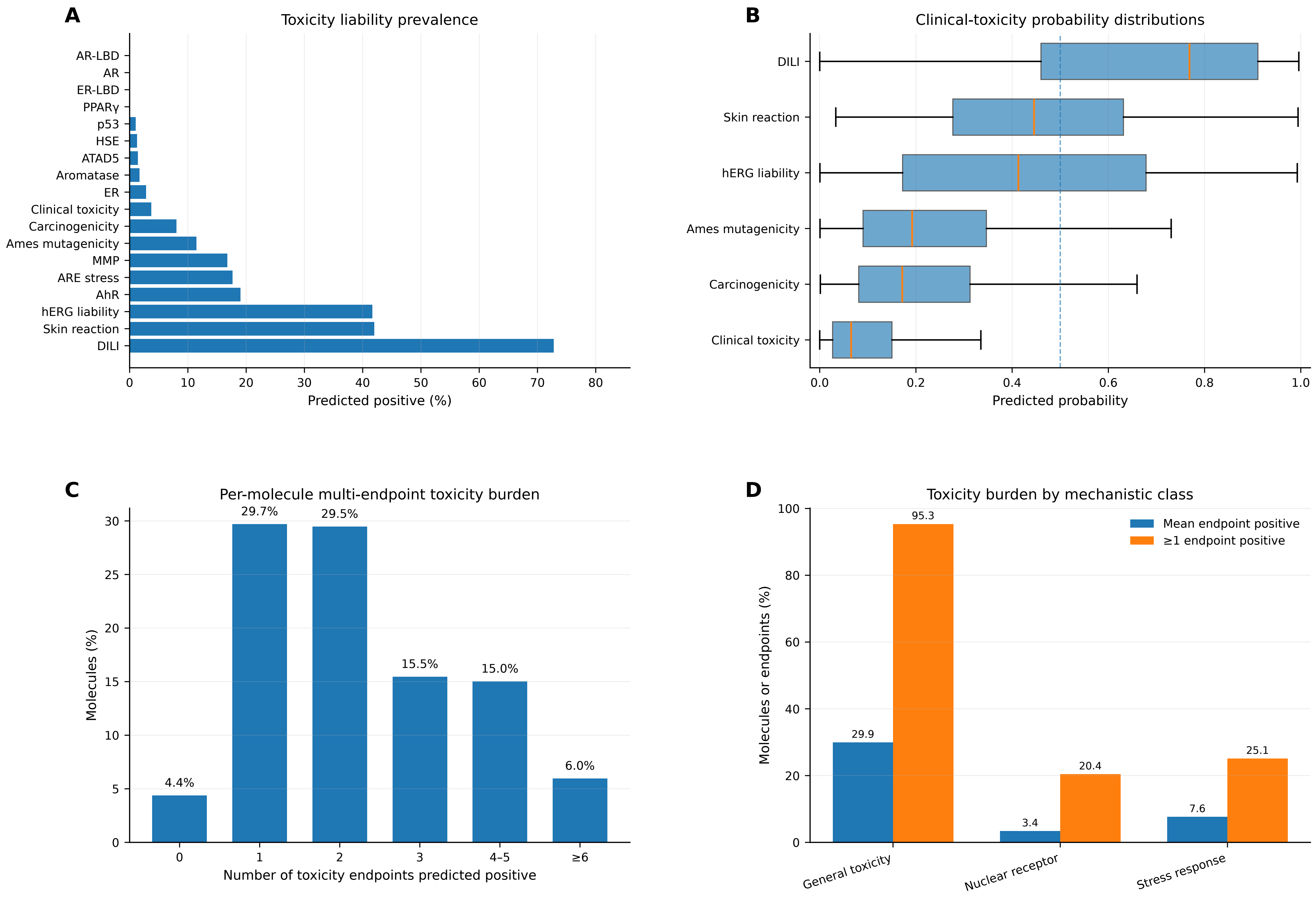


n = 17,498 molecules across 175 targets; predicted-positive threshold = 0.5.

### Fig. S29. ADMET-AI toxicity profiling of Pocket2Mol-generated molecules

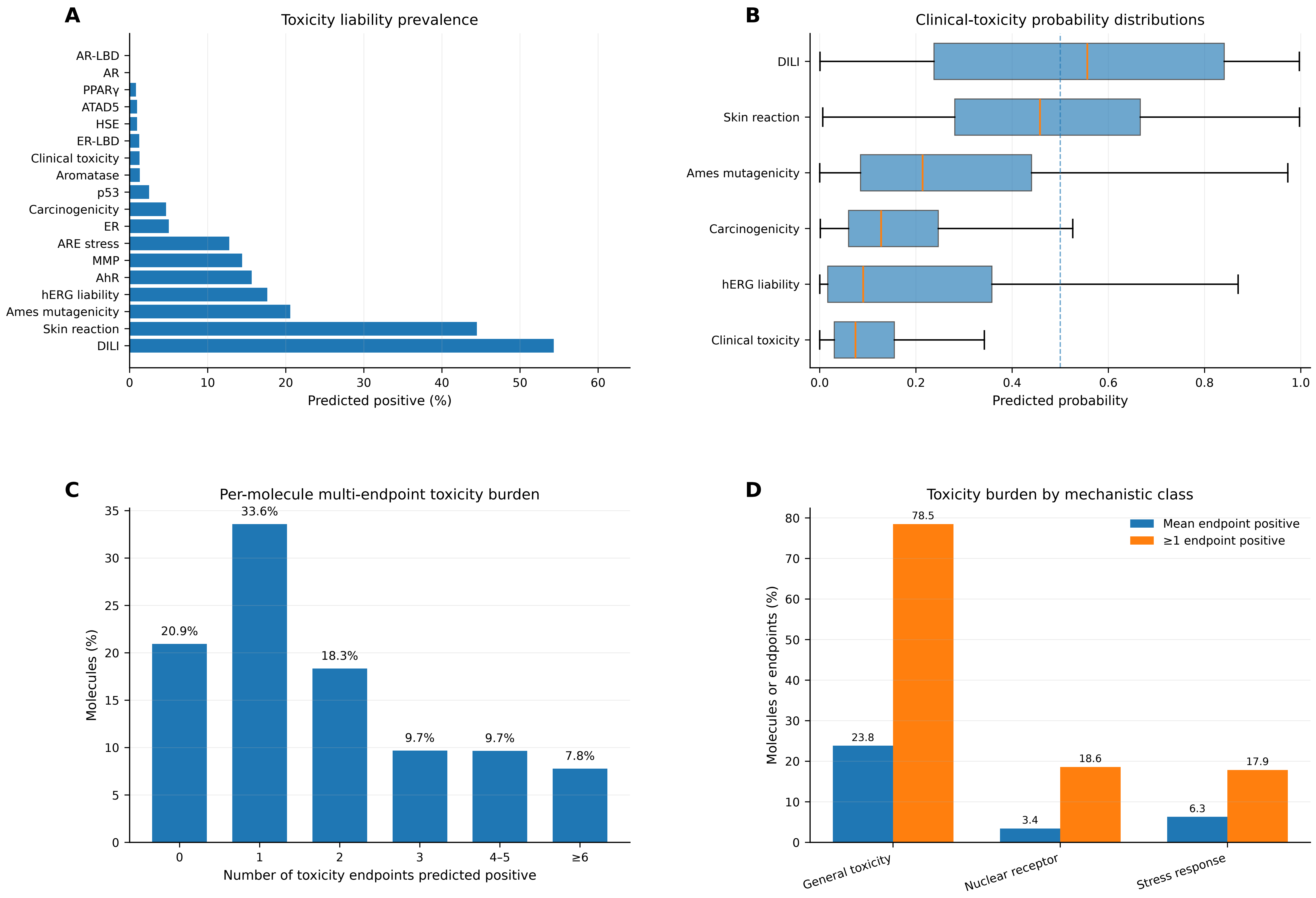


n = 18,682 unique molecules across 175 targets; predicted-positive threshold = 0.5.

### Fig. S30. ADMET-AI toxicity profiling of ShEPhERD-generated molecules

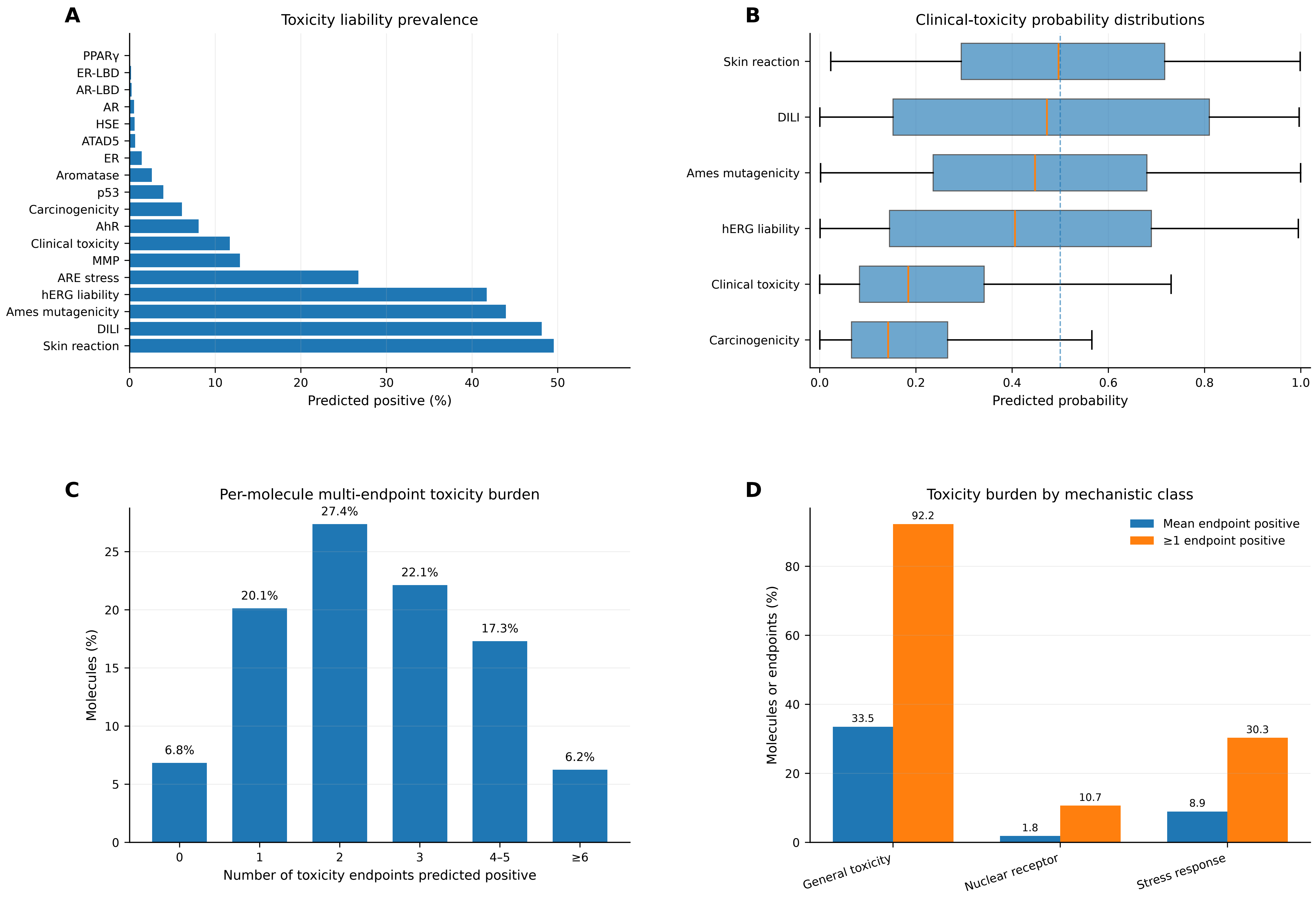


n = 16,307 molecules across 165 targets; predicted-positive threshold = 0.5.

### Fig. S31. ADMET-AI toxicity profiling of REINVENT4-generated molecules

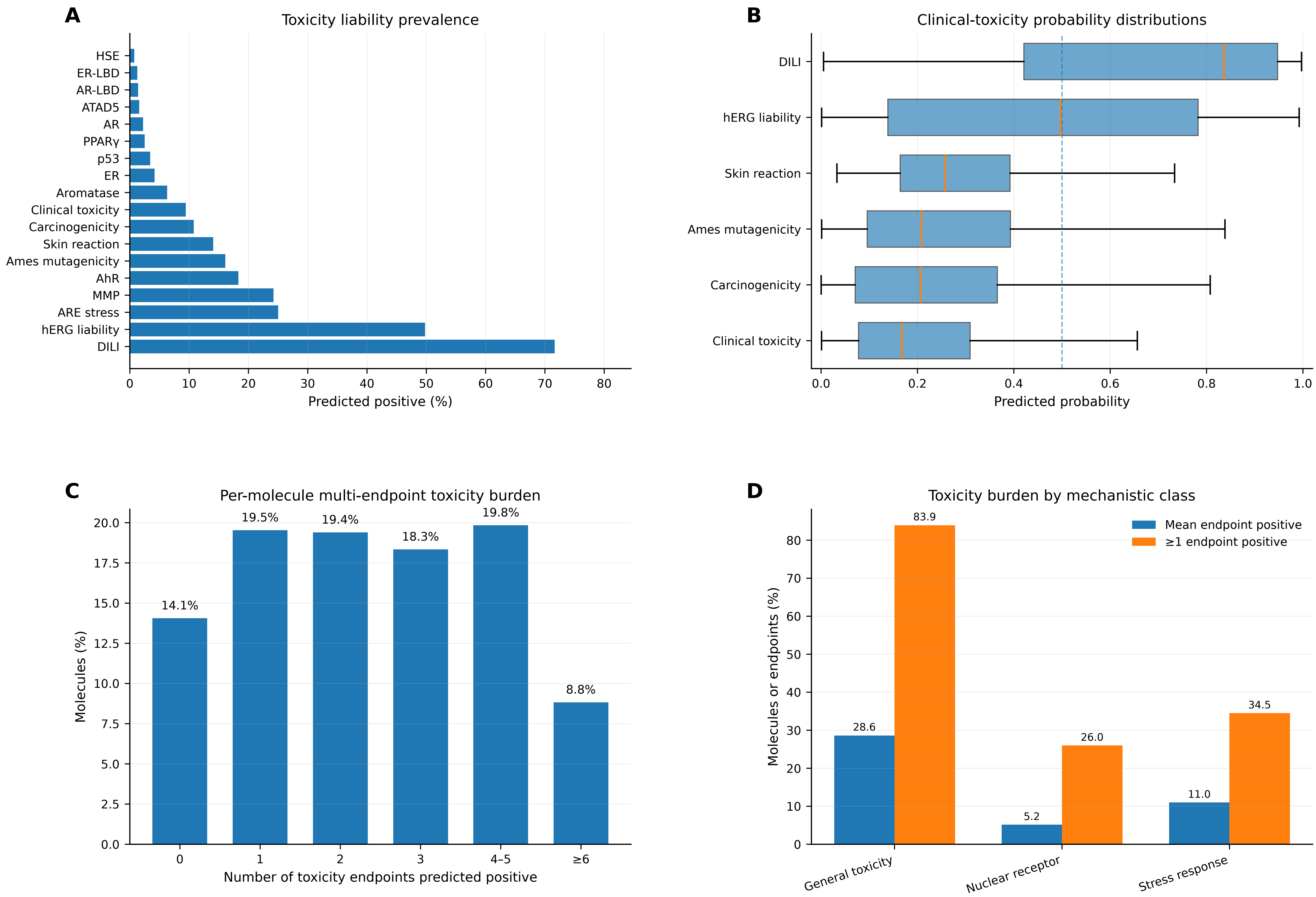


n = 11,137 molecules across 114 targets; predicted-positive threshold = 0.5.

### Fig. S32. ADMET-AI toxicity profiling of ReaSyn-generated molecules

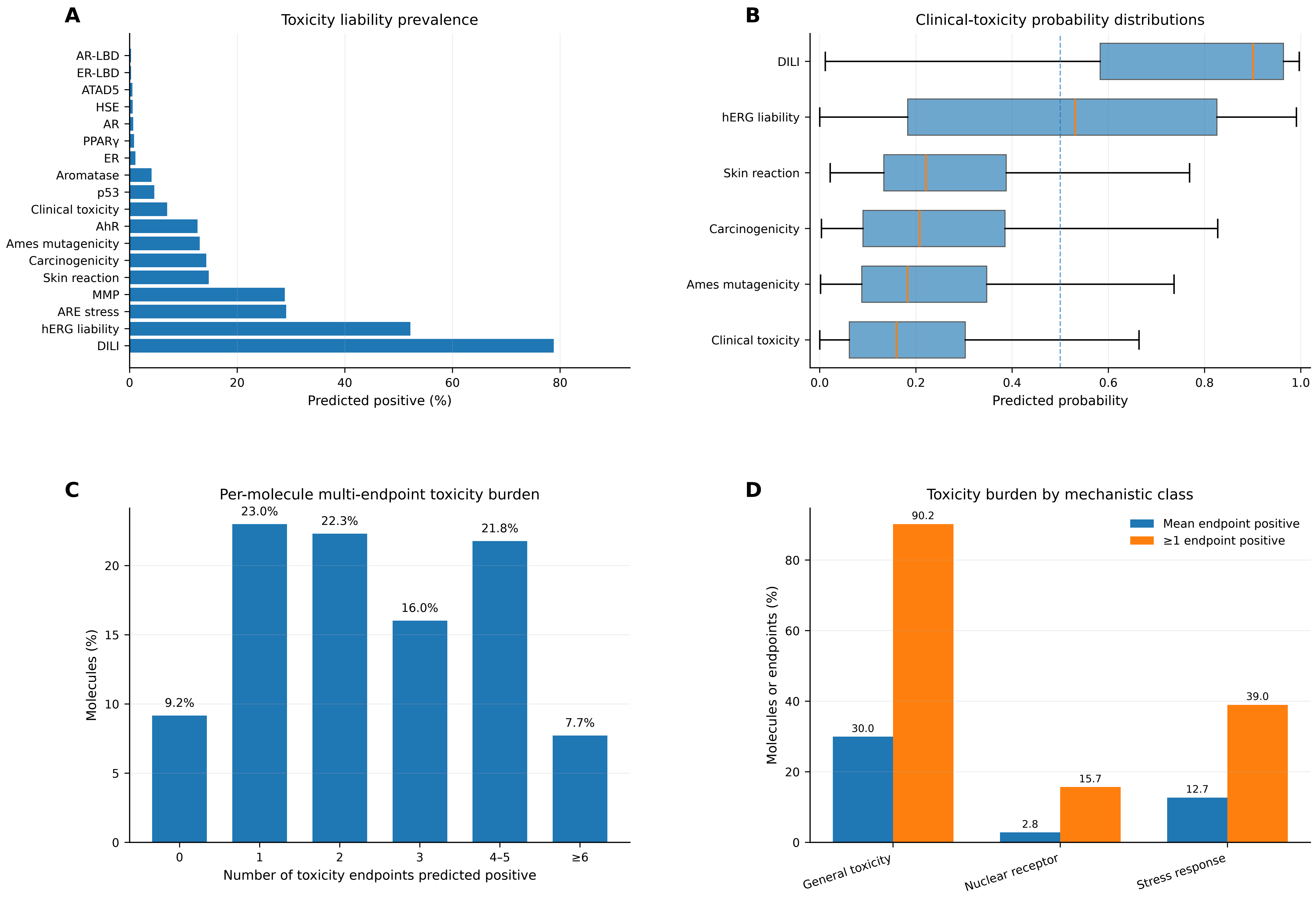


n = 12,094 molecules across 151 targets; predicted-positive threshold = 0.5.

### Fig. S33. ADMET-AI toxicity profiling of SynFormer-generated molecules

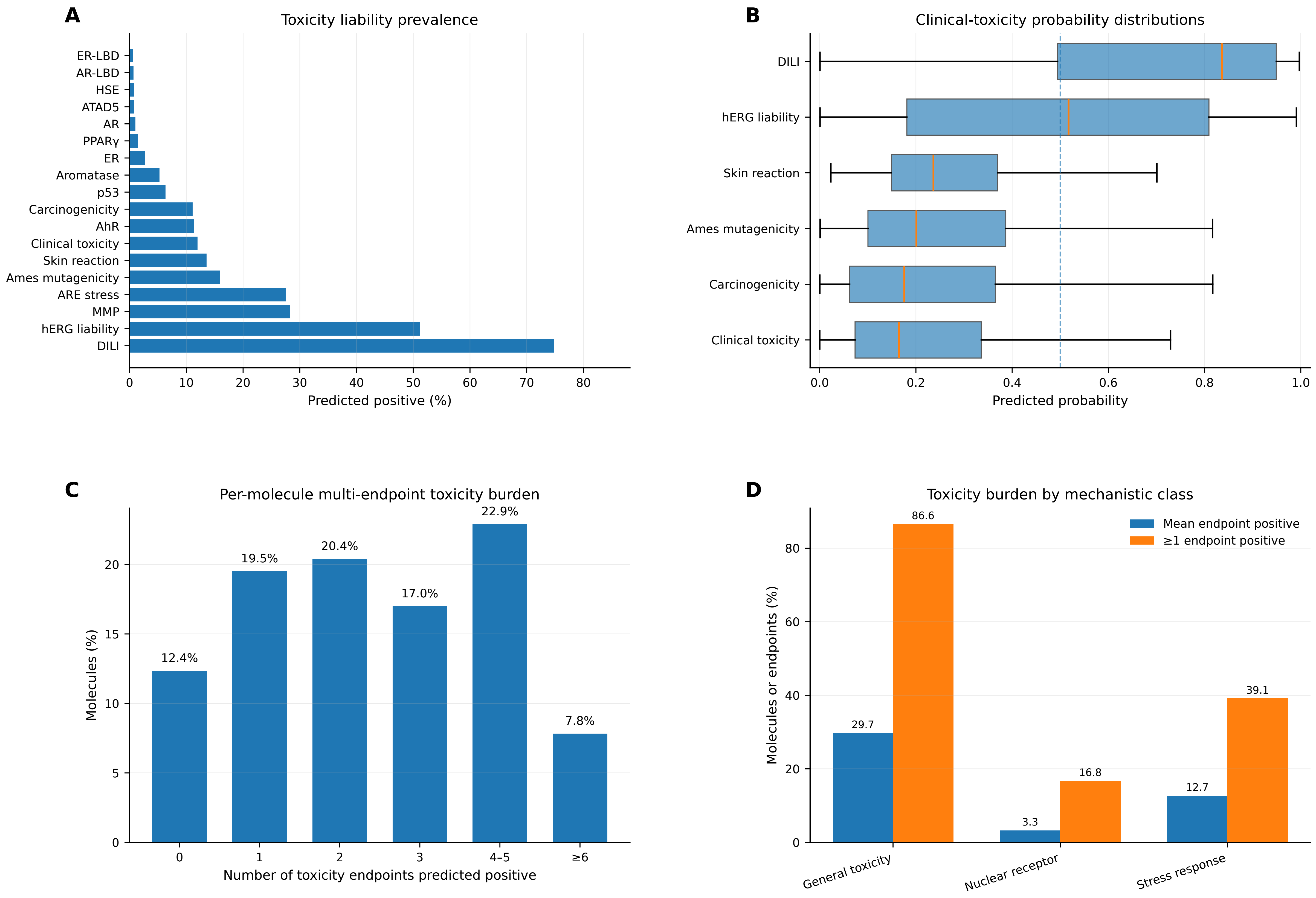


n = 14,603 molecules across 164 targets; predicted-positive threshold = 0.5.

### Fig. S34. ADMET-AI toxicity profiling of PrexSyn-generated molecules

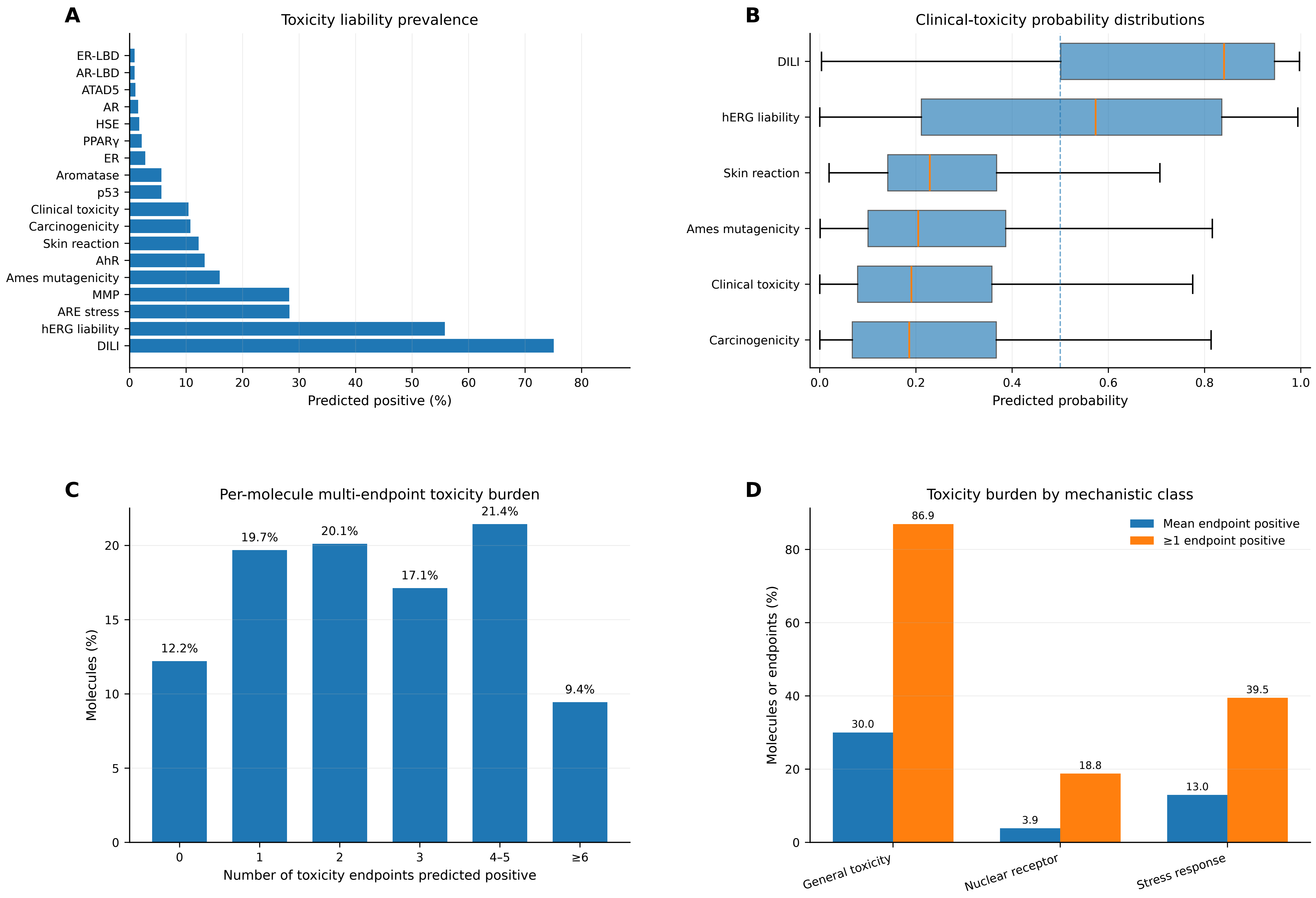


n = 13,804 molecules across 163 targets; predicted-positive threshold = 0.5.

### Fig. S35. ADMET-AI toxicity profiling of TargetDiff-generated molecules

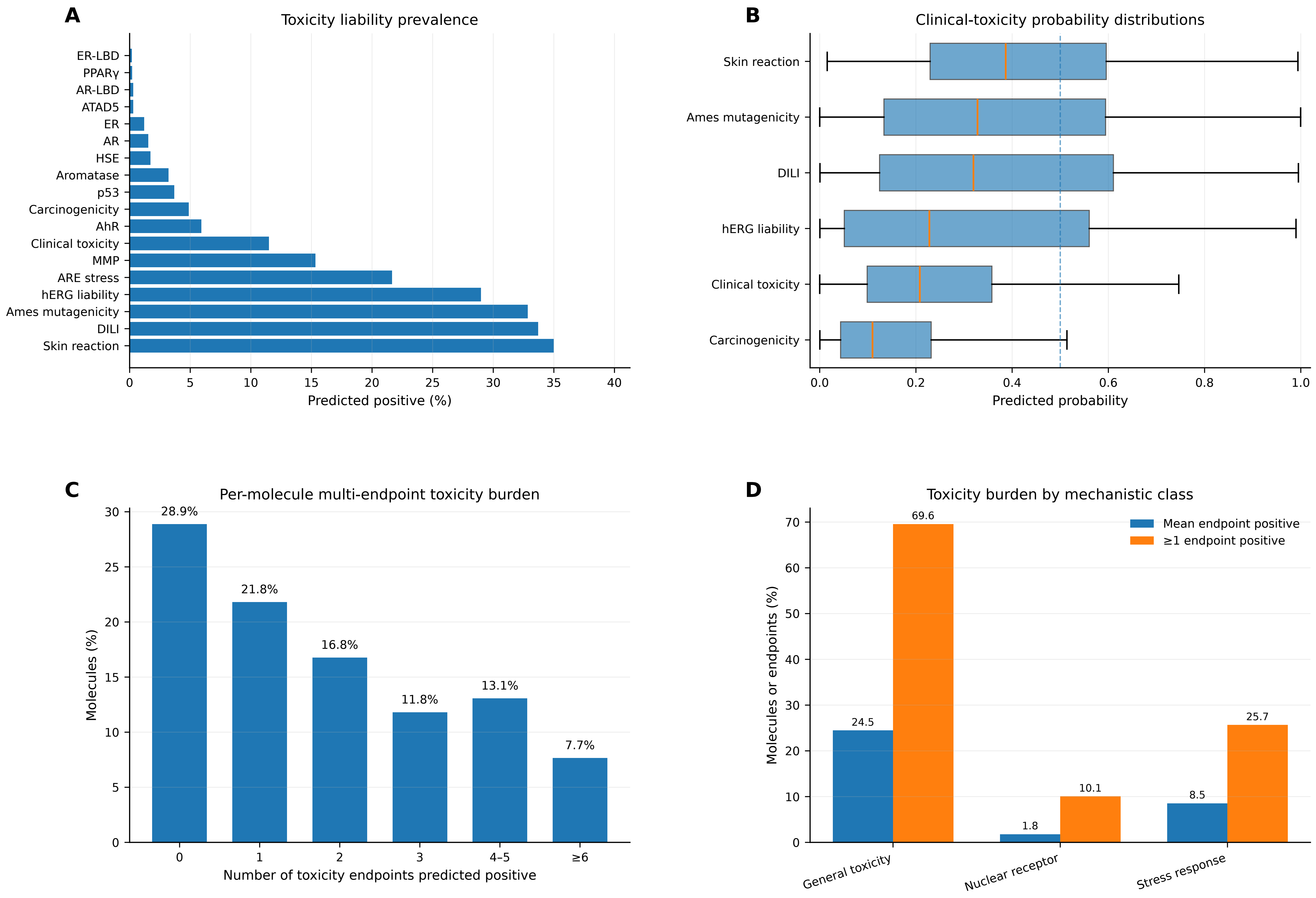


n = 14,779 molecules across 174 targets; predicted-positive threshold = 0.5.

### Fig. S36. ADMET-AI toxicity profiling of FLOWR-generated molecules

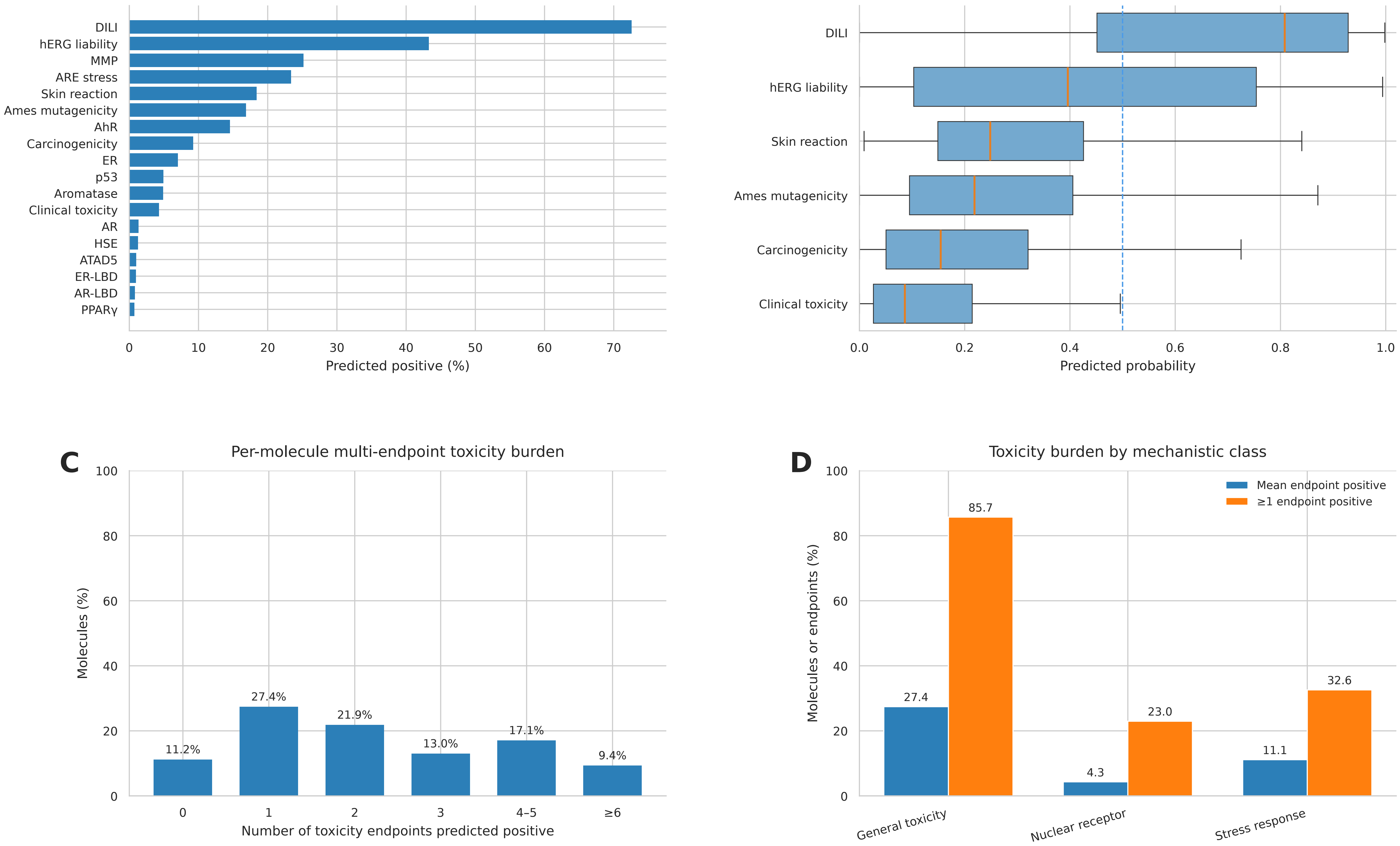


n = 16,285 molecules across 164 targets; predicted-positive threshold = 0.5.

### Fig. S37. State-aware functional classifier (SAFC) pipeline

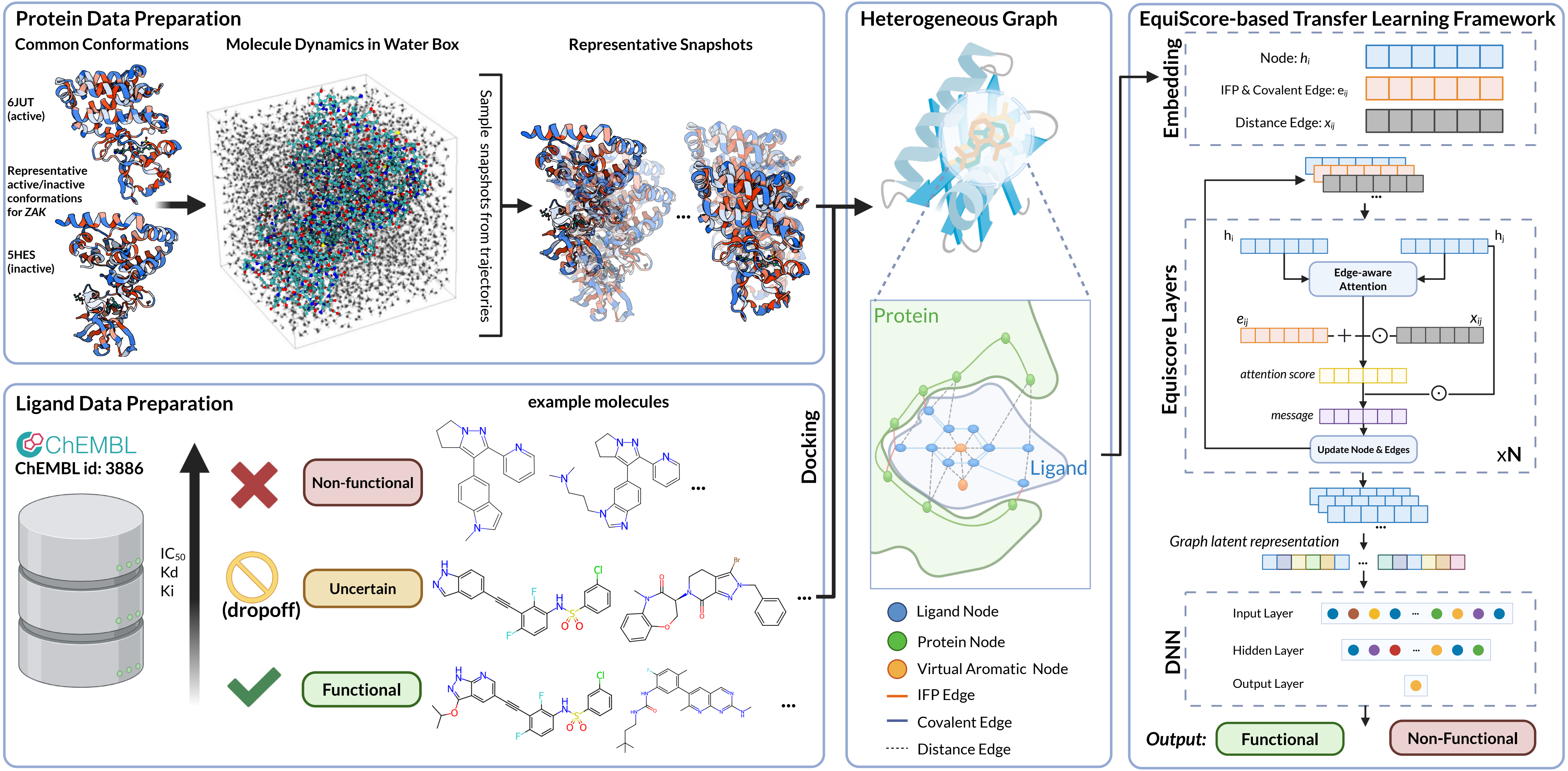
